# Randomised Badger Culling Trial lacks evidence for proactive badger culling effect on tuberculosis in cattle: comment on Mills et al. 2024, Parts I & II

**DOI:** 10.1101/2024.09.18.613634

**Authors:** PR Torgerson, S Hartnack, P Rasmussen, F Lewis, P O’Donnell, TES Langton

## Abstract

Re-evaluation of statistical analysis of the Randomised Badger Culling Trial (RBCT) by Torgerson et al. 2024 was rebutted by Mills et al. 2024 Parts I and II. The rebuttal defended the use of count rather than rate when considering bovine tuberculosis herd incidence. The defence makes biologically implausible use of Information Criterion for appraisal diagnostics; overfits data; and has erroneous Bayesian analyses. It favours ‘goodness of fit’ over ‘predictive power’, for a small data set, when the study was to inform application. Importantly, for ‘total’ bTB breakdown: (‘confirmed’ (OTF-W) +’unconfirmed’ (OTF-S)), where modern interpretation of the main diagnostic bTB test better indicates the incidence rate of herd breakdown, there is no effect in cull and neighbouring areas, across all statistical models. The RBCT was a small, single experiment with unknown factors. With respect to the paradigm of reproducibility and the FAIR principles, the original RBCT analysis and recent efforts to support it are wholly unconvincing. The 2006 conclusion of the RBCT that “*badger culling is unlikely to contribute positively to the control of cattle TB in Britain*” is supported, but the route to such a position is revised in the light of modern veterinary understanding and statistical reappraisal.

## 1. Introduction

The Randomised Badger Culling Trial (RBCT) (1,2) was an experiment with ten intervention-control comparison areas, designed to investigate if culling of European badger *Meles meles*, by trapping and shooting across wide areas in England (Proactive culling) could have an effect on the incidence of tuberculosis (bTB) in cattle herds. The RBCT proactive cull analyses were first published in Nature in 2006 (2) [“the 2006 paper”] and indicated that such an effect existed. Two separate re-evaluations of data from the 2006 paper have produced one view of badger culling having no effect on bTB herd incidence (rates)(3) and two where an effect is said to be supported(4,5).

Much of the debate surrounds the use of Bayesian information criterion (BIC) and small sample size Akaike information criterion (AICc) criteria. Mills *et al*. (4,5) are reliant on the use BIC in frequentist model diagnostics stating “*We recall here that unlike AIC (and AICc) which measure predictive accuracy, BIC measures goodness-of-fit*”. Consequently Mills *et al*. concluded that the BIC approach selected the model with the best *“goodness of fit”* and therefore the 2006 paper findings (2) were “robust”. The optimal model proposed by Torgerson *et al*. (3) performed far better by AICc criteria (ie “*predictive accuracy*”). There is, therefore, some agreement between the two analyses. Torgerson *et al*. (3) who published first, stated that the preferred model, first reported in 2006 (3), is now “*useful in reference only to its initial data set, which would include the specific idiosyncrasies of the data within each triplet, but it would have little predictive power* “. Predicting the outcome of widespread badger culling was the aim of the RBCT and, therefore, model selection from the perspective of predictive power more closely aligns with the RBCT’s applied interpretation.

Concerns over the position of Mills *et al*. (4,5) include a failure to address the biological implausibility of the methods of analysing incidence rate and the importance of incorporating diagnostic error in the analysis consistent with sound epidemiological practice. When analysis is adjusted for diagnostic error: i.e. models that encompass total herd breakdowns, which included unconfirmed breakdowns (OTF-S), there is no evidence of an effect of badger culling on bovine tuberculosis. This finding is consistent across all statistical models utilized in the analysis of the RBCT data and all four analyses(2–5) agree that using all test reactors show no effect was present. This contrasts with the models that examine only confirmed breakdowns (OTF-W). Taking all these issues together, it is concluded that the RBCT failed to provide evidence that culling of badgers had any significant effect on the incidence of bovine tuberculosis in cattle herds. Further, the Bayesian approach as presented by Mills *et al*. (4,5) has too many errors in the model code, and reported effect sizes, to be functional.

Recent scrutiny of the 2006 paper data ‘confirmed’ breakdowns only data suggests that its findings are unsound (3). Mills *et al*. (4,5) who included two of the authors of the original RBCT study, present a detailed analysis of the RBCT data from both within cull areas and their neighbouring (surrounding) areas and conclude that the RBCT finding are “robust”. The two publications of Mills *et al*. (4,5) draw almost entirely on the peer reviewed publication (3) which provided contrary evidence. The detailed and extensive use of statistical appraisal and diagnostic techniques used by Mills *et al*. (4,5) were examined in the present study to assess the strength of their claims, using the same methods of model appraisal and diagnostics and to check initial, more obvious concerns with the Bayesian analysis which might change or invalidate their conclusions.

Further information on study justification and context is provided in the supplementary information 1.

## 2. Effect of proactive badger culling on incidence of bovine tuberculosis in cattle

The original statistical model that analysed RBCT data was a log-linear Poisson regression, with number of incident cases as the dependent variable, and Treatment (culled or not culled), log of historical incident cases, Triplet (experimental pairs with a culled group and not culled group) and log of the number of herds as explanatory variables. Treatment effect was highly significant, concluding that culling badgers reduced the number of bTB herd incidents in cattle (2). *Torgerson et al*. (3) found that this conclusion was not reliable. However, Mills *et al*. (4,5) used several appraisal methods to imply the statistical model used to analyse RBCT data resulted in “robust” results. Table 1 presents finding and appraisal analytics for the model used to analyse the RBCT data in (2) and three other models. Model 1 is used to defend the original conclusions first published in the 2006 paper. These four models are presented to clearly demonstrate the statistical issues at hand. Further details including statistical code are available in the supplementary information 2 & 3 and data files 4.

**Table 1.** A selection of the frequentist models analysed in Mills et al 2024(4) and Torgerson et al.2024(3). This compares the original model from(2) (and its equivalent in generalized Poisson form). Mills *et al*. 2024a claim to be a robust interpretation of the RBCT results and “strongly” supports an effect of proactive badger culling upon bTB herd incidence. The present study compares this to those models with the lowest AICc values which provide no support for an effect of culling.

| Models from Mills et al. | Equivalent in Torgerson et al. | Structure | Estimated effect of culling (95% CI) | BIC | AICc | LOOCV RMSE | Number of estimated parameters | Parameter for exposure (95% CI) | Comments |
| --- | --- | --- | --- | --- | --- | --- | --- | --- | --- |
| 1 | 1 | original Poisson GLM | −18.7% (−29.5%, −6.2%) | 155.2 | 203.0 | 9.97 | 13 | 0.05 (−0.44, 0.53) | Very high AICc indicates poor predictive powers. Exposure parameter not significantly different from zero indicating implausibility. |
| 3 | 3 | original GLM in generalized Poisson form | −18.7% (−24.6%, −12.3%) | 147.1 | 217.2 | 9.95 | 13+1* | 0.04 (−0.21, 0.30) | Very high AICc indicates poor predictive powers. Exposure parameter not significantly different from zero indicating implausibility. |
| 4 | 3a | generalized Poisson (without any culling effect) | None | 161.7 | 160.4 | 10.93 | 3+1* | 0.66 (0.33, 0.98) | Low AICc, plausible parameter for exposure parameter |
| 8 | 4d | generalized Poisson with herd-years-at-risk covariate without any culling effect | None | 155.5 | 154.2 | 8.81 | 3+1* | 0.51 (0.32, 0.70) | Lowest AICc of all the Poisson family models, indicating the best predictive power. Lowest LOOCV RSME indicating generalizability. Exposure parameter is significantly higher than zero, indicating plausibility, although as it is somewhat less than 1 it might indicate the offset model would be more appropriate. |
| Null |  | Generalised Poisson model with no predictors | NA | 178.1 | 176.8 | 17.29 | 1+1* | NA | Null intercept only model (when correcting for overdispersion) has lower AICc than model 1 and 3. |

\*Additional parameter in generalized Poisson models to model overdispersion

## 2.1. Incidence rates and counts

The unit of comparison for the RBCT data between culled and control areas is fundamental to the interpretation and robustness of the results in Nature (2) and the restatement in Mills *et al*. (4,5). Rates are the number of herd breakdowns per unit herd per unit time. That is they can be used to assess any difference in herd incidence by correctly adjusting for sample size and time of exposure. Counts are the unadjusted number of herd breakdowns without any reference to numbers of herds or time of observation in the sample. Although rates are specified in the 2006 paper (2), it was counts that were statistically modelled. Mills *et al*. (4) claimed this model (Model 1) was nevertheless “robust” as it passed many (but not all) of the statistical appraisal methods and *post hoc* analyses.

Model 1 used the number of herds as an explanatory variable rather than as a denominator. It can be simply converted to a rate by use of the variable as an offset, which as Mills *et al*. (4) correctly explain: “*Another possible option for a Poisson regression model is the usage of an offset variable which enables modelling the count variable (here confirmed herd breakdowns) as a rate, and the usage of an offset variable means that the corresponding regression coefficient is constrained to be 1*.” The algebraic derivation of this for Poisson regression models has long been known, but is restated in Torgerson *et al*. (3). However in Model 1, with the unconstrained variable of log(number of herds), the parameter value is 0.04 which is very close to, and not significantly different, from zero. The interpretation of this is that the number of herd breakdowns does not vary with the number of herds (that is the count remains the same regardless of sample size). This approach is biologically implausible. Mills *et al*. (4), states: *“Alternatively, assuming an offset variable not be supported by evidence (i*.*e. the number of events may increase non-proportionally with the population at risk) one could use an unconstrained regression coefficient and hence, instead of assuming the slope for the variable is exactly 1, the slope parameter is estimated*.*”* This is also discussed in detail in Torgerson *et al*. (3). However, in the purported “robust” model there is no *“non proportional increase in the counts with increase sample size”*. Indeed there is no increase at all. Thus the model must be misspecified and misleading even if claimed statistical checks suggest otherwise. This exemplifies the tension between plain statistical approach and the experience of the epidemiologist. This substantial problem can be managed in two ways. Firstly by having an offset variable rather than an unconstrained variable. When this is done (Model 2) and corrected for overdispersion, the treatment effect of culling becomes non-significant. Secondly by removing the 9 free parameters of Triplet. There is strong evidence of collinearity of the variable triplet and removal from the model leaves to a dramatic fall in the AICc. When triplet is removed, but the parameter coefficient is unrestrained, the value of new coefficient of the exposure variable becomes 0.64. If corrected for overdispersion (i.e. the generalized Poisson model), the upper confidence interval is close to 1. This is at least biologically plausible as there is an increase in counts with sample size. Removing the variable of Triplet also results in no significant effect of culling when the model is corrected for overdispersion. Similar issues can be identified in the analysis of post-trial period (See supplementary material 2 & 3).

### 2.2. Model appraisal and diagnostics

Mills *et al*. (4,5) make extensive use of model diagnostics to suggest that the 2006 (2) results were “robust”. In particular they use small sample size Akaike information criterion (AICc), Bayesian information criterion (BIC), leave-one-out cross validation (LOOCV) and posterior predictive checks (PPC). In all cases the AICc of the optimal model published in Torgerson *et al*. (3) has substantially lower AICc than Model 1 defended by Mills *et al*. (4) as “robust”. In addition the LOOCV values are better (Table 1). This is accepted, however Mills *et al*. dismissed the use of AICc (a standard diagnostic for statistical performance, especially when there is a small sample size) as being useful only as a *“predictive diagnostic”* but would rather use the BIC as it gives *“better performance for goodness of fit*.*”* Leaving aside the point that a predictive model would be a better outcome for a trial of the type conducted, which is used to inform wild animal culling policy, the BIC of the model reported in the 2006 paper (155.24) (Table 1, Model 1) is only marginally better than the optimal model reported by Torgerson *et al*. (3) (155.52) (Table 1, Model 8). The difference is so small that it can be dismissed as useful for model selection. As can be seen in Table 1, Model 8 in terms of AICc is far superior compared to Model 1. This suggests, by Mills et al.(4) own arguments, that it has far better predictive powers. The LOOCV values also perform better with the Model 8 (8.81), compared to Model 1 (9.97). Mills *et al*. (5) state that LOOCV approximates to “model generalizability”. LOOCV also indicates that out of sample best predictive model does not contain the culling effect. Consequently any such “effect” is likely to be specific to the areas used, not the population the experiment was supposed to represent. Similar issues can be identified in the models from to the time to follow up models (detailed in supplementary material). In addition, for cross validation, LOOCV methods are preferable as bias is negligible (6).

Mills *et al*. (4) claim that a visual PPC indicates that posterior predictive distribution of the model originally reported in Donnelly *et al*. (2) (Model 1, Table 1) resembles the observed data. In contrast, they claim for the model with the lowest AICc (Model 8, Table 1) the PPC check implies potential model misfit due to systematic discrepancies between model-predicted data and confirmed incidence. The crucial issue with PPC is that it uses the data twice (7). The data are first used for estimating the model and then, for checking if the model fits the data. Essentially PPC checks how close the observations are to the model predictions, but the model parameters are dependent on the observations. LOOCV, in which model 8 performs better avoids this issue. Also Mills *et al*. relied on a visual PPC. However, this can appear markedly different between simulations. Further details are given in the the supplementary material (supplementary material 2).

### 2.3. Overfitting

Mills *et al*. have neglected the overfitting issue. This explains why, generally, Model 1 has poorer diagnostics (i.e. AICc, LOOCV) than the optimal Model 8. Model 1 has 13 free parameters with only 20 data points; while Model 8 has just 3 free parameters (2 predictors and 1 to model overdispersion), with generally better model diagnostics. There has been much debate in the statistical literature surrounding the number of predictors compared to the number of data points. Depending on the type of study and statistical model this has been suggested to be as little as 5 (8). The preferred model of Mills et al.(4) has 13 predictors for 20 data points. Although it satisfies the p<n rule (p is the number of predictors and n the number of data points) so avoids saturation (although not by much), there are clear issues of the potential for overfitting. It is also notable that Model 1 has a higher AICc than the null model or intercept only (generalised Poisson) model with no predictors. One suggested solution to the problem of overfitting is to combine dichotomous variables into a continuous variable (8). Model 8 effectively does this my combining all the 10 dichotomous variables of Triplet into a single continuous one of years at risk (further details in the supplementary material).

### 2.4. Quasipoisson model

Mills *et al*. make an issue of the quasipoisson model, highlighting that model comparisons cannot be made due to no likelihood structure. But the use of a generalized Poisson model deals with this issue for model comparison (3) and avoids this unnecessary distraction.

### 2.5. Modern interpretation of SICCT test reactors

In epidemiological studies, diagnostic tests are frequently used to categorize animals or groups of animals into diseased categories and non-diseased categories. This is almost always undertaken with the use of diagnostic test(s). Diagnostic tests rarely, if ever, have a diagnostic accuracy of 100%: that is both the sensitivity and specificity of the test is 100%. Modern epidemiological theory demands that analyses should, as much as possible, include the diagnostic error of the test in the analysis (9). Such adjustments are increasingly used, such as modelling the covid-19 pandemic (10). bTB should be no exception. The comparative intradermal skin test (SICCT) is the primary screening test for this purpose and was used in the RBCT trial. Recent work has shown that the specificity of the SICCT was close to 100% at standard interpretation (11), but with a low sensitivity. Mills *et al*. (4) have avoided detailed mention the key analytical issue of whether “unconfirmed” breakdowns should be included. However it is clear that the analysis was undertaken as it is documented in their supplementary material. ‘Unconfirmed’ breakdowns, as defined in the ISG report (1) and elsewhere are when one or more cattle in the herd test positive for the SICCT test but cannot be confirmed by finding lesions and/or a positive culture of *Mycobacterium bovis* at necropsy. Because the test specificity of the SICCT test is 100%, these animals would almost certainly have had bTB, and the inability to confirm it at *post mortem* was likely due to the poor sensitivity of necropsy. The later has an estimated sensitivity of 46% by routine meat inspection and 76% by detailed necropsy in the laboratory (11)

RBCT cattle that were SICCT positive, but had no visible lesions, at *post mortem* were likely in the earlier stages of infection. Therefore, in hindsight, they are essential in the analysis of an experiment that was designed to monitor any effect of an intervention on the rate of new infections in cattle herds. There was no evidence of an effect of badger culling on total number of herd breakdowns (confirmed and unconfirmed together) either in the 2006 (2) analysis or in the more extensive recent analysis (3). Mills *et al*. (4) neglect to mention that the model from 2006 fails to give any indication of a cull effect on total breakdowns, but rather preferred to be critical of the Torgerson *et al*. (3) re-evaluation analysis. For example, it is worth noting that the critiqued quasipoisson approach (supplement Mills et al.(4)) has 2 data points classified has highly influential. The same analysis of the model from 2006 indicates 4 influential data points, which is, surprisingly, not mentioned. However, arguments surrounding the model fitting for the total breakdowns can be put aside. All models give the same result implying that badger culling has no effect on total breakdowns regardless of modelling approach. This supports the approach to be the most robust consistent and strongest result and further implies that conclusion

In 2006 (2), Donnelly *et al*. stated *“Our finding that widespread culling of badgers has simultaneous positive and negative effects on the incidence of TB in cattle has important implications for the development of sustainable control policies. We would expect the overall reduction in cattle TB to be greatest for very large culling areas (with consequently lower perimeter:area ratios), although in absolute terms the costs, as well as the benefits, will be greatest for large areas. Detailed consideration is needed to determine whether culling on any particular scale would be economically and environmentally sustainable*.*”* Further in 2015, Donnelly and Woodroffe who are co-authors on the Mills et al. manuscripts, based on evidence from the RBCT also predicted that, “*better prospects for the control of cattle TB are offered by badger populations that are either reduced by more than 70% or left undisturbed — and potentially vaccinated*”(12). In other words they were using their results in a highly predictive manner to argue how a reduction in bTB would be achieved in practice. However, in 2024, Mills *et al*. (4) set aside their own analysis as inferior in “*predictive accuracy*.” The models with better predictive accuracy would suggest no overall reduction in bTB even over large culling areas.

## 3. Bayesian Analysis

Mills *et al*. (4,5) purportedly support their conclusions with a series of Bayesian models. They compare their models with similar Bayesian models proposed by Torgerson *et al*. (3). The equivalent models to those in Torgerson et al are defined in the supplementary material of Mills *et al*. (4,5). There are important issues that invalidate all of the Bayesian modelling presented by Mills *et al*. Firstly, there was a false claim that several models are a direct comparison to those of Torgerson *et al*. When examining the code and the effect size of culling it becomes apparent that the models are not the same. Other models claim to use an offset but due to coding errors the offset is omitted and the results of the analysis are without the offset. The issues of the Mills *et al*. Bayesian models are summarized in Table 2.

**Table 2:**
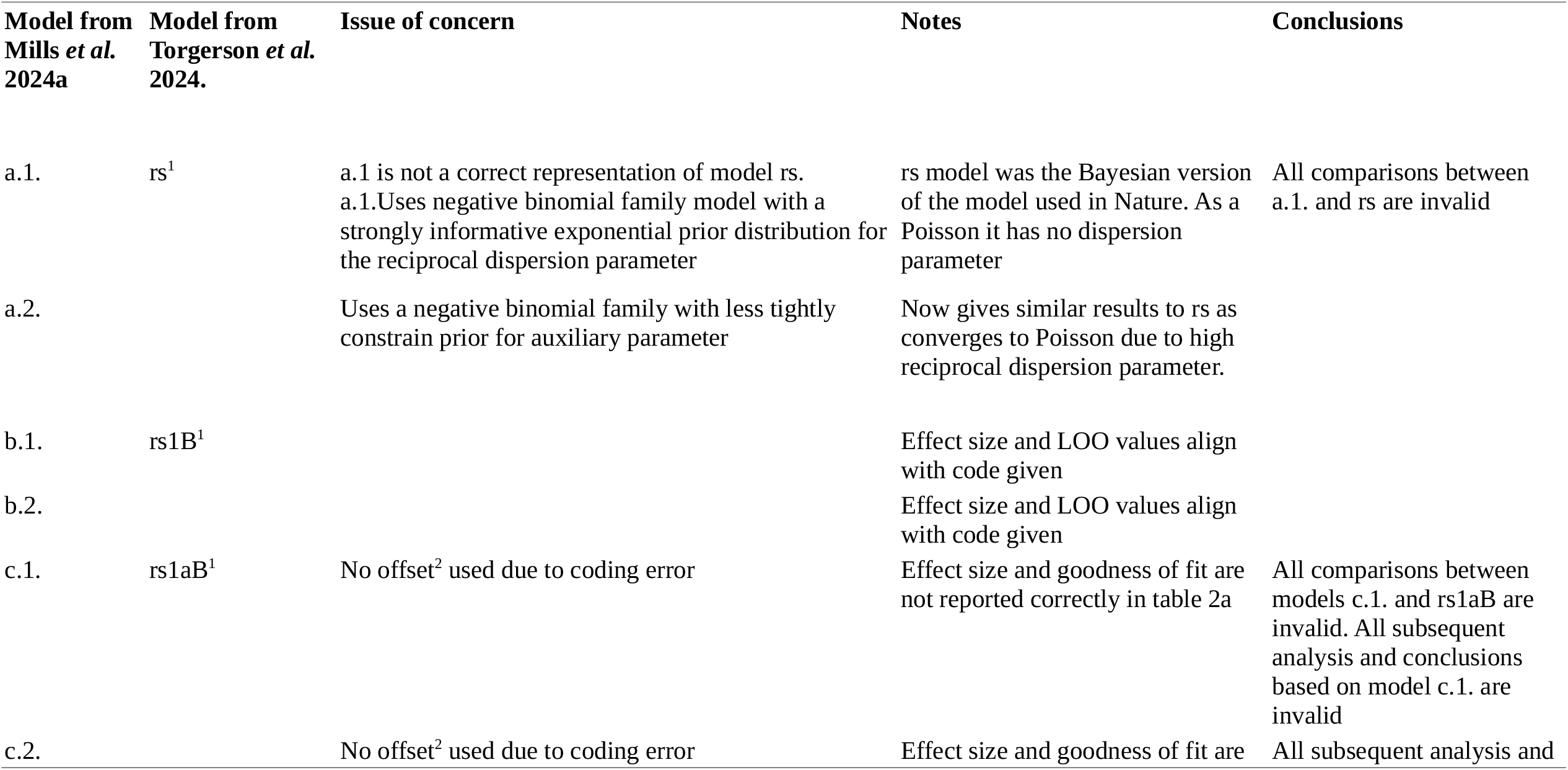
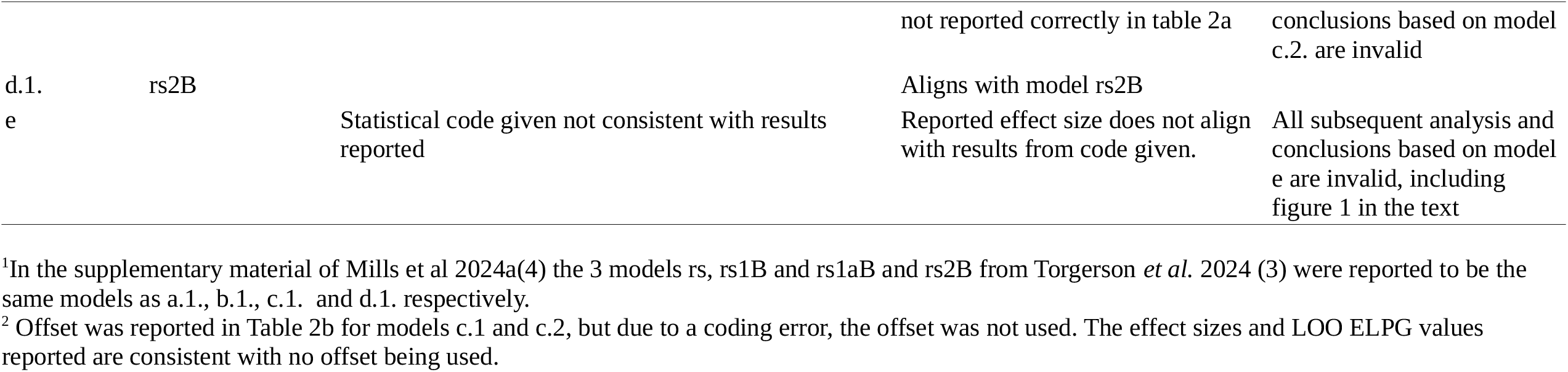
A selection of Bayesian models for confirmed bTB herd breakdowns from initial cull until 4 September 2005 within proactive culling/control areas of the RBCT experiment. The original, frequentist Poisson GLM used in Donnelly *et al*. in 2006(2) was re-specified in the Bayesian paradigm in the initial preprint, subsequently published paper by Torgerson *et al*.(3). However the Model rs was not specified by a negative binomial likelihood, although Mills et al. 2024a(4) reported it as Model a1 in error. Other error and inconsistencies between those reported in Mills *et al*. 2024a(4) and Torgerson *et al*. initial preprint and paper (see supplementary files 1 & 2) for the Bayesian analysis paradigm (tables 2a and 2b in Mills *et al*. 2024(4) are summarized here). In total of the 8 models specified in tables 2a and 2b of Mills et al. 2024a, 5 had errors. Hence the analysis cannot be relied upon for any of the Bayesian analyses in Mills *et al*. 2024a(4).

From the Bayesian paradigm it is important to note the use of model selection techniques, such as Bayes factors, which account for the complexity of the model (compare model rs2B with rs in Torgerson *et al*. (3), for example). Here, it can clearly be shown that the models without Triplet and Treatment as covariates are better supported by many orders of magnitude compared to those including these explanatory variables. Thus, the conclusion is that evidence points to no effect of culling on bTB herd incidence rates. Leaving aside the errors documented in Table 2, Mills et al made no comparisons or efforts at model selection. Mills *et al*. simply concluded that there was a greater probability of culling having an effect, with no comparison to suitable null models. An advantage of the use of Bayes factors is that it automatically penalizes the inclusion of too much model structure guarding against over fitting. As we have seen with the frequentist models, the modelling of 13 explanatory variables with just 20 data points is at high risk of over-fitting.

## 4. Statistical audit

One of the peer reviewers (13) of Mills *et al*. (4) requested further details of the statistical audit, which we now provide. Throughout the text Mills *et al*. repeatedly state that the statistical analyses of the RBCT were *“pre-defined and also independently audited by a statistical auditor”* as a further justification to defend the results of the RBCT. This is considered further in our supplementary material (supplementary file 1). Our analysis focussed mainly on Poisson regression models (and their over dispersed analogues). It is also interesting that, in the first report of the statistical auditor, it was stated that, *“to some extent, the number of triplets and the years of observation are interchangeable”*(14). This interchange is implemented when herd years at risk is used as an explanatory or offset variable and such an implementation fails to demonstrate an effect of culling (see models 6 and 4d in Table 1). Further details are in the supplementary file 1. Thus, in the RBCT this alternative analysis implied by the statistical auditor, if done, was not reported. In addition, the statistical auditor recommended that the primary analysis should consist of “*log number of breakdowns per trial area in the form: Treatments; Triplets; Treatment x Triplets; Poisson error* “. Including an interactive term of Treatment x Triplet leads to a saturated model (at least 20 predictors for 20 data points) and, hence, is invalid. This is also evidence that the “*independent audit”* was inadequate. The only method by which such an interaction can be analysed is by replacing Triplet with herd years at risk and, therefore, having sufficient degrees of freedom to avoid saturation. Such an analysis demonstrates no evidence for a culling effect (see supplementary information 2).

## 5. Neighbouring area study, Mills et al 2024b

### 5.1. Frequentist approach

The statistical concerns relating to the study of neighbouring areas beyond the boundary of badger cull areas, are similar to those issues found in Mills *et al*. 2024a (4) We summarize them in Table 3. Further details can be found in the supplementary information (supplementary files 1&2). Mills *et al*. 2024b (5) also did not report any effect on total breakdowns, which were analysed in Torgerson *et al*. (3)

**Table 3.**
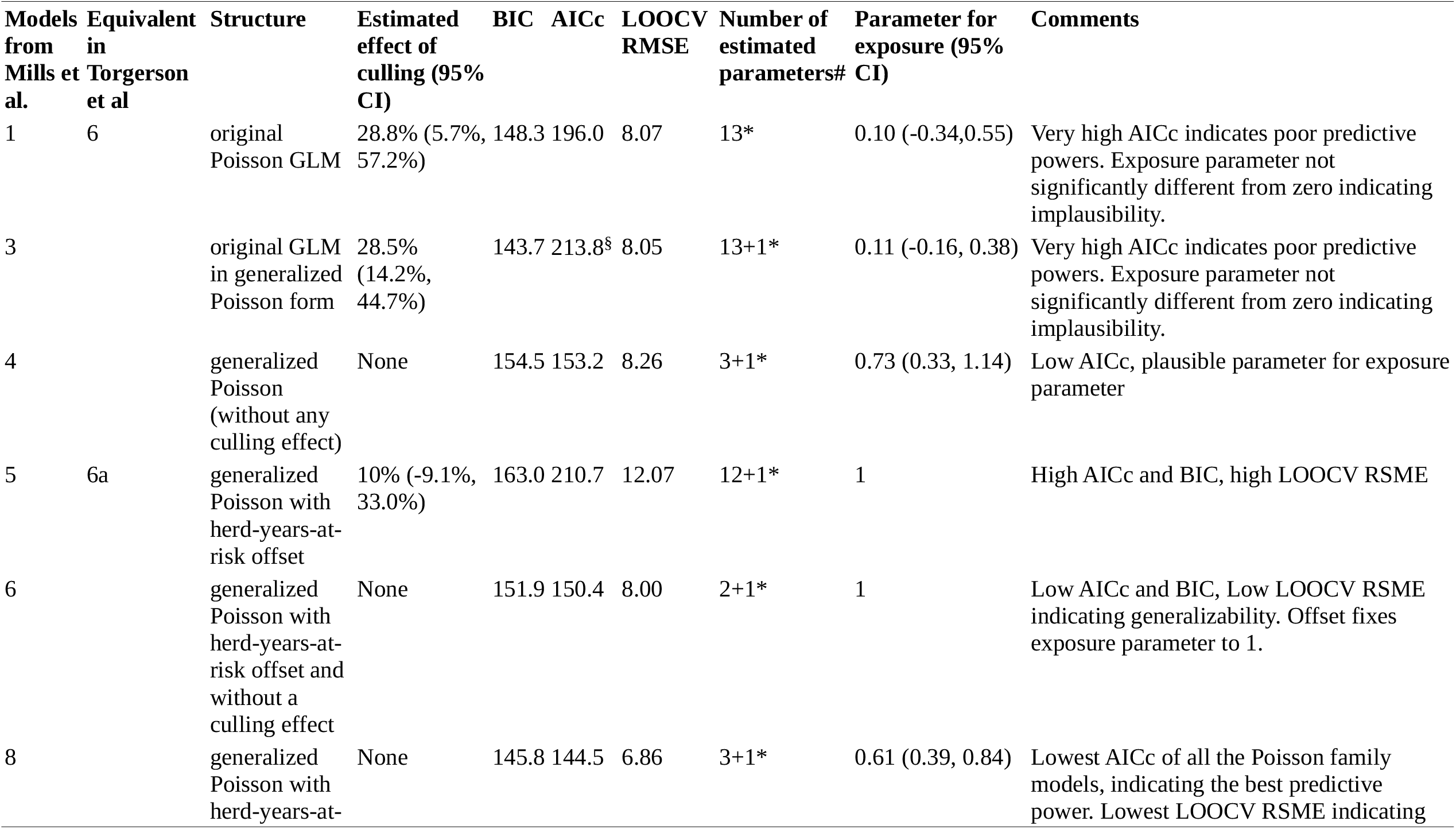
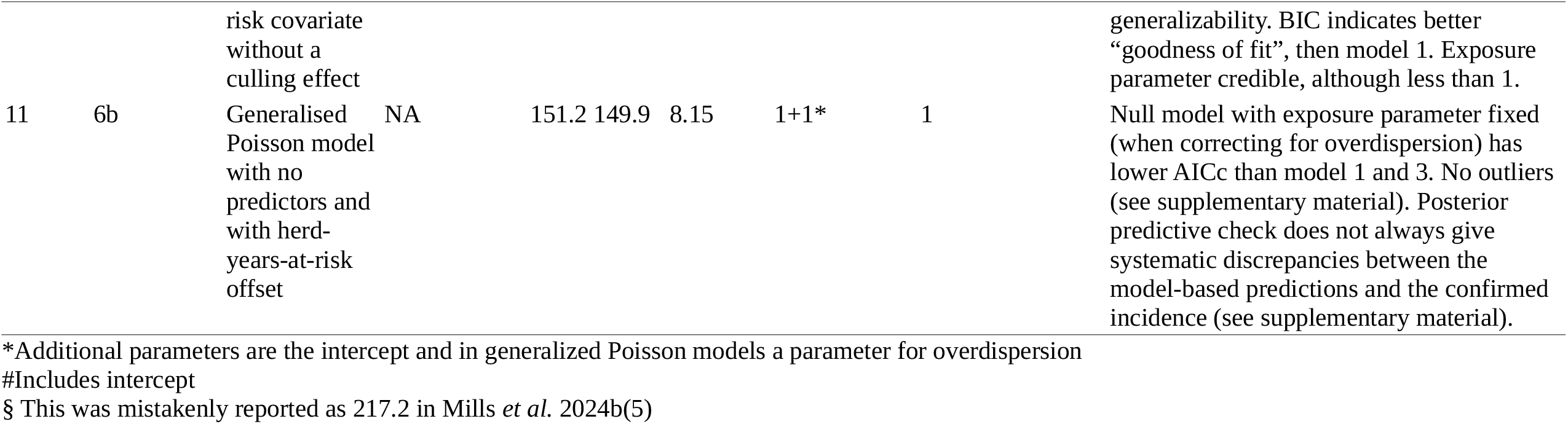
A selection of the frequentist models analysed in Mills *et al*. 2024b(5) and Torgerson *et al*. 2024(3). This compares the original model from the 2006 study(2) (and its equivalent in generalized Poisson form). This is the model that Mills *et al*. 2024b(5) claim to be a robust interpretation of the RBCT results and that “strongly” supports a culling effect. We compare this to those models with the lowest AICc values, which provide no support for an effect of culling.

### 5.2. Bayesian approach

The issues in the Bayesian approach in Mills *et al*. 2024b (5) are similar to those in Mills *et al*. 2023A (4) and are summarized in Table 4. It is worth noting that model d.2, (one model that was correctly coded) by their own analysis, *“ does not contain the implausibly large synthetic model-based predictions; furthermore, the estimated out-of-sample predictive accuracy (measured by LOO ELPD) and, hence, the generalizability of the model are improved”*. Nevertheless, Mills *et al*. dismiss it because it *“does not account for any effect of culling”*. Here the key point is that the modelled incidence is independent of culling (i.e. culling has no effect). Furthermore, this model is supported substantially by Bayes factors compared to the original model used in the RBCT (model e, without offset). Here model d.2. is favoured over model e by a Bayes factor of 183. Such a value is decisive (15) thus completely discounting any effect of culling.

**Table 4:**
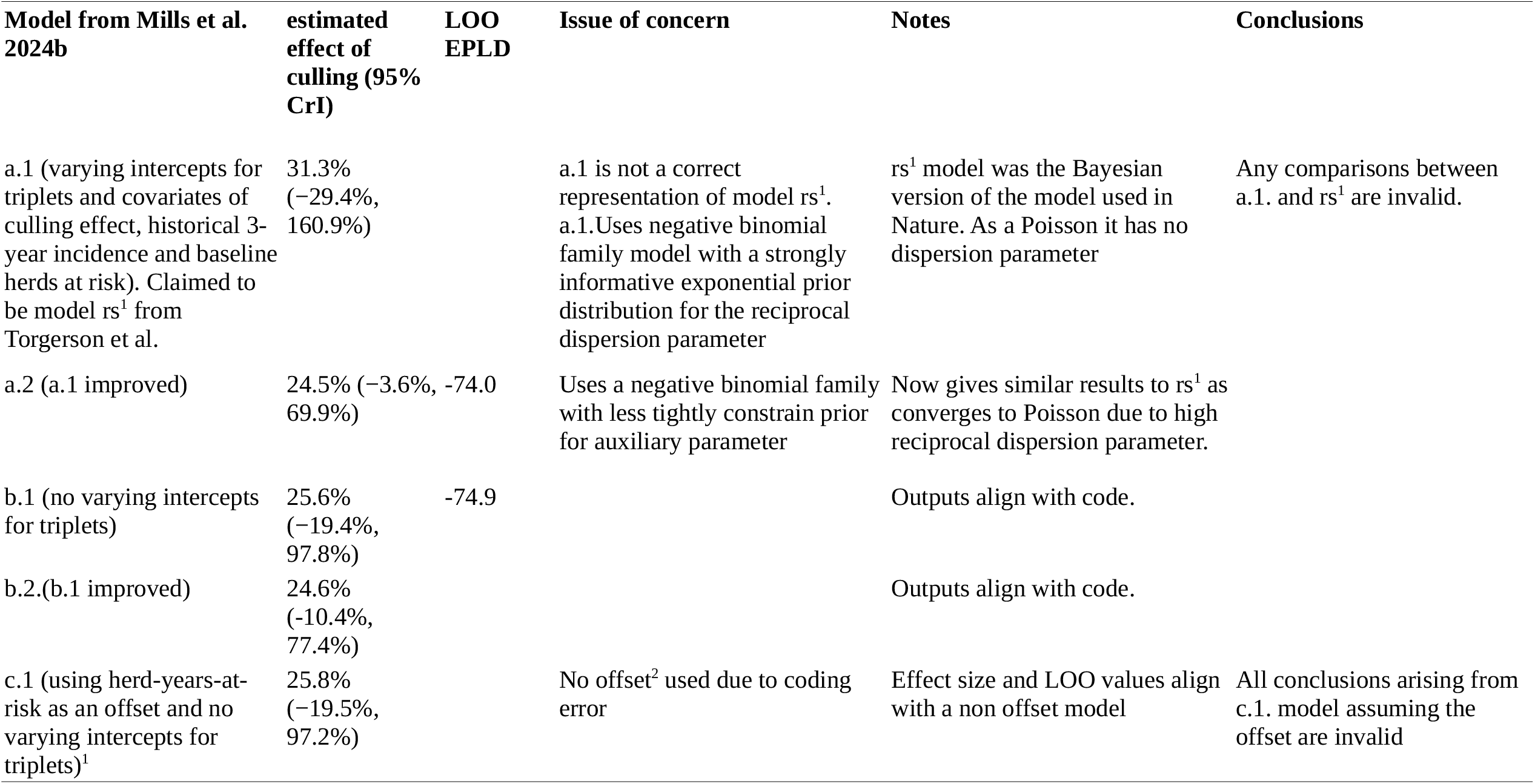
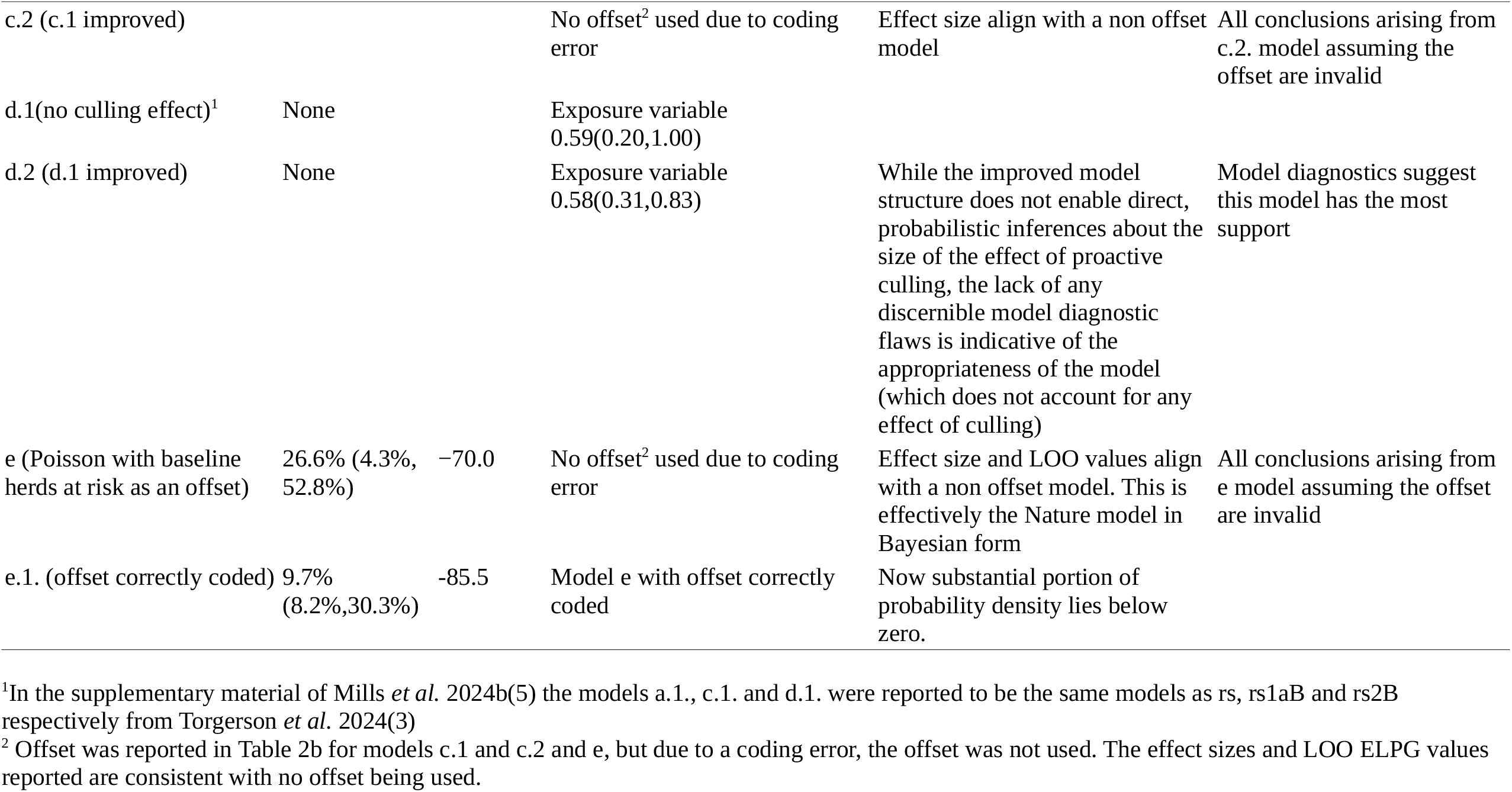
A range of Bayesian models for confirmed bTB herd breakdowns from initial RBCT cull until 4 September 2005, within proactive culling areas. The original, frequentist Poisson GLM used in Donnelly *et al*. 2006(2) was re-specified in the Bayesian paradigm in the initial preprint subsequently published by Torgerson *et al*. (2024)(3). However, the Model rs in Torgerson *et al*. (3) was not specified by a negative binomial likelihood although Mills *et al*. 2024b(5) reported it as Model a.1. in error. Other errors and inconsistencies between those reported in Mills *et al*. 2024b(5) and Torgerson *et al*. (3) for the Bayesian analysis paradigm (tables 2a and 2b in Mills et al 2024b) are summarized here. In total of the 8 models specified in tables 2a and 2b of Mills *et al*. 2024b, 5 had errors. Hence the analysis cannot be relied upon for any of the Bayesian analysis in Mills *et al*. 2024b.

## 6. Scientific Reproducibility

The present study together with those of Mills *et al*. (4,5), Torgerson *et al*. (3) and Donnelly *et al*. makes an important case study with respect to the paradigm of reproducibility and compromises of the FAIR principles(16), as demonstrated by this comment. Also a recent manuscript which implies a reduction in bTB is due to badger culling implemented from 2013 onwards and was “*roughly consistent with previously reported effects of interventions including RBCT* “ (17). However, this interpretation can be dismissed. Badger culling was implemented concurrently with improved cattle measures, such as enhanced testing. The analysis only looked at changes in areas where cattle measures and culling were introduced concomitantly. There was no reference to a comparator where only cattle measures were undertaken throughout the study period. Thus any change in bTB incidence can equally well be attributed to cattle measures rather than badger culling.

## 7. Conclusions

In the frequentist approach to the examination of the 2006 RBCT data (2) both within and beyond badger cull areas, there are 3 main issues: i.) the use of counts rather than rates as the response variable, ii.) over fitting by using too many parameters for the number of data points and iii.) modelling of ‘confirmed’ breakdowns only rather than total (OTFW+OTFS) breakdowns. Mills *et al*. (4,5) fail to address all three issues adequately and use BIC above all other appraisal techniques to “confirm” that the original analysis in Nature was ‘robust’. This is justified on the basis that BIC is optimal for ‘goodness of fit’. In contrast, Torgerson *et al*. (3) addresses these issues and concludes, through the use of AICc and, amongst other evidence, that models with best predictive powers do not show an effect of culling. All models, regardless of method of statistical inference and modelling approach confirm that proactive culling of badgers had no influence on total herd breakdowns.

The Bayesian analysis of Mills *et al*. (4,5) has too many errors to be able to provide a full critique. But referring back to the analysis of Torgerson *et al*. (3) models that show that incidence (rates) are independent of culling are far better supported statistically, such as by Bayes factors, than models which suggest an effect of culling.

Mills *et al*. (4,5) state *“Our extension to a wide array of statistical techniques and study periods allows us to make robust conclusions regarding the effects of proactive badger culling which are informed by consistent scientific evidence from trial data, irrespective of which approach to statistical inference is taken*.*”* This statement is demonstrably untrue because the analysis of *“confirmed breakdowns”* show that the effects of culling are not consistent and are highly dependent on the approach to statistical inference, as demonstrated in the present study, in Torgerson *et al*. (3) and indeed in Mills *et al*. (4,5). However, the finding of an absence of any effect on badger culling on the incidence of bTB, when total breakdowns (i.e. “confirmed” and “unconfirmed” OTF-W +OTF-S) are considered, is a robust conclusion irrespective of which approach to statistical inference is taken.

Donnelly, the senior author of the Mills *et al*. papers, in a commentary in the journal *Biostatistics* stated *“ the suggestion of requiring independent replication of specific statistical analyses as a general check before publication seems not merely unnecessary but a misuse of relatively scarce expertise”*(18). In view of the numerous anomalies in the Bayesian analysis, divergent conclusions dependent on statistical inference and model, and other misconceptions presented in Mills *et al*. (4,5) this idea needs revisiting. In addition, the reviewers of manuscripts under consideration for publication should consider more rigorous checks of statistical analyses.

The RBCT was a relatively small, single study with several destabilising factors that may not have been clear at the time and that interfered with the experiment. With respect to the paradigm of reproducibility and the FAIR principles, the original RBCT analysis and recent efforts to support it are wholly unconvincing. The 2006 conclusion of the RBCT that “*badger culling is unlikely to contribute positively to the control of cattle TB in Britain*” is supported (1). However, the route to such a position is revised in the light of modern veterinary understanding and statistical reappraisal.

## Supporting information

Breakdown data 2013 analysis

Confirmed RBCT Conf after trial

Confirmed VetNet

Confirmed VetNet Followup

Outside Areas RBCT

