## Supplementary Information 2 R Code for "Randomised Badger Culling Trial lacks evidence for proactive badger culling effect on tuberculosis in cattle: comment on Mills et al. 2024, Parts I & II"

### Supplementary material R code

2024-09-11

#### Comment on Mills et al. 2024ab: "Extensive re-evaluation the Randomised Badger Culling Trial (RBCT) fails to show evidence of effects of widespread badger culling on tuberculosis in cattle"


#### Authors: Torgerson PR, Hartnack S, Rasmussen P, Lewis F, O'Donnell P, and TES Langton


Code and explanatory text is given for all models in the text. Where necessary it is important to refer to the models and model code given in Mills et al. 2024ab (https://royalsocietypublishing.org/doi/10.1098/rsos.240386 and https://doi.org/10.1098/rsos.240385) and Torgerson et al. 2024 (https://www.nature.com/articles/s41598-024-67160-0) and their supplementary materials for full understanding.

```
library(glmmTMB)
library(MuMIn)
library(rstanarm)
library(bridgesampling)
library(DHARMa)
library(readxl)
library(AER)
library(vcd)
library(performance)
library(cowplot)
library(rstan)
library(boot)
library(olsrr)
library(lme4)
library(caret)
library(bayesplot)
library(gridExtra)
library(vctrs)
library(purrr)
library(loo)
library(plyr)
```

Data as per Donnelly et al, Nature (2006) http://dx.doi.org/10.1038/nature04454

#### Codebook

| Variable | Value |
|---|---|
| Triplet | This is |
| Treatment | This is |
| Incidence | This is |
| Baseline | This is |
| Hist3yr | This is |
| hdyrsrisk | This is |

```r
# Incidence data from initial cull until September 2005
rbctconf <- data.frame(read_xlsx("confirmed_vetnet.xlsx"))

rbctconf[c(1:6)] <- lapply(rbctconf[c(1:6)], factor)
```

```r
# Relevel triplets and treatment (proactive culling vs survey-only)
# to correspond to Mills et al. and Torgerson et al. and thus, enable model review

rbctconf$Triplet <- relevel(factor(rbctconf$Triplet), ref="J")
rbctconf$Treatment <- relevel(factor(rbctconf$Treatment), ref="Survey-only")
```

### Table 1

#### Model 1. Original Poisson GLM as per Donnelly et al, Nature (2006).

http://dx.doi.org/10.1038/nature04454.

```r
# Fit to initial period
model1 <- glm(Incidence ~ Treatment + Triplet + log(Hist3yr) + log(Baseline),
            family = poisson, data=rbctconf)

summary(model1)
```

```
##
## Call:
## glm(formula = Incidence ~ Treatment + Triplet + log(Hist3yr) +
##     log(Baseline), family = poisson, data = rbctconf)
##
## Coefficients:
##                     Estimate Std. Error z value Pr(>|z|)
## (Intercept)         -0.36167    0.93212  -0.388  0.69801
## TreatmentProactive -0.20661    0.07287  -2.835  0.00458 **
## TripletA           -0.26894    0.23410  -1.149  0.25064
## TripletB            0.15640    0.16604   0.942  0.34621
## TripletC            0.43196    0.16141   2.676  0.00745 **
## TripletD           -0.30400    0.19043  -1.596  0.11041
## TripletE           -0.02578    0.17511  -0.147  0.88298
## TripletF           -0.15938    0.18547  -0.859  0.39017
## TripletG            0.48182    0.19488   2.472  0.01342 *
## TripletH           -0.24769    0.18918  -1.309  0.19044
## TripletI           -0.50242    0.19730  -2.546  0.01088 *
## log(Hist3yr)        1.24144    0.21255   5.841  5.2e-09 ***
## log(Baseline)       0.04678    0.24805   0.189  0.85042
## ---
## Signif. codes:  0 '***' 0.001 '**' 0.01 '*' 0.05 '.' 0.1 ' ' 1
##
## (Dispersion parameter for poisson family taken to be 1)
##
##     Null deviance: 167.3255  on 19  degrees of freedom
## Residual deviance:   5.6596  on  7  degrees of freedom
## AIC: 142.29
##
## Number of Fisher Scoring iterations: 4
```

##### Table 1, model1: Estimated effect of culling (95% CI)

```r
round((exp(coef(model1))[2] - 1),3)*100   # point estimate
```

```
## TreatmentProactive
##              -18.7
```

```r
round((exp(coefci(model1))[2,1]-1),3)*100 # lower 95% CI
```

```
## [1] -29.5
```

```r
round((exp(coefci(model1))[2,2]-1),3)*100 # upper 95% CI
```

```
## [1] -6.2
```

##### Table 1, model1: BIC and AICc

```r
round(BIC(model1),1)
```

```
## [1] 155.2
```

```r
round(AICc(model1),1)
```

```
## [1] 203
```

##### Table 1, model1: LOOCV RMSE

Leave-One-Out Cross-Validation and Root Mean Squared Error, a metric used to evaluate the performance of a model.

```r
rmse <- numeric()
actual_vs_predicted <- data.frame(Actual = numeric(), Predicted = numeric())
for (i in 1:nrow(rbctconf)) {
  # Exclude the ith row
  test_data <- rbctconf[i, ]
  train_data <-rbctconf[-i, ]
  model <-glm(Incidence ~ Treatment + Triplet + log(Hist3yr) + log(Baseline),
              family = poisson, data = train_data)
  predictions <- predict(model, newdata = test_data, type="response")
  rmse[i] <- sqrt(mean((test_data$Incidence - predictions)^2))
  actual_vs_predicted <- rbind(actual_vs_predicted,
                               data.frame(Actual = test_data$Incidence, Predicted = predictions))
}

rmse1<-actual_vs_predicted %>%
  mutate(rmse = sqrt((Actual - Predicted)^2))

round(mean(rmse1$rmse),2)
```

```
## [1] 9.97
```

##### Table 1, model1: Number of explanatory parameters

```r
length(coef(model1))
```

```
## [1] 13
```

##### Table 1, model1: Parameter for exposure (95% CI)

```r
# this is on the log scale
round(coef(model1)[13],2)     # point estimate
```

```
## log(Baseline)
##          0.05
```

```r
round(coefci(model1)[13,1],2) # lower 95% CI
```

```
## [1] -0.44
```

```r
round(coefci(model1)[13,2],2) # upper 95% CI
```

```
## [1] 0.53
```

In the text there is also mentionned this model for other time periods.

```r
### Incidence data from first follow-up cull until September 2005
rbctconf_follow <- data.frame(read_xlsx("confirmed_vetnet_FOLLOW_UP.xlsx"))
rbctconf_follow[c(1:6)] <- lapply(rbctconf_follow[c(1:6)], factor)

# relevel
rbctconf_follow$Triplet <- relevel(factor(rbctconf_follow$Triplet), ref="J")
rbctconf_follow$Treatment <- relevel(factor(rbctconf_follow$Treatment),
                                    ref="Survey-only")

#Follow-Up Period
model1_follow<-glm(Incidence ~ Treatment + Triplet + log(Hist3yr)
                   + log(Baseline),
                   family = poisson, data=rbctconf_follow)

summary(model1_follow)
```

```
##
## Call:
## glm(formula = Incidence ~ Treatment + Triplet + log(Hist3yr) +
##     log(Baseline), family = poisson, data = rbctconf_follow)
##
## Coefficients:
##                    Estimate Std. Error z value Pr(>|z|)
## (Intercept)        -0.90163    1.10251  -0.818  0.41347
## TreatmentProactive -0.25752    0.08434  -3.053  0.00226 **
## TripletA           -0.35918    0.28917  -1.242  0.21421
## TripletB            0.53645    0.19328   2.775  0.00551 **
## TripletC            0.59732    0.18957   3.151  0.00163 **
## TripletD           -0.08866    0.22585  -0.393  0.69462
## TripletE            0.26160    0.20531   1.274  0.20259
## TripletF           -0.27022    0.22761  -1.187  0.23515
## TripletG            0.38358    0.23448   1.636  0.10187
## TripletH           -0.11334    0.22276  -0.509  0.61091
## TripletI           -0.43034    0.23864  -1.803  0.07134 .
## log(Hist3yr)        1.06421    0.24877   4.278 1.89e-05 ***
## log(Baseline)       0.19603    0.29475   0.665  0.50600
## ---
## Signif. codes:  0 '***' 0.001 '**' 0.01 '*' 0.05 '.' 0.1 ' ' 1
##
## (Dispersion parameter for poisson family taken to be 1)
##
##     Null deviance: 177.9342  on 19  degrees of freedom
## Residual deviance:   9.2266  on  7  degrees of freedom
## AIC: 139.3
##
## Number of Fisher Scoring iterations: 4
```

```r
BIC(model1_follow)
```

```
## [1] 152.2405
```

```r
AICc(model1_follow)
```

```
## [1] 199.9626
```

```r
### Incidence data from post-trial period
### (1 year after last proactive cull until 2013)
rbctconf_after <- data.frame(read_xlsx("confirmed_rbctconf_after_trial.xlsx"))
rbctconf_after[c(1:6)] <- lapply(rbctconf_after[c(1:6)], factor)

# relevel
rbctconf_after$Triplet <- relevel(factor(rbctconf_follow$Triplet), ref="J")
rbctconf_after$Treatment <- relevel(factor(rbctconf_follow$Treatment), ref="Survey-only")

#Post trial  period
model1_after<-glm(Incidence ~ Treatment + Triplet + log(Hist3yr) +log(Baseline),
                  family = poisson, data=rbctconf_after)

summary(model1_after)
```

```
##
## Call:
## glm(formula = Incidence ~ Treatment + Triplet + log(Hist3yr) +
##     log(Baseline), family = poisson, data = rbctconf_after)
##
## Coefficients:
##                     Estimate Std. Error z value Pr(>|z|)
## (Intercept)          1.91722    0.70964   2.702 0.006899 **
## TreatmentProactive -0.32062    0.05489  -5.841 5.18e-09 ***
## TripletA           -0.88461    0.17603  -5.025 5.03e-07 ***
## TripletB           -0.45796    0.11837  -3.869 0.000109 ***
## TripletC           -0.04259    0.10831  -0.393 0.694161
## TripletD           -0.59435    0.13319  -4.463 8.10e-06 ***
## TripletE           -0.31929    0.11940  -2.674 0.007494 **
## TripletF           -0.60887    0.13009  -4.680 2.86e-06 ***
## TripletG           -0.01121    0.13411  -0.084 0.933368
## TripletH           -0.31043    0.11978  -2.592 0.009547 **
## TripletI           -0.38088    0.11971  -3.182 0.001464 **
## log(Hist3yr)        0.82165    0.15409   5.332 9.70e-08 ***
## log(Baseline)       0.05880    0.18598   0.316 0.751883
## ---
## Signif. codes:  0 '***' 0.001 '**' 0.01 '*' 0.05 '.' 0.1 ' ' 1
##
## (Dispersion parameter for poisson family taken to be 1)
##
##     Null deviance: 158.345  on 19  degrees of freedom
## Residual deviance:  23.527  on  7  degrees of freedom
## AIC: 172.17
##
## Number of Fisher Scoring iterations: 4
```

```r
BIC(model1_after)
```

```
## [1] 185.1128
```

```r
AICc(model1_after)
```

```
## [1] 232.8349
```

#### Model 3: Original GLM in generalized Poisson form

##### Table 1, model3

```
model3 <- glmmTMB(Incidence ~ Treatment + Triplet + log(Hist3yr) + log(Baseline),
                family = genpois, data=rbctconf)

summary(model3)
```

```
##  Family: genpois  ( log )
## Formula:
## Incidence ~ Treatment + Triplet + log(Hist3yr) + log(Baseline)
## Data: rbctconf
##
##      AIC      BIC   logLik deviance df.resid
##    133.2    147.1    -52.6    105.2        6
##
##
## Dispersion parameter for genpois family (): 0.28
##
## Conditional model:
##                     Estimate Std. Error z value Pr(>|z|)
## (Intercept)         -0.34633    0.50217  -0.690  0.49041
## TreatmentProactive -0.20663    0.03861  -5.351 8.73e-08 ***
## TripletA           -0.27815    0.12361  -2.250  0.02443 *
## TripletB            0.14526    0.08835   1.644  0.10014
## TripletC            0.42270    0.08429   5.015 5.31e-07 ***
## TripletD           -0.31088    0.10055  -3.092  0.00199 **
## TripletE           -0.03455    0.09394  -0.368  0.71303
## TripletF           -0.16994    0.09619  -1.767  0.07728 .
## TripletG            0.47684    0.10351   4.607 4.09e-06 ***
## TripletH           -0.24178    0.10453  -2.313  0.02072 *
## TripletI           -0.52013    0.10479  -4.964 6.92e-07 ***
## log(Hist3yr)        1.24342    0.10867  11.442  < 2e-16 ***
## log(Baseline)       0.04384    0.13139   0.334  0.73865
## ---
## Signif. codes:  0 '***' 0.001 '**' 0.01 '*' 0.05 '.' 0.1 ' ' 1
```

##### Table 1, model3: Estimated effect of culling (95% CI)

```
round((exp(confint(model3, level = 0.95)[2,3])-1),3)*100 # point estimate
```

```
## [1] -18.7
```

```
round((exp(confint(model3, level = 0.95)[2,1])-1),3)*100 # lower 95% CI
```

```
## [1] -24.6
```

```
round((exp(confint(model3, level = 0.95)[2,2])-1),3)*100 # upper 95% CI
```

```
## [1] -12.3
```

##### Table 1, model3: BIC and AICc

```
round(BIC(model3),1)
```

```
## [1] 147.1
```

```r
round(AICc(model3),1)
```

```
## [1] 217.2
```

###### Table 1, model3: LOOCV RMSE

Leave-One-Out Cross-Validation and Root Mean Squared Error, a metric used to evaluate the performance of a model.

```r
rmse <- numeric()
actual_vs_predicted <- data.frame(Actual = numeric(), Predicted = numeric())
for (i in 1:nrow(rbctconf)) {
  # Exclude the ith row
  test_data <- rbctconf[i, ]
  train_data <-rbctconf[-i, ]
  model <-glmmTMB(Incidence ~ Treatment + Triplet + log(Hist3yr) + log(Baseline),
                family = genpois, data = train_data)
  predictions <- predict(model, newdata = test_data, type="response")
  rmse[i] <- sqrt(mean((test_data$Incidence - predictions)^2))
  actual_vs_predicted <- rbind(actual_vs_predicted,
                               data.frame(Actual = test_data$Incidence,
                                          Predicted = predictions))
}

rmse1<-actual_vs_predicted %>%
  mutate(rmse = sqrt((Actual - Predicted)^2))

round(mean(rmse1$rmse),2)
```

```
## [1] 9.95
```

###### Table 1, model3: Number of explanatory parameters

```r
length(fixef(model3)$cond) # plus an additional parameter for overdispersion
```

```
## [1] 13
```

###### Table 1, model3: Parameter for exposure (95% CI)

```r
# this is on the log scale
round(confint(model3, level = 0.95)[13,3],2) # point estimate
```

```
## [1] 0.04
```

```r
round(confint(model3, level = 0.95)[13,1],2) # lower 95% CI
```

```
## [1] -0.21
```

```r
round(confint(model3, level = 0.95)[13,2],2) # upper 95% CI
```

```
## [1] 0.3
```

In the text there is also mentionned this model for other time periods.

```r
#Follow-Up Period
model3_follow <- glmmTMB(Incidence ~ Treatment + Triplet + log(Hist3yr)
                         + log(Baseline),
                        family = genpois, data=rbctconf_follow)

summary(model3_follow)
```

```
##  Family: genpois  ( log )
## Formula:
## Incidence ~ Treatment + Triplet + log(Hist3yr) + log(Baseline)
## Data: rbctconf_follow
##
##      AIC      BIC   logLik deviance df.resid
##    136.5    150.4    -54.2    108.5        6
##
##
## Dispersion parameter for genpois family (): 0.457
##
## Conditional model:
##                     Estimate Std. Error z value Pr(>|z|)
## (Intercept)         -0.87925    0.74974  -1.173  0.24090
## TreatmentProactive -0.25661    0.05720  -4.486 7.26e-06 ***
## TripletA            -0.36963    0.19469  -1.899  0.05762 .
## TripletB             0.52336    0.13172   3.973 7.09e-05 ***
## TripletC             0.58992    0.12679   4.653 3.28e-06 ***
## TripletD            -0.08304    0.15146  -0.548  0.58348
## TripletE             0.24708    0.14021   1.762  0.07804 .
## TripletF            -0.28452    0.15231  -1.868  0.06176 .
## TripletG             0.37635    0.15756   2.389  0.01691 *
## TripletH            -0.11479    0.15441  -0.743  0.45726
## TripletI            -0.45162    0.16199  -2.788  0.00531 **
## log(Hist3yr)         1.07196    0.16593   6.460 1.05e-10 ***
## log(Baseline)        0.18787    0.19769   0.950  0.34194
## ---
## Signif. codes:  0 '***' 0.001 '**' 0.01 '*' 0.05 '.' 0.1 ' ' 1
```

```r
BIC(model3_follow)
```

```
## [1] 150.4353
```

```r
AICc(model3_follow)
```

```
## [1] 220.495
```

```r
#Post trial  period
model3_after <- glmmTMB(Incidence ~ Treatment + Triplet + log(Hist3yr)
                        + log(Baseline),
                       family = genpois, data=rbctconf_after)

summary(model3_after)
```

```
##  Family: genpois  ( log )
## Formula:
## Incidence ~ Treatment + Triplet + log(Hist3yr) + log(Baseline)
## Data: rbctconf_after
```

```
##
##      AIC      BIC   logLik deviance df.resid
##    173.9    187.8    -72.9    145.9        6
##
##
## Dispersion parameter for genpois family (): 1.18
##
## Conditional model:
##                     Estimate Std. Error z value Pr(>|z|)
## (Intercept)          1.92029    0.77258   2.486 0.012935 *
## TreatmentProactive -0.32144    0.05978  -5.378 7.55e-08 ***
## TripletA           -0.88584    0.19110  -4.635 3.56e-06 ***
## TripletB           -0.45891    0.12841  -3.574 0.000352 ***
## TripletC           -0.04316    0.11773  -0.367 0.713921
## TripletD           -0.59753    0.14499  -4.121 3.77e-05 ***
## TripletE           -0.32437    0.13082  -2.479 0.013157 *
## TripletF           -0.60971    0.14142  -4.311 1.62e-05 ***
## TripletG           -0.01144    0.14656  -0.078 0.937773
## TripletH           -0.31062    0.13018  -2.386 0.017030 *
## TripletI           -0.38197    0.13024  -2.933 0.003359 **
## log(Hist3yr)        0.82425    0.16691   4.938 7.89e-07 ***
## log(Baseline)       0.05677    0.20248   0.280 0.779171
## ---
## Signif. codes:  0 '***' 0.001 '**' 0.01 '*' 0.05 '.' 0.1 ' ' 1
```

```
BIC(model3_after)
```

```
## [1] 187.8063
```

```
AICc(model3_after)
```

```
## [1] 257.866
```

#### Model 4: Generalized Poisson (without any culling effect)

##### Tabe 1, model4

```
model4 <- glmmTMB(Incidence ~ log(Hist3yr) + log(Baseline), family = genpois,
                  data=rbctconf)

summary(model4)
```

```
##  Family: genpois  ( log )
## Formula:          Incidence ~ log(Hist3yr) + log(Baseline)
## Data: rbctconf
##
##      AIC      BIC   logLik deviance df.resid
##    157.7    161.7    -74.9    149.7       16
##
##
## Dispersion parameter for genpois family (): 2.63
##
## Conditional model:
##               Estimate Std. Error z value Pr(>|z|)
## (Intercept)    -2.4021     0.9581  -2.507   0.0122 *
## log(Hist3yr)    0.9352     0.1938   4.825 1.40e-06 ***
## log(Baseline)   0.6569     0.1650   3.982 6.83e-05 ***
## ---
## Signif. codes:  0 '***' 0.001 '**' 0.01 '*' 0.05 '.' 0.1 ' ' 1
```

```
confint(model4)
```

```
##                    2.5 %     97.5 %   Estimate
## (Intercept)   -4.2799350 -0.5241955 -2.4020653
## log(Hist3yr)   0.5553377  1.3151395  0.9352386
## log(Baseline)  0.3335948  0.9802854  0.6569401
```

##### Table 1, model4: BIC and AICc

```
round(BIC(model4),1)
```

```
## [1] 161.7
```

```
round(AICc(model4),1)
```

```
## [1] 160.4
```

##### Table 1, model4: LOOCV RMSE

Leave-One-Out Cross-Validation and Root Mean Squared Error, a metric used to evaluate the performance of a model.

```
rmse <- numeric()
actual_vs_predicted <- data.frame(Actual = numeric(), Predicted = numeric())
for (i in 1:nrow(rbctconf)) {
  # Exclude the ith row
  test_data <- rbctconf[i, ]
  train_data <-rbctconf[-i, ]
  model <- glmmTMB(Incidence ~ log(Hist3yr) + log(Baseline), family = genpois,
                   data = train_data)
```

```
  predictions <- predict(model, newdata = test_data, type="response")
  rmse[i] <- sqrt(mean((test_data$Incidence - predictions)^2))
  actual_vs_predicted <- rbind(actual_vs_predicted,
                               data.frame(Actual = test_data$Incidence,
                                          Predicted = predictions))
}

rmse1<-actual_vs_predicted %>%
  mutate(rmse = sqrt((Actual - Predicted)^2))

round(mean(rmse1$rmse),2)
```

```
## [1] 10.93
```

###### Table 1, model4: Number of explanatory parameters

```
length(fixef(model4)$cond) # plus an additional parameter for overdispersion
```

```
## [1] 3
```

###### Table 1, model4: Parameter for exposure (95% CI)

```
# this is on the log scale
round(confint(model4, level = 0.95)[3,3],2) # point estimate
```

```
## [1] 0.66
```

```
round(confint(model4, level = 0.95)[3,1],2) # lower 95% CI
```

```
## [1] 0.33
```

```
round(confint(model4, level = 0.95)[3,2],2) # upper 95% CI
```

```
## [1] 0.98
```

In the text there is also mentionned this model for other time periods.

```r
#Follow-Up Period
model4_follow <- glmmTMB(Incidence ~ log(Hist3yr) + log(Baseline),
                         family = genpois, data=rbctconf_follow)


summary(model4_follow)
```

```
##  Family: genpois  ( log )
## Formula:          Incidence ~ log(Hist3yr) + log(Baseline)
## Data: rbctconf_follow
##
##      AIC      BIC   logLik deviance df.resid
##    159.3    163.3    -75.7    151.3       16
##
##
## Dispersion parameter for genpois family (): 4.14
##
## Conditional model:
##               Estimate Std. Error z value Pr(>|z|)
## (Intercept)    -2.7461     1.3467  -2.039 0.041436 *
## log(Hist3yr)    0.8968     0.2723   3.294 0.000989 ***
## log(Baseline)   0.6935     0.2233   3.106 0.001898 **
## ---
## Signif. codes:  0 '***' 0.001 '**' 0.01 '*' 0.05 '.' 0.1 ' ' 1
```

```r
BIC(model4_follow)
```

```
## [1] 163.3179
```

```r
AICc(model4_follow)
```

```
## [1] 162.0016
```

```r
#Post trial  period
model4_after <- glmmTMB(Incidence ~ log(Hist3yr) + log(Baseline),
                        family = genpois, data=rbctconf_after)

summary(model4_after)
```

```
##  Family: genpois  ( log )
## Formula:          Incidence ~ log(Hist3yr) + log(Baseline)
## Data: rbctconf_after
##
##      AIC      BIC   logLik deviance df.resid
##    186.5    190.5    -89.2    178.5       16
##
##
## Dispersion parameter for genpois family (): 5.95
##
## Conditional model:
##               Estimate Std. Error z value Pr(>|z|)
## (Intercept)     0.8870     1.0819   0.820   0.4123
## log(Hist3yr)    0.4431     0.1911   2.319   0.0204 *
## log(Baseline)   0.4248     0.1829   2.322   0.0202 *
## ---
```

```
## Signif. codes:  0 '***' 0.001 '**' 0.01 '*' 0.05 '.' 0.1 ' ' 1
```

```r
BIC(model4_after)
```

```
## [1] 190.46
```

```r
AICc(model4_after)
```

```
## [1] 189.1437
```

#### Model 8: Generalized Poisson with herd-years-at-risk covariate without any culling effect

##### Table 1, model8

```r
model8 <- glmmTMB(Incidence ~ log(Hist3yr) + log(hdyrsrisk), family = genpois,
                  data=rbctconf)

summary(model8)
```

```
##  Family: genpois  ( log )
## Formula:          Incidence ~ log(Hist3yr) + log(hdyrsrisk)
## Data: rbctconf
##
##      AIC      BIC   logLik deviance df.resid
##    151.5    155.5    -71.8    143.5       16
##
##
## Dispersion parameter for genpois family (): 1.89
##
## Conditional model:
##                Estimate Std. Error z value Pr(>|z|)
## (Intercept)    -2.13664    0.71509  -2.988  0.00281 **
## log(Hist3yr)    0.82669    0.16096   5.136 2.81e-07 ***
## log(hdyrsrisk)  0.51201    0.09616   5.325 1.01e-07 ***
## ---
## Signif. codes:  0 '***' 0.001 '**' 0.01 '*' 0.05 '.' 0.1 ' ' 1
```

##### Table 1, model8: BIC and AICc

```r
round(BIC(model8),1)
```

```
## [1] 155.5
```

```r
round(AICc(model8),1)
```

```
## [1] 154.2
```

##### Table 1, model8: LOOCV RMSE

Leave-One-Out Cross-Validation and Root Mean Squared Error, a metric used to evaluate the performance of a model.

```r
rmse <- numeric()
actual_vs_predicted <- data.frame(Actual = numeric(), Predicted = numeric())
for (i in 1:nrow(rbctconf)) {
  # Exclude the ith row
```

```
  test_data <- rbctconf[i, ]
  train_data <-rbctconf[-i, ]
  model <- glmmTMB(Incidence ~ log(Hist3yr) + log(hdyrsrisk), family = genpois,
                   data = train_data)
  predictions <- predict(model, newdata = test_data, type="response")
  rmse[i] <- sqrt(mean((test_data$Incidence - predictions)^2))
  actual_vs_predicted <- rbind(actual_vs_predicted,
                               data.frame(Actual = test_data$Incidence,
                                          Predicted = predictions))
}

rmse1<-actual_vs_predicted %>%
  mutate(rmse = sqrt((Actual - Predicted)^2))

round(mean(rmse1$rmse),2)
```

```
## [1] 8.81
```

##### Table 1, model8: Number of explanatory parameters

```
length(fixef(model8)$cond) # plus an additional parameter for overdispersion
```

```
## [1] 3
```

##### Table 1, model8: Parameter for exposure (95% CI)

```
# this is on the log scale
round(confint(model8, level = 0.95)[3,3],2) # point estimate
```

```
## [1] 0.51
```

```
round(confint(model8, level = 0.95)[3,1],2) # lower 95% CI
```

```
## [1] 0.32
```

```
round(confint(model8, level = 0.95)[3,2],2) # upper 95% CI
```

```
## [1] 0.7
```

In the text there is also mentionned this model for other time periods.

```r
#Follow-Up Period
model8_follow <- glmmTMB(Incidence ~ log(Hist3yr) + log(hdyrsrisk),
                       family = genpois, data=rbctconf_follow)

summary(model8_follow)
```

```
##  Family: genpois  ( log )
## Formula:          Incidence ~ log(Hist3yr) + log(hdyrsrisk)
## Data: rbctconf_follow
##
##      AIC      BIC   logLik deviance df.resid
##    146.2    150.2    -69.1    138.2       16
##
##
## Dispersion parameter for genpois family ():    2
##
## Conditional model:
##                Estimate Std. Error z value Pr(>|z|)
## (Intercept)     -2.9200     0.7875  -3.708 0.000209 ***
## log(Hist3yr)     0.7328     0.1898   3.861 0.000113 ***
## log(hdyrsrisk)   0.6684     0.1105   6.049 1.46e-09 ***
## ---
## Signif. codes:  0 '***' 0.001 '**' 0.01 '*' 0.05 '.' 0.1 ' ' 1
```

```r
BIC(model8_follow)
```

```
## [1] 150.1776
```

```r
AICc(model8_follow)
```

```
## [1] 148.8614
```

```r
#Post trial  period
model8_after <- glmmTMB(Incidence ~ log(Hist3yr) + log(hdyrsrisk),
                      family = genpois, data=rbctconf_after)

summary(model8_after)
```

```
##  Family: genpois  ( log )
## Formula:          Incidence ~ log(Hist3yr) + log(hdyrsrisk)
## Data: rbctconf_after
##
##      AIC      BIC   logLik deviance df.resid
##    186.5    190.5    -89.2    178.5       16
##
##
## Dispersion parameter for genpois family (): 5.95
##
## Conditional model:
##                Estimate Std. Error z value Pr(>|z|)
## (Intercept)     0.08527    1.37874   0.062   0.9507
## log(Hist3yr)    0.44314    0.19111   2.319   0.0204 *
## log(hdyrsrisk)  0.42485    0.18294   2.322   0.0202 *
## ---
## Signif. codes:  0 '***' 0.001 '**' 0.01 '*' 0.05 '.' 0.1 ' ' 1
```

```r
BIC(model8_after)
```

```
## [1] 190.46
```

```r
AICc(model8_after)
```

```
## [1] 189.1437
```

#### Null model without any predictors

##### Table 1, null model

```r
model.null <- glmmTMB(Incidence ~ 1, family = genpois,
                  data=rbctconf)
summary(model.null)
```

```
##  Family: genpois  ( log )
## Formula:          Incidence ~ 1
## Data: rbctconf
##
##      AIC      BIC   logLik deviance df.resid
##    176.1    178.1    -86.0    172.1       18
##
##
## Dispersion parameter for genpois family ():  8.7
##
## Conditional model:
##             Estimate Std. Error z value Pr(>|z|)
## (Intercept)   3.7865     0.0993   38.13   <2e-16 ***
## ---
## Signif. codes:  0 '***' 0.001 '**' 0.01 '*' 0.05 '.' 0.1 ' ' 1
```

##### Table 1, model null: BIC and AICc

```r
round(BIC(model.null),1)
```

```
## [1] 178.1
```

```r
round(AICc(model.null),1)
```

```
## [1] 176.8
```

##### Table 1, model null: LOOCV RMSE

Leave-One-Out Cross-Validation and Root Mean Squared Error, a metric used to evaluate the performance of a model.

```r
rmse <- numeric()
actual_vs_predicted <- data.frame(Actual = numeric(), Predicted = numeric())
for (i in 1:nrow(rbctconf)) {
  # Exclude the ith row
  test_data <- rbctconf[i, ]
  train_data <-rbctconf[-i, ]
  model <- glmmTMB(Incidence ~ 1, family = genpois,
                  data = train_data)
  predictions <- predict(model, newdata = test_data, type="response")
```

```
  rmse[i] <- sqrt(mean((test_data$Incidence - predictions)^2))
  actual_vs_predicted <- rbind(actual_vs_predicted,
                               data.frame(Actual = test_data$Incidence,
                                          Predicted = predictions))
}

rmse1<-actual_vs_predicted %>%
  mutate(rmse = sqrt((Actual - Predicted)^2))

round(mean(rmse1$rmse),2)
```

```
## [1] 17.29
```

##### Table 1, model8: Number of explanatory parameters

```
length(fixef(model.null)$cond) # plus an additional parameter in generalized Poisson model to model ove[illegible]
```

```
## [1] 1
```

### Table 2: Bayesian analysis

#### Model a.1 as coded by Mills et al.

Note the stan_glm.nb function, which is modelling a negative binomial regression

```
modela1 <- stan_glm.nb(Incidence~Treatment+A+B+C+D+E+F+G+H+I+log(Hist3yr)+log(Baseline),
                  data = rbctconf, prior_intercept=normal(0, 10),
                  diagnostic_file = file.path(tempdir(), "df.csv"),refresh=0)

summary(modela1, digits=5, probs=c(0.025,0.5,0.975))
```

```
##
## Model Info:
##  function:     stan_glm.nb
##  family:       neg_binomial_2 [log]
##  formula:      Incidence ~ Treatment + A + B + C + D + E + F + G + H + I + log(Hist3yr) +
##     log(Baseline)
##  algorithm:    sampling
##  sample:       4000 (posterior sample size)
##  priors:       see help('prior_summary')
##  observations: 20
##  predictors:   13
##
## Estimates:
##                         mean     sd       2.5%     50%      97.5%
## (Intercept)           -0.06708  3.52495 -7.17146 -0.07284  6.73558
## TreatmentProactive    -0.20535  0.27071 -0.73973 -0.20502  0.31439
## A1                    -0.29127  0.83268 -1.92354 -0.30926  1.36874
## B1                     0.15820  0.64014 -1.11920  0.16349  1.43737
## C1                     0.40427  0.58477 -0.76292  0.42182  1.52893
## D1                    -0.29965  0.66464 -1.56991 -0.31911  1.02903
## E1                    -0.05241  0.64950 -1.30123 -0.05827  1.23193
## F1                    -0.09735  0.65034 -1.37849 -0.10682  1.23299
## G1                     0.48689  0.69170 -0.89437  0.47397  1.86941
## H1                    -0.22158  0.67099 -1.50458 -0.23705  1.17010
## I1                    -0.50785  0.64017 -1.72746 -0.50499  0.74152
## log(Hist3yr)           1.21676  0.67788 -0.13495  1.22182  2.56233
## log(Baseline)          0.01977  0.88812 -1.74762  0.01332  1.79201
## reciprocal_dispersion  4.28657  1.93894  1.50823  3.97269  8.87496
##
## Fit Diagnostics:
##            mean     sd       2.5%     50%      97.5%
## mean_PPD 54.88424 24.25094 35.19750 51.27500 92.05750
##
## The mean_ppd is the sample average posterior predictive distribution of the outcome variable (for de
##
## MCMC diagnostics
##                       mcse    Rhat    n_eff
## (Intercept)           0.08697 0.99958 1643
## TreatmentProactive    0.00538 1.00119 2535
## A1                    0.02601 1.00173 1025
## B1                    0.01727 1.00134 1374
## C1                    0.01488 1.00094 1544
## D1                    0.01884 1.00137 1245
```

```
## E1                     0.01857 1.00152 1223
## F1                     0.01588 1.00024 1678
## G1                     0.01767 1.00050 1532
## H1                     0.01921 1.00086 1220
## I1                     0.01767 1.00155 1312
## log(Hist3yr)           0.01811 1.00077 1401
## log(Baseline)          0.02637 1.00009 1135
## reciprocal_dispersion 0.05489 0.99992 1248
## mean_PPD               0.62502 1.00238 1505
## log-posterior          0.16696 1.00316  588
##
## For each parameter, mcse is Monte Carlo standard error, n_eff is a crude measure of effective sample
```

```
#This code allows the software to provide minimally informative priors automatically

prior_summary(modela1)
```

```
## Priors for model 'modela1'
## ------
## Intercept (after predictors centered)
##  ~ normal(location = 0, scale = 10)
##
## Coefficients
##   Specified prior:
##     ~ normal(location = [0,0,0,...], scale = [2.5,2.5,2.5,...])
##   Adjusted prior:
##     ~ normal(location = [0,0,0,...], scale = [4.87,8.12,8.12,...])
##
## Auxiliary (reciprocal_dispersion)
##  ~ exponential(rate = 1)
## ------
## See help('prior_summary.stanreg') for more details
```

```
#prior for auxiliary (reciprocal dispersion) ~ exponential(rate1)
#This strongly informative prior forces a negative binomial model,
#when the data is consistent with a poisson.
#This was not the model rs proposed by Torgerson et al.
```

##### Table 2, Modelrs as coded by Torgerson et al.

This is the properly coded model proposed by Torgerson et al, with weakly informative priors, note the stan_glm function and poisson family defined

```
modelrs<-stan_glm(Incidence~Treatment+A+B+C+D+E+F+G+H+I+log(Hist3yr)+log(Baseline),poisson,
                     data = rbctconf, prior_intercept=normal(0, 10),
                     diagnostic_file = file.path(tempdir(), "df.csv"),refresh=0)

summary(modelrs, digits=5, probs=c(0.025,0.5,0.975))
```

```
##
## Model Info:
##  function:     stan_glm
##  family:       poisson [log]
##  formula:      Incidence ~ Treatment + A + B + C + D + E + F + G + H + I + log(Hist3yr) +
##     log(Baseline)
##  algorithm:    sampling
```

```
##  sample:       4000 (posterior sample size)
##  priors:       see help('prior_summary')
##  observations: 20
##  predictors:   13
##
## Estimates:
##                      mean     sd       2.5%     50%      97.5%
## (Intercept)        -0.39014  0.92750 -2.30381 -0.38599  1.37531
## TreatmentProactive -0.20952  0.07542 -0.36094 -0.21024 -0.06003
## A1                 -0.26715  0.24080 -0.72590 -0.26973  0.19937
## B1                  0.16123  0.16818 -0.16529  0.15971  0.48370
## C1                  0.42862  0.16014  0.11019  0.42912  0.74792
## D1                 -0.30485  0.19263 -0.68336 -0.30479  0.06795
## E1                 -0.02484  0.17603 -0.37091 -0.02495  0.32155
## F1                 -0.16716  0.18517 -0.53370 -0.16124  0.19512
## G1                  0.48009  0.19515  0.10731  0.48123  0.86872
## H1                 -0.25309  0.19270 -0.63361 -0.25536  0.12581
## I1                 -0.50634  0.20276 -0.90556 -0.50118 -0.11386
## log(Hist3yr)        1.24364  0.21745  0.82493  1.24175  1.66946
## log(Baseline)       0.04995  0.24913 -0.44270  0.04446  0.54209
##
## Fit Diagnostics:
##            mean     sd       2.5%     50%      97.5%
## mean_PPD 44.08219  2.11880 40.00000 44.10000 48.40000
##
## The mean_ppd is the sample average posterior predictive distribution of the outcome variable (for de
##
## MCMC diagnostics
##                    mcse    Rhat    n_eff
## (Intercept)        0.01887 1.00035 2415
## TreatmentProactive 0.00129 1.00050 3414
## A1                 0.00671 1.00193 1288
## B1                 0.00436 1.00094 1490
## C1                 0.00377 1.00005 1804
## D1                 0.00498 1.00166 1494
## E1                 0.00467 1.00182 1421
## F1                 0.00447 1.00123 1714
## G1                 0.00462 1.00062 1783
## H1                 0.00455 1.00118 1792
## I1                 0.00474 1.00162 1826
## log(Hist3yr)       0.00532 1.00086 1669
## log(Baseline)      0.00633 1.00100 1551
## mean_PPD           0.03368 1.00030 3957
## log-posterior      0.06000 1.00104 1793
##
## For each parameter, mcse is Monte Carlo standard error, n_eff is a crude measure of effective sample
```

##### Table 2, Model b.1.

```
modelb1 <- stan_glm.nb(Incidence ~ Treatment + log(hdyrsrisk) + log(Hist3yr),
                  prior_intercept=normal(0,10), prior=normal(0,10),
                  data=rbctconf,diagnostic_file =file.path(tempdir(),"df.csv"),
                  refresh=0)
```

```r
summary(modelb1, digits=5, probs=c(0.025,0.5,0.975))
```

```
##
## Model Info:
##  function:     stan_glm.nb
##  family:       neg_binomial_2 [log]
##  formula:      Incidence ~ Treatment + log(hdyrsrisk) + log(Hist3yr)
##  algorithm:    sampling
##  sample:       4000 (posterior sample size)
##  priors:       see help('prior_summary')
##  observations: 20
##  predictors:   4
##
## Estimates:
##                         mean     sd       2.5%     50%      97.5%
## (Intercept)           -2.07591  1.63930 -5.30776 -2.03944  1.20386
## TreatmentProactive    -0.15702  0.21117 -0.57691 -0.15714  0.24880
## log(hdyrsrisk)         0.49058  0.20541  0.09623  0.49581  0.89921
## log(Hist3yr)           0.87729  0.36190  0.16165  0.88680  1.58548
## reciprocal_dispersion  6.22191  2.11072  2.88045  5.93113 11.02304
##
## Fit Diagnostics:
##            mean     sd       2.5%     50%      97.5%
## mean_PPD 45.78733  7.43645 33.30000 45.00000 62.30375
##
## The mean_ppd is the sample average posterior predictive distribution of the outcome variable (for de
##
## MCMC diagnostics
##                       mcse    Rhat    n_eff
## (Intercept)           0.02301 0.99943 5077
## TreatmentProactive    0.00320 0.99942 4353
## log(hdyrsrisk)        0.00300 0.99971 4675
## log(Hist3yr)          0.00585 0.99985 3822
## reciprocal_dispersion 0.03557 1.00046 3521
## mean_PPD              0.12088 0.99967 3785
## log-posterior         0.04676 1.00306 1440
##
## For each parameter, mcse is Monte Carlo standard error, n_eff is a crude measure of effective sample
```

```r
#Effect size does not appear to align with that reported as model rsB1 in Torgerson et al.
```

#### Table 2 model c.1. in table 2 in the text

```r
#Code as reported by Mills et al. Note log(hdyrsrisk) is both an explanatory
#variable and an offset variable
modelc1 <- stan_glm.nb(Incidence ~ Treatment + log(hdyrsrisk) + log(Hist3yr),
                   offset = log(hdyrsrisk),
                   prior_intercept=normal(0,10), prior=normal(0,10),
                   data = rbctconf, refresh=0)

summary(modelc1, digits=5, probs=c(0.025,0.5,0.975))
```

```
##
```

```
## Model Info:
##  function:     stan_glm.nb
##  family:       neg_binomial_2 [log]
##  formula:      Incidence ~ Treatment + log(hdyrsrisk) + log(Hist3yr)
##  algorithm:    sampling
##  sample:       4000 (posterior sample size)
##  priors:       see help('prior_summary')
##  observations: 20
##  predictors:   4
##
## Estimates:
##                         mean     sd       2.5%     50%      97.5%
## (Intercept)           -2.09018  1.62146 -5.25976 -2.10358  1.12624
## TreatmentProactive    -0.15863  0.20170 -0.57134 -0.15721  0.22467
## log(hdyrsrisk)        -0.51665  0.20822 -0.93850 -0.51580 -0.11480
## log(Hist3yr)           0.89674  0.34887  0.20559  0.89482  1.56538
## reciprocal_dispersion  6.23248  2.11588  2.93313  5.96128 11.02562
##
## Fit Diagnostics:
##            mean     sd       2.5%     50%      97.5%
## mean_PPD 45.92026  7.41117 33.35000 45.15000 62.85000
##
## The mean_ppd is the sample average posterior predictive distribution of the outcome variable (for de
##
## MCMC diagnostics
##                       mcse    Rhat    n_eff
## (Intercept)           0.02366 0.99982 4695
## TreatmentProactive    0.00291 0.99970 4791
## log(hdyrsrisk)        0.00328 1.00058 4024
## log(Hist3yr)          0.00514 0.99947 4609
## reciprocal_dispersion 0.03455 1.00060 3750
## mean_PPD              0.11719 0.99973 3999
## log-posterior         0.04450 1.00112 1566
##
## For each parameter, mcse is Monte Carlo standard error, n_eff is a crude measure of effective sample
```

```
loo(modelc1)
```

```
##
## Computed from 4000 by 20 log-likelihood matrix.
##
##          Estimate  SE
## elpd_loo    -81.6 1.8
## p_loo         1.7 0.3
## looic       163.1 3.6
## ------
## MCSE of elpd_loo is 0.0.
## MCSE and ESS estimates assume MCMC draws (r_eff in [0.5, 1.3]).
##
## All Pareto k estimates are good (k < 0.7).
## See help('pareto-k-diagnostic') for details.
```

```
#Summary gives parameter value for explanatory variable log(hdyrsrisk)
#Parameter value for Treatment and LOO ELPD align with values reported
#in table 2a in Mills et al.
```

```r
#The correct code so that log(hdyrsrisk) is an offset variable is:
modelc1 <- stan_glm.nb(Incidence ~ Treatment + log(Hist3yr),
                       offset = log(hdyrsrisk),
                       prior_intercept=normal(0,10), prior=normal(0,10),
                       data = rbctconf, refresh=0)

#Same coding issue for follow up and post trial periods in code written by Mills et al.

summary(modelc1, digits=5, probs=c(0.025,0.5,0.975))
```

```
##
## Model Info:
##  function:     stan_glm.nb
##  family:       neg_binomial_2 [log]
##  formula:      Incidence ~ Treatment + log(Hist3yr)
##  algorithm:    sampling
##  sample:       4000 (posterior sample size)
##  priors:       see help('prior_summary')
##  observations: 20
##  predictors:   3
##
## Estimates:
##                         mean     sd       2.5%     50%      97.5%
## (Intercept)           -4.99059  1.31203 -7.57166 -5.02475 -2.24850
## TreatmentProactive    -0.10755  0.22488 -0.56495 -0.10892  0.32697
## log(Hist3yr)           0.79043  0.40696 -0.05465  0.79295  1.60527
## reciprocal_dispersion  4.91399  1.61190  2.32492  4.72747  8.53512
##
## Fit Diagnostics:
##            mean     sd       2.5%     50%      97.5%
## mean_PPD 51.35446  9.54913 35.74875 50.10000 73.35375
##
## The mean_ppd is the sample average posterior predictive distribution of the outcome variable (for de
##
## MCMC diagnostics
##                       mcse    Rhat    n_eff
## (Intercept)           0.01956 0.99990 4500
## TreatmentProactive    0.00362 0.99918 3869
## log(Hist3yr)          0.00601 0.99982 4587
## reciprocal_dispersion 0.02692 0.99963 3584
## mean_PPD              0.15472 0.99971 3809
## log-posterior         0.03959 1.00014 1522
##
## For each parameter, mcse is Monte Carlo standard error, n_eff is a crude measure of effective sample
```

```r
#Coding the offset correctly gives different parameter estimates

loo(modelc1)
```

```
##
## Computed from 4000 by 20 log-likelihood matrix.
##
##          Estimate  SE
```

```
## elpd_loo    -85.7 2.0
## p_loo         2.0 0.5
## looic       171.4 3.9
## ------
## MCSE of elpd_loo is 0.0.
## MCSE and ESS estimates assume MCMC draws (r_eff in [0.5, 1.3]).
##
## All Pareto k estimates are good (k < 0.7).
## See help('pareto-k-diagnostic') for details.
```

```
#and different LOO ELPD
```

##### Table 2, Model c2

```
#Code used by Mills et al.
set.seed(98356)
modelc2 <- stan_glm.nb(Incidence ~ Treatment + log(hdyrsrisk) + log(Hist3yr),
                            prior_intercept = normal(0, 1, autoscale = TRUE),
                            prior = normal(0, 1, autoscale = TRUE),
                            prior_aux= cauchy(0, 5, autoscale = TRUE),
                            offset = log(hdyrsrisk),
                            data = rbctconf, refresh=0)

#As with model c1, Mills et al. coded the log(hdyrsrisk) as both an explanatory
#and offset variable

#Correct code is:
modelc2 <- stan_glm.nb(Incidence ~ Treatment + log(Hist3yr),
                     prior_intercept = normal(0, 1, autoscale = TRUE),
                     prior = normal(0, 1, autoscale = TRUE),
                     prior_aux= cauchy(0, 5, autoscale = TRUE),
                     offset = log(hdyrsrisk),
                     data = rbctconf, refresh=0)

#Same coding issue for follow up and post trial periods in code written by Mills et al.

summary(modelc2, digits=5, probs=c(0.025,0.5,0.975))
```

```
##
## Model Info:
##  function:     stan_glm.nb
##  family:       neg_binomial_2 [log]
##  formula:      Incidence ~ Treatment + log(Hist3yr)
##  algorithm:    sampling
##  sample:       4000 (posterior sample size)
##  priors:       see help('prior_summary')
##  observations: 20
##  predictors:   3
##
## Estimates:
##                        mean     sd       2.5%     50%      97.5%
## (Intercept)          -4.94188  0.99170 -6.89377 -4.93955 -2.98628
## TreatmentProactive   -0.10118  0.17091 -0.43150 -0.10169  0.24189
## log(Hist3yr)          0.77455  0.30816  0.17298  0.77370  1.38254
```

```
## reciprocal_dispersion 10.20895  4.62378  4.13211  9.30999 21.33824
##
## Fit Diagnostics:
##            mean     sd       2.5%     50%      97.5%
## mean_PPD 50.71708  6.74583 38.99875 50.05000 65.75125
##
## The mean_ppd is the sample average posterior predictive distribution of the outcome variable (for de
##
## MCMC diagnostics
##                       mcse    Rhat    n_eff
## (Intercept)           0.01578 0.99967 3951
## TreatmentProactive    0.00257 0.99965 4428
## log(Hist3yr)          0.00497 0.99978 3850
## reciprocal_dispersion 0.09323 1.00042 2460
## mean_PPD              0.11331 1.00096 3544
## log-posterior         0.03612 0.99960 1781
##
## For each parameter, mcse is Monte Carlo standard error, n_eff is a crude measure of effective sample
```

#### Table 2, Model d1

```
#Code given by Mills et al. (table 2b)
modeld1 <- stan_glm.nb(Incidence ~ log(hdyrsrisk) + log(Hist3yr),
                  prior_intercept=normal(0,10), prior=normal(0,10),
                  data=rbctconf,
                  diagnostic_file = file.path(tempdir(), "df.csv"),refresh=0)

summary(modeld1, digits=5, probs=c(0.025,0.5,0.975))
```

```
##
## Model Info:
##  function:     stan_glm.nb
##  family:       neg_binomial_2 [log]
##  formula:      Incidence ~ log(hdyrsrisk) + log(Hist3yr)
##  algorithm:    sampling
##  sample:       4000 (posterior sample size)
##  priors:       see help('prior_summary')
##  observations: 20
##  predictors:   3
##
## Estimates:
##                         mean     sd       2.5%     50%      97.5%
## (Intercept)           -2.20207  1.56744 -5.37649 -2.20579  0.83539
## log(hdyrsrisk)         0.49682  0.20109  0.09657  0.49715  0.89260
## log(Hist3yr)           0.87973  0.34927  0.18295  0.88025  1.58121
## reciprocal_dispersion  6.33292  2.09565  2.99697  6.07052 11.11810
##
## Fit Diagnostics:
##            mean     sd       2.5%     50%      97.5%
## mean_PPD 45.29289  7.19427 33.00000 44.55000 61.50000
##
## The mean_ppd is the sample average posterior predictive distribution of the outcome variable (for de
##
```

```
## MCMC diagnostics
##                       mcse    Rhat    n_eff
## (Intercept)           0.02404 0.99992 4253
## log(hdyrsrisk)        0.00316 0.99986 4048
## log(Hist3yr)          0.00570 0.99977 3753
## reciprocal_dispersion 0.03493 1.00022 3600
## mean_PPD              0.11835 1.00029 3695
## log-posterior         0.03913 1.00066 1477
##
## For each parameter, mcse is Monte Carlo standard error, n_eff is a crude measure of effective sample
```

```
#This aligns with the results reported in table 2b of Mills et al,
#and model rs2b in Torgerson et al
```

### Table 2, Model e

```
#Model e, table 2b Mills et al.
set.seed(150499)
modele <- stan_glm(Incidence ~ Treatment + A + B + C + D + E + F + G + H + I
                   +log(Hist3yr),
                              data = rbctconf, family = "poisson",
                              offset = log(Baseline),
                              prior = normal(0, 1),
                              prior_intercept = normal(0, 2),
                              diagnostic_file = file.path(tempdir(), "df.csv"),
                   refresh=0)

summary(modele, digits=5, probs=c(0.025,0.5,0.975))
```

```
##
## Model Info:
##  function:     stan_glm
##  family:       poisson [log]
##  formula:      Incidence ~ Treatment + A + B + C + D + E + F + G + H + I + log(Hist3yr)
##  algorithm:    sampling
##  sample:       4000 (posterior sample size)
##  priors:       see help('prior_summary')
##  observations: 20
##  predictors:   12
##
## Estimates:
##                      mean     sd       2.5%     50%      97.5%
## (Intercept)        -3.38331  0.49811 -4.35441 -3.38224 -2.40610
## TreatmentProactive -0.14608  0.07369 -0.28543 -0.14722 -0.00086
## A1                  0.32704  0.16475  0.00599  0.32555  0.65096
## B1                  0.22008  0.15846 -0.07634  0.21850  0.53463
## C1                  0.24272  0.14712 -0.04292  0.24127  0.53813
## D1                 -0.00472  0.16559 -0.33314 -0.00527  0.31974
## E1                  0.20355  0.15526 -0.08646  0.20204  0.50336
## F1                 -0.39123  0.16497 -0.70897 -0.39535 -0.07014
## G1                 -0.00111  0.14956 -0.29769 -0.00017  0.28983
## H1                 -0.05148  0.17022 -0.37678 -0.05287  0.28566
## I1                 -0.27745  0.17998 -0.64196 -0.27662  0.07900
```

```
## log(Hist3yr)        0.73110  0.15828  0.41891  0.73190  1.03773
##
## Fit Diagnostics:
##            mean     sd       2.5%     50%      97.5%
## mean_PPD 44.14225  2.11312 40.09875 44.15000 48.40125
##
## The mean_ppd is the sample average posterior predictive distribution of the outcome variable (for de
##
## MCMC diagnostics
##                     mcse    Rhat    n_eff
## (Intercept)         0.00991 0.99965 2527
## TreatmentProactive 0.00125 1.00059 3459
## A1                  0.00434 1.00317 1442
## B1                  0.00462 1.00382 1176
## C1                  0.00405 1.00233 1322
## D1                  0.00434 1.00193 1453
## E1                  0.00436 1.00173 1268
## F1                  0.00460 1.00258 1289
## G1                  0.00408 1.00275 1346
## H1                  0.00409 1.00263 1735
## I1                  0.00436 1.00163 1707
## log(Hist3yr)        0.00340 1.00044 2170
## mean_PPD            0.03496 0.99965 3653
## log-posterior       0.06203 0.99968 1575
##
## For each parameter, mcse is Monte Carlo standard error, n_eff is a crude measure of effective sample
```

```r
#The results do not align with the reported results in table 2b Mills et al.

round((exp(coef(modele))[2]-1),3)*100                             # point estimate -13.7%
```

```
## TreatmentProactive
##              -13.7
```

```r
round((exp(posterior_interval(modele, prob = 0.95))[2,1]-1),3)*100 # lower 95% CI
```

```
## [1] -24.8
```

```r
round((exp(posterior_interval(modele, prob = 0.95))[2,2]-1),3)*100 # upper 95% CI
```

```
## [1] -0.1
```

```r
#These results with this model, give the culling effect of 13.6% reduction( CrI 0.1%,24.8%)
#This is considerable less than the effect reported in Mills et al.
#And there is less probability distribution for a culling effect
#Therefore, figure 1 in the main text is incorrect
```

##### Model selection by Bayes Factor

#### Original log linear poisson model Model proposed by Donnelly et al as Bayesian

```r
set.seed(150499)
modelrs <- stan_glm(Incidence ~ Treatment + A + B + C + D + E + F + G + H + I
                    + log(Hist3yr) +log(Baseline), poisson,
```

```r
                   data = rbctconf, prior_intercept=normal(0, 10),
                   diagnostic_file = file.path(tempdir(), "df.csv"),refresh=0)

bridge_1 <- bridge_sampler(modelrs)
```

```
## Iteration: 1
## Iteration: 2
## Iteration: 3
## Iteration: 4
```

```r
modelrs_hdyrsrisk <- stan_glm(Incidence ~ Treatment + A + B + C + D + E + F + G
                              + H + I + log(Hist3yr)+ log(hdyrsrisk), poisson,
                   data = rbctconf, prior_intercept=normal(0, 10),
                   diagnostic_file = file.path(tempdir(), "df.csv"),refresh=0)

bridge_2 <- bridge_sampler(modelrs_hdyrsrisk)
```

```
## Iteration: 1
## Iteration: 2
## Iteration: 3
## Iteration: 4
## Iteration: 5
```

```r
modeld1 <- stan_glm.nb(Incidence ~ log(hdyrsrisk) + log(Hist3yr),
                      prior_intercept=normal(0,10), prior=normal(0,10),
                      data=rbctconf,
                      diagnostic_file = file.path(tempdir(), "df.csv"),refresh=0)

bridge_3 <- bridge_sampler(modeld1)
```

```
## Iteration: 1
## Iteration: 2
## Iteration: 3
## Iteration: 4
```

```r
bf(bridge_3, bridge_1)
```

```
## Estimated Bayes factor in favor of bridge_3 over bridge_1: 75651736.92252
```

```r
#Estimated Bayes factor in favor of bridge_3 over bridge_1: 75962823
#Model without culling effect is supported by a large amount over the original
#log linear poisson model by Donnelly et al.

#compare using the same exposure variable
bf(bridge_3, bridge_2)
```

```
## Estimated Bayes factor in favor of bridge_3 over bridge_2: 48552806.07630
```

```r
#Estimated Bayes factor in favor of bridge_3 over bridge_2: 48949011
#Model with no effect of culling is supported by a large amount over one that has.


model_d2 <- stan_glm.nb(Incidence ~ log(hdyrsrisk) + log(Hist3yr),
                       prior=normal(0, 1),
                       prior_aux = cauchy(0, 5),
                       data=rbctconf,
                       diagnostic_file = file.path(tempdir(), "df.csv"),
```

```r
                      refresh=0)

bridge_4 <- bridge_sampler(model_d2)
```

```
## Iteration: 1
## Iteration: 2
## Iteration: 3
## Iteration: 4
## Iteration: 5
```

```r
bf( bridge_4,bridge_1)
```

```
## Estimated Bayes factor in favor of bridge_4 over bridge_1: 8880524220379.22461
```

```r
#Estimated Bayes factor in favor of bridge_3 over bridge_1: 405.82877

bf( bridge_4,bridge_2)
```

```
## Estimated Bayes factor in favor of bridge_4 over bridge_2: 5699464253802.00293
```

```r
#better comparison as has the same exposure variable: hdyrsrisk
#Estimated Bayes factor in favor of bridge_3 over bridge_2: 367.21850
```

#### Table 3

##### Mills et al, 2024b: paper 2.

```
rbctconf_outside <- read_xlsx("outside_areas_rbct.xlsx", sheet="During_Trial")

#relevel variables so correspond to Mills et al and Nature
rbctconf_outside$Triplet<-relevel(factor(rbctconf_outside$Triplet), ref="J")
rbctconf_outside$Treatment<-relevel(factor(rbctconf_outside$Treatment), ref="Survey-only")
```

##### Table 3, model1: Original Poisson GLM as per Donnelly et al, Nature (2006)

http://dx.doi.org/10.1038/nature04454

```
model1 <- glm(Incidence ~ Treatment + Triplet + log(Hist3yr)
             + log(Baseline),
            family = poisson, data=rbctconf_outside)

summary(model1)
```

```
##
## Call:
## glm(formula = Incidence ~ Treatment + Triplet + log(Hist3yr) +
##     log(Baseline), family = poisson, data = rbctconf_outside)
##
## Coefficients:
##                      Estimate Std. Error z value Pr(>|z|)
## (Intercept)           1.87929    0.98260   1.913  0.05580 .
## TreatmentProactive    0.25310    0.10083   2.510  0.01207 *
## TripletA             -0.27471    0.26640  -1.031  0.30245
## TripletB              0.67142    0.17204   3.903 9.51e-05 ***
## TripletC              0.33289    0.18395   1.810  0.07034 .
## TripletD             -0.33151    0.28116  -1.179  0.23837
## TripletE              0.10470    0.20659   0.507  0.61229
## TripletF              0.12297    0.21294   0.577  0.56362
## TripletG              0.24100    0.20903   1.153  0.24894
## TripletH              0.31090    0.20272   1.534  0.12513
## TripletI             -0.56201    0.26726  -2.103  0.03548 *
## log(Hist3yr)          0.29362    0.09797   2.997  0.00273 **
## log(Baseline)         0.10807    0.22944   0.471  0.63762
## ---
## Signif. codes:  0 '***' 0.001 '**' 0.01 '*' 0.05 '.' 0.1 ' ' 1
##
## (Dispersion parameter for poisson family taken to be 1)
##
##     Null deviance: 108.3610  on 19  degrees of freedom
## Residual deviance:   7.4481  on  7  degrees of freedom
## AIC: 135.36
##
## Number of Fisher Scoring iterations: 4
```

###### Table 3, model1: Estimated effect of culling (95% CI)

```
round((exp(coef(model1))[2] - 1),3)*100   # point estimate
```

```
## TreatmentProactive
##                28.8
```

```r
round((exp(coefci(model1))[2,1]-1),3)*100 # lower 95% CI
```

```
## [1] 5.7
```

```r
round((exp(coefci(model1))[2,2]-1),3)*100 # upper 95% CI
```

```
## [1] 56.9
```

##### Table 3, model1: BIC and AICc

```r
round(BIC(model1),1)
```

```
## [1] 148.3
```

```r
round(AICc(model1),1)
```

```
## [1] 196
```

##### Table 3, model1: LOOCV RMSE

Leave-One-Out Cross-Validation and Root Mean Squared Error, a metric used to evaluate the performance of a model.

```r
rmse <- numeric()
actual_vs_predicted <- data.frame(Actual = numeric(), Predicted = numeric())
for (i in 1:nrow(rbctconf_outside)) {
  # Exclude the ith row
  test_data <- rbctconf_outside[i, ]
  train_data <-rbctconf_outside[-i, ]
  model <- glm(Incidence ~ Treatment + Triplet + log(Hist3yr)
              + log(Baseline), family = poisson,
                   data = train_data)
  predictions <- predict(model, newdata = test_data, type="response")
  rmse[i] <- sqrt(mean((test_data$Incidence - predictions)^2))
  actual_vs_predicted <- rbind(actual_vs_predicted,
                               data.frame(Actual = test_data$Incidence,
                                          Predicted = predictions))
}

rmse1<-actual_vs_predicted %>%
  mutate(rmse = sqrt((Actual - Predicted)^2))

round(mean(rmse1$rmse),2)
```

```
## [1] 8.07
```

##### Table 3, model1: Number of estimated parameters

```r
length(coef(model1))
```

```
## [1] 13
```

**Table 3, model1: Parameter for exposure (95% CI)**

```
# this is on the log scale
round(coef(model1)[13],2)     # point estimate
```

```
## log(Baseline)
##          0.11
```

```
round(coefci(model1)[13,1],2) # lower 95% CI
```

```
## [1] -0.34
```

```
round(coefci(model1)[13,2],2) # upper 95% CI
```

```
## [1] 0.56
```

In the text there is also mentionned this model for other time periods.

```r
# Follow-up period
rbctconf_outside_follow <- read_xlsx("outside_areas_rbct.xlsx",
                                     sheet="Follow_Up")

# relevel
rbctconf_outside_follow$Triplet<-relevel(factor(rbctconf_outside_follow$Triplet),
                                         ref="J")
rbctconf_outside_follow$Treatment<-relevel(factor(rbctconf_outside_follow$Treatment),
                                           ref="Survey-only")

model1_follow <- glm(Incidence ~ Treatment + Triplet + log(Hist3yr)
                     + log(Baseline),
                   family = poisson, data=rbctconf_outside_follow)

summary(model1_follow)
```

```
##
## Call:
## glm(formula = Incidence ~ Treatment + Triplet + log(Hist3yr) +
##     log(Baseline), family = poisson, data = rbctconf_outside_follow)
##
## Coefficients:
##                      Estimate Std. Error z value Pr(>|z|)
## (Intercept)           1.52170    1.07902   1.410  0.15846
## TreatmentProactive    0.19637    0.11896   1.651  0.09879 .
## TripletA             -0.18352    0.32139  -0.571  0.56799
## TripletB              1.06526    0.20689   5.149 2.62e-07 ***
## TripletC              0.67364    0.21721   3.101  0.00193 **
## TripletD             -0.07919    0.32283  -0.245  0.80623
## TripletE              0.40427    0.24354   1.660  0.09692 .
## TripletF              0.20309    0.25920   0.784  0.43332
## TripletG              0.28195    0.25525   1.105  0.26932
## TripletH              0.41385    0.24752   1.672  0.09453 .
## TripletI             -0.37781    0.31496  -1.200  0.23032
## log(Hist3yr)          0.23316    0.11711   1.991  0.04648 *
## log(Baseline)         0.12440    0.25502   0.488  0.62570
## ---
## Signif. codes:  0 '***' 0.001 '**' 0.01 '*' 0.05 '.' 0.1 ' ' 1
##
## (Dispersion parameter for poisson family taken to be 1)
##
##     Null deviance: 116.4616  on 19  degrees of freedom
## Residual deviance:   9.2069  on  7  degrees of freedom
## AIC: 131.37
##
## Number of Fisher Scoring iterations: 4
```

```r
confint(model1_follow)
```

```
## Waiting for profiling to be done...
```

```
##                          2.5 %    97.5 %
## (Intercept)        -0.60876148 3.6270469
## TreatmentProactive -0.03567438 0.4314169
```

```
## TripletA           -0.81563736 0.4472270
## TripletB            0.67187393 1.4856817
## TripletC            0.25630213 1.1110543
## TripletD           -0.71587618 0.5531425
## TripletE           -0.06831512 0.8895939
## TripletF           -0.30714216 0.7136327
## TripletG           -0.21519347 0.7889770
## TripletH           -0.06698282 0.9063126
## TripletI           -1.00541429 0.2346161
## log(Hist3yr)        0.00564154 0.4656558
## log(Baseline)      -0.37497030 0.6260649
```

```
# Post-Trial period
rbctconf_outside_after <- read_xlsx("outside_areas_rbct.xlsx", sheet ="Post_Trial")

# relevel
rbctconf_outside_after$Triplet <- relevel(factor(rbctconf_outside_after$Triplet),
                                        ref="J")
rbctconf_outside_after$Treatment <- relevel(factor(rbctconf_outside_after$Treatment),
                                          ref="Survey-only")

model1_after <- glm(Incidence ~ Treatment + Triplet + log(Hist3yr)
                  + log(Baseline), family = poisson,
                  data=rbctconf_outside_after)

summary(model1_after)
```

```
##
## Call:
## glm(formula = Incidence ~ Treatment + Triplet + log(Hist3yr) +
##     log(Baseline), family = poisson, data = rbctconf_outside_after)
##
## Coefficients:
##                     Estimate Std. Error z value Pr(>|z|)
## (Intercept)         0.283684   0.843415   0.336  0.73661
## TreatmentProactive -0.004161   0.070374  -0.059  0.95285
## TripletA           -0.544971   0.197024  -2.766  0.00567 **
## TripletB           -0.095522   0.121312  -0.787  0.43104
## TripletC           -0.168479   0.120477  -1.398  0.16198
## TripletD           -0.141091   0.199704  -0.707  0.47988
## TripletE           -0.264275   0.143760  -1.838  0.06602 .
## TripletF           -0.612028   0.149462  -4.095 4.22e-05 ***
## TripletG           -0.442045   0.145330  -3.042  0.00235 **
## TripletH           -0.016775   0.140176  -0.120  0.90474
## TripletI           -0.449272   0.181805  -2.471  0.01347 *
## log(Hist3yr)        0.163031   0.066449   2.453  0.01415 *
## log(Baseline)       0.770332   0.189787   4.059 4.93e-05 ***
## ---
## Signif. codes:  0 '***' 0.001 '**' 0.01 '*' 0.05 '.' 0.1 ' ' 1
##
## (Dispersion parameter for poisson family taken to be 1)
##
##     Null deviance: 163.063  on 19  degrees of freedom
## Residual deviance:  25.689  on  7  degrees of freedom
## AIC: 165.33
```

```
##
## Number of Fisher Scoring iterations: 4
```

```
confint(model1_after)
```

```
## Waiting for profiling to be done...

##                          2.5 %      97.5 %
## (Intercept)        -1.38839551  1.92109529
## TreatmentProactive -0.14210842  0.13389510
## TripletA           -0.93459267 -0.16156360
## TripletB           -0.33346215  0.14258542
## TripletC           -0.40527897  0.06750971
## TripletD           -0.53404266  0.24943257
## TripletE           -0.54810258  0.01602872
## TripletF           -0.91043410 -0.32350148
## TripletG           -0.72841578 -0.15821877
## TripletH           -0.29265423  0.25725184
## TripletI           -0.80905575 -0.09558009
## log(Hist3yr)        0.03346172  0.29413312
## log(Baseline)       0.40110035  1.14571269
```

```
# Plausible exposure paramter = 0.77 (CIs 0.40,1.15)
# But no effect of culling
```

##### Table 3, Model 3: Original GLM in generalized Poisson form

```r
#Model 3 (=model 1 in generalized poisson form)
model3 <- glmmTMB(Incidence ~ Treatment + Triplet + log(Hist3yr)
                  + log(Baseline),
                family = genpois, data=rbctconf_outside)
summary(model3)
```

```
##  Family: genpois  ( log )
## Formula:
## Incidence ~ Treatment + Triplet + log(Hist3yr) + log(Baseline)
## Data: rbctconf_outside
##
##      AIC      BIC   logLik deviance df.resid
##    129.8    143.8    -50.9    101.8        6
##
##
## Dispersion parameter for genpois family (): 0.359
##
## Conditional model:
##                      Estimate Std. Error z value Pr(>|z|)
## (Intercept)           1.86271    0.59693   3.120 0.001806 **
## TreatmentProactive    0.25091    0.06048   4.148 3.35e-05 ***
## TripletA             -0.27500    0.16041  -1.714 0.086473 .
## TripletB              0.67389    0.10388   6.487 8.75e-11 ***
## TripletC              0.33942    0.11009   3.083 0.002048 **
## TripletD             -0.32474    0.17036  -1.906 0.056622 .
## TripletE              0.12339    0.12502   0.987 0.323642
## TripletF              0.12370    0.12549   0.986 0.324254
## TripletG              0.24206    0.12486   1.939 0.052542 .
## TripletH              0.32310    0.12499   2.585 0.009737 **
## TripletI             -0.52907    0.15859  -3.336 0.000849 ***
## log(Hist3yr)          0.29308    0.05920   4.951 7.39e-07 ***
## log(Baseline)         0.11075    0.13869   0.799 0.424534
## ---
## Signif. codes:  0 '***' 0.001 '**' 0.01 '*' 0.05 '.' 0.1 ' ' 1
```

```r
confint(model3)
```

```
##                             2.5 %       97.5 %   Estimate
## (Intercept)           0.692746540  3.032669442  1.8627080
## TreatmentProactive    0.132362069  0.369449548  0.2509058
## TripletA             -0.589403537  0.039407038 -0.2749982
## TripletB              0.470282614  0.877492216  0.6738874
## TripletC              0.123655074  0.555183211  0.3394191
## TripletD             -0.658631651  0.009156287 -0.3247377
## TripletE             -0.121635601  0.368416580  0.1233905
## TripletF             -0.122253321  0.369657489  0.1237021
## TripletG             -0.002658888  0.486786464  0.2420638
## TripletH              0.078126480  0.568065462  0.3230960
## TripletI             -0.839896818 -0.218247313 -0.5290721
## log(Hist3yr)          0.177056297  0.409111931  0.2930841
## log(Baseline)        -0.161069459  0.382575519  0.1107530
```

```r
#Implausible exposure parameter = 0.11 (CIs-0.16, 0.38)
exp(confint(model3))
```

```
##                        2.5 %     97.5 %  Estimate
## (Intercept)        1.9991989 20.7525565 6.4411558
## TreatmentProactive 1.1415216  1.4469379 1.2851890
## TripletA           0.5546580  1.0401938 0.7595735
## TripletB           1.6004464  2.4048613 1.9618490
## TripletC           1.1316255  1.7422602 1.4041318
## TripletD           0.5175591  1.0091983 0.7227169
## TripletE           0.8854710  1.4454441 1.1313261
## TripletF           0.8849242  1.4472388 1.1316787
## TripletG           0.9973446  1.6270791 1.2738754
## TripletH           1.0812594  1.7648496 1.3813979
## TripletI           0.4317551  0.8039266 0.5891514
## log(Hist3yr)       1.1936983  1.5054802 1.3405555
## log(Baseline)      0.8512329  1.4660556 1.1171190
```

##### Table 3, model3: Estimated effect of culling (95% CI)

```r
round((exp(confint(model3, level = 0.95)[2,3])-1),3)*100 # point estimate
```

```
## [1] 28.5
```

```r
round((exp(confint(model3, level = 0.95)[2,1])-1),3)*100 # lower 95% CI
```

```
## [1] 14.2
```

```r
round((exp(confint(model3, level = 0.95)[2,2])-1),3)*100 # upper 95% CI
```

```
## [1] 44.7
```

##### Table 3, model1: BIC and AICc

```r
round(BIC(model3),1)
```

```
## [1] 143.8
```

```r
round(AICc(model3),1)
```

```
## [1] 213.8
```

##### Table 3, model3: LOOCV RMSE

Leave-One-Out Cross-Validation and Root Mean Squared Error, a metric used to evaluate the performance of a model.

```r
rmse <- numeric()
actual_vs_predicted <- data.frame(Actual = numeric(), Predicted = numeric())
for (i in 1:nrow(rbctconf_outside)) {
  # Exclude the ith row
  test_data <- rbctconf_outside[i, ]
  train_data <-rbctconf_outside[-i, ]
  model <- glmmTMB(Incidence ~ Treatment + Triplet + log(Hist3yr)
                  + log(Baseline),
                family = genpois,
                   data = train_data)
```

```r
  predictions <- predict(model, newdata = test_data, type="response")
  rmse[i] <- sqrt(mean((test_data$Incidence - predictions)^2))
  actual_vs_predicted <- rbind(actual_vs_predicted,
                               data.frame(Actual = test_data$Incidence,
                                          Predicted = predictions))
}

rmse1<-actual_vs_predicted %>%
  mutate(rmse = sqrt((Actual - Predicted)^2))

round(mean(rmse1$rmse),2)
```

```
## [1] 8.05
```

##### Table 3, model3: Number of estimated parameters

```r
length(fixef(model3)$cond) # plus an additional parameter for overdispersion
```

```
## [1] 13
```

##### Table 3, model3: Parameter for exposure (95% CI)

```r
# this is on the log scale
round(confint(model3, level = 0.95)[13,3],2) # point estimate
```

```
## [1] 0.11
```

```r
round(confint(model3, level = 0.95)[13,1],2) # lower 95% CI
```

```
## [1] -0.16
```

```r
round(confint(model3, level = 0.95)[13,2],2) # upper 95% CI
```

```
## [1] 0.38
```

In the text there is also mentionned this model for other time periods.

```
model3_follow <- glmmTMB(Incidence ~ Treatment + Triplet + log(Hist3yr)
                         + log(Baseline), family = genpois,
                         data=rbctconf_outside_follow)

summary(model3_follow)
```

```
##  Family: genpois  ( log )
## Formula:
## Incidence ~ Treatment + Triplet + log(Hist3yr) + log(Baseline)
## Data: rbctconf_outside_follow
##
##      AIC      BIC   logLik deviance df.resid
##    127.9    141.8    -49.9     99.9        6
##
##
## Dispersion parameter for genpois family (): 0.416
##
## Conditional model:
##                      Estimate Std. Error z value Pr(>|z|)
## (Intercept)           1.49450    0.70217   2.128  0.03330 *
## TreatmentProactive    0.19334    0.07713   2.507  0.01219 *
## TripletA             -0.17295    0.20472  -0.845  0.39823
## TripletB              1.07496    0.13450   7.992 1.33e-15 ***
## TripletC              0.68826    0.14127   4.872 1.11e-06 ***
## TripletD             -0.05326    0.20990  -0.254  0.79970
## TripletE              0.42215    0.15863   2.661  0.00778 **
## TripletF              0.20409    0.16826   1.213  0.22516
## TripletG              0.28652    0.16531   1.733  0.08306 .
## TripletH              0.42515    0.16277   2.612  0.00900 **
## TripletI             -0.29081    0.19930  -1.459  0.14454
## log(Hist3yr)          0.23540    0.07638   3.082  0.00206 **
## log(Baseline)         0.12629    0.16578   0.762  0.44617
## ---
## Signif. codes:  0 '***' 0.001 '**' 0.01 '*' 0.05 '.' 0.1 ' ' 1
```

```
confint(model3_follow)
```

```
##                            2.5 %     97.5 %   Estimate
## (Intercept)           0.11827034 2.87073436  1.4945024
## TreatmentProactive    0.04216680 0.34450889  0.1933378
## TripletA             -0.57419201 0.22829980 -0.1729461
## TripletB              0.81133524 1.33857599  1.0749556
## TripletC              0.41136951 0.96515519  0.6882623
## TripletD             -0.46465907 0.35814186 -0.0532586
## TripletE              0.11125313 0.73305228  0.4221527
## TripletF             -0.12569991 0.53387314  0.2040866
## TripletG             -0.03748627 0.61052084  0.2865173
## TripletH              0.10612693 0.74418245  0.4251547
## TripletI             -0.68143576 0.09982537 -0.2908052
## log(Hist3yr)          0.08569769 0.38509518  0.2353964
## log(Baseline)        -0.19862964 0.45121668  0.1262935
```

```
#Implausible exposure parameter = 0.12 (CIs-0.19, 0.12)
```

```r
model3_after <- glmmTMB(Incidence ~ Treatment + Triplet + log(Hist3yr)
                        + log(Baseline),
                      family = genpois, data=rbctconf_outside_after)
summary(model3_after)
```

```
##  Family: genpois  ( log )
## Formula:
## Incidence ~ Treatment + Triplet + log(Hist3yr) + log(Baseline)
## Data: rbctconf_outside_after
##
##      AIC      BIC   logLik deviance df.resid
##    166.8    180.7    -69.4    138.8        6
##
##
## Dispersion parameter for genpois family (): 1.26
##
## Conditional model:
##                     Estimate Std. Error z value Pr(>|z|)
## (Intercept)         0.291902   0.946804   0.308 0.757852
## TreatmentProactive -0.004212   0.079087  -0.053 0.957525
## TripletA           -0.542957   0.220833  -2.459 0.013945 *
## TripletB           -0.095707   0.136247  -0.702 0.482397
## TripletC           -0.172478   0.135465  -1.273 0.202935
## TripletD           -0.140520   0.223804  -0.628 0.530088
## TripletE           -0.264099   0.161161  -1.639 0.101269
## TripletF           -0.611564   0.167583  -3.649 0.000263 ***
## TripletG           -0.443429   0.163694  -2.709 0.006751 **
## TripletH           -0.016851   0.157471  -0.107 0.914783
## TripletI           -0.473200   0.208676  -2.268 0.023352 *
## log(Hist3yr)        0.162816   0.074757   2.178 0.029411 *
## log(Baseline)       0.769101   0.212818   3.614 0.000302 ***
## ---
## Signif. codes:  0 '***' 0.001 '**' 0.01 '*' 0.05 '.' 0.1 ' ' 1
```

```r
confint(model3_after)
```

```
##                          2.5 %      97.5 %     Estimate
## (Intercept)        -1.56379900  2.14760379  0.291902395
## TreatmentProactive -0.15922084  0.15079641 -0.004212212
## TripletA           -0.97578187 -0.11013175 -0.542956811
## TripletB           -0.36274641  0.17133220 -0.095707105
## TripletC           -0.43798405  0.09302754 -0.172478259
## TripletD           -0.57916828  0.29812796 -0.140520158
## TripletE           -0.57996854  0.05176990 -0.264099320
## TripletF           -0.94002009 -0.28310753 -0.611563812
## TripletG           -0.76426305 -0.12259563 -0.443429342
## TripletH           -0.32548729  0.29178616 -0.016850563
## TripletI           -0.88219779 -0.06420275 -0.473200270
## log(Hist3yr)        0.01629466  0.30933643  0.162815548
## log(Baseline)       0.35198480  1.18621788  0.769101342
```

```r
# Plausible exposure parameter = 0.77 (CIs 0.35, 1.19)
# But no effect of culling
```

##### Table 3, Model 4: Generalized Poisson without culling

```r
model4 <- glmmTMB(Incidence ~ log(Hist3yr) + log(Baseline),
                family = genpois, data=rbctconf_outside)
summary(model4)
```

```
##  Family: genpois  ( log )
## Formula:          Incidence ~ log(Hist3yr) + log(Baseline)
## Data: rbctconf_outside
##
##      AIC      BIC   logLik deviance df.resid
##    150.5    154.5    -71.3    142.5       16
##
##
## Dispersion parameter for genpois family (): 2.94
##
## Conditional model:
##               Estimate Std. Error z value Pr(>|z|)
## (Intercept)    -0.4758     0.9928  -0.479 0.631774
## log(Hist3yr)    0.2088     0.1161   1.798 0.072235 .
## log(Baseline)   0.7334     0.2057   3.566 0.000362 ***
## ---
## Signif. codes:  0 '***' 0.001 '**' 0.01 '*' 0.05 '.' 0.1 ' ' 1
```

```r
confint(model4)
```

```
##                    2.5 %    97.5 %   Estimate
## (Intercept)   -2.4216279 1.4700681 -0.4757799
## log(Hist3yr)  -0.0188528 0.4364055  0.2087763
## log(Baseline)  0.3303486 1.1365001  0.7334244
```

```r
# Plausible exposure parameter = 0.73 (CIs 0.33, 1.14)
# But no effect of culling
```

###### Table 3, model4: BIC and AICc

```r
round(BIC(model4),1)
```

```
## [1] 154.5
```

```r
round(AICc(model4),1)
```

```
## [1] 153.2
```

```r
AICc(model1, model3, model4)
```

```
##        df     AICc
## model1 13 196.0238
## model3 14 213.8349
## model4  4 153.1727
```

```r
#Model 4 has lowest AICc compared to models which report a culling effect
```

###### Table 3, model4: LOOCV RMSE

Leave-One-Out Cross-Validation and Root Mean Squared Error, a metric used to evaluate the performance of a model.

```r
rmse <- numeric()
actual_vs_predicted <- data.frame(Actual = numeric(), Predicted = numeric())
for (i in 1:nrow(rbctconf_outside)) {
  # Exclude the ith row
  test_data <- rbctconf_outside[i, ]
  train_data <-rbctconf_outside[-i, ]
  model <- glmmTMB(Incidence ~ log(Hist3yr) + log(Baseline),
              family = genpois,
                 data = train_data)
  predictions <- predict(model, newdata = test_data, type="response")
  rmse[i] <- sqrt(mean((test_data$Incidence - predictions)^2))
  actual_vs_predicted <- rbind(actual_vs_predicted,
                               data.frame(Actual = test_data$Incidence,
                                          Predicted = predictions))
}

rmse1<-actual_vs_predicted %>%
  mutate(rmse = sqrt((Actual - Predicted)^2))

round(mean(rmse1$rmse),2)
```

```
## [1] 8.26
```

**Table 3, model4: Number of estimated parameters**

```r
length(fixef(model4)$cond) # plus an additional parameter for overdispersion
```

```
## [1] 3
```

**Table 3, model4: Parameter for exposure (95% CI)**

```r
# this is on the log scale
round(confint(model4, level = 0.95)[3,3],2) # point estimate
```

```
## [1] 0.73
```

```r
round(confint(model4, level = 0.95)[3,1],2) # lower 95% CI
```

```
## [1] 0.33
```

```r
round(confint(model4, level = 0.95)[3,2],2) # upper 95% CI
```

```
## [1] 1.14
```

In the text there is also mentionned this model for other time periods.

```
model4_follow <- glmmTMB(Incidence ~ log(Hist3yr) + log(Baseline),
                        family = genpois, data=rbctconf_outside_follow)
summary(model4_follow)
```

```
##  Family: genpois  ( log )
## Formula:          Incidence ~ log(Hist3yr) + log(Baseline)
## Data: rbctconf_outside_follow
##
##      AIC      BIC   logLik deviance df.resid
##    150.2    154.2    -71.1    142.2       16
##
##
## Dispersion parameter for genpois family (): 3.92
##
## Conditional model:
##               Estimate Std. Error z value Pr(>|z|)
## (Intercept)    -0.5656     1.2923  -0.438  0.66164
## log(Hist3yr)    0.1846     0.1434   1.288  0.19787
## log(Baseline)   0.7107     0.2648   2.683  0.00729 **
## ---
## Signif. codes:  0 '***' 0.001 '**' 0.01 '*' 0.05 '.' 0.1 ' ' 1
```

```
confint(model4_follow)
```

```
##                     2.5 %    97.5 %   Estimate
## (Intercept)   -3.09848426 1.9673201 -0.5655821
## log(Hist3yr)  -0.09637678 0.4655536  0.1845884
## log(Baseline)  0.19156700 1.2297479  0.7106574
```

```
# Plausible exposure parameter = 0.71 (CIs 0.19, 1.23)
# But no effect of culling

AICc(model1_follow, model3_follow, model4_follow)
```

```
##               df     AICc
## model1_follow 13 192.0358
## model3_follow 14 211.8985
## model4_follow  4 152.8651
```

```
#Model4_follow has the lowest AICc compared to models which report a culling effect


model4_after <- glmmTMB(Incidence ~ log(Hist3yr) + log(Baseline),
                       family = genpois, data=rbctconf_outside_after)
summary(model4_after)
```

```
##  Family: genpois  ( log )
## Formula:          Incidence ~ log(Hist3yr) + log(Baseline)
## Data: rbctconf_outside_after
##
##      AIC      BIC   logLik deviance df.resid
##    167.3    171.3    -79.7    159.3       16
##
##
## Dispersion parameter for genpois family (): 3.52
```

```
##
## Conditional model:
##                Estimate Std. Error z value Pr(>|z|)
## (Intercept)    -0.05124    0.81374  -0.063   0.9498
## log(Hist3yr)    0.17792    0.08645   2.058   0.0396 *
## log(Baseline)  0.78149    0.16466   4.746 2.08e-06 ***
## ---
## Signif. codes:  0 '***' 0.001 '**' 0.01 '*' 0.05 '.' 0.1 ' ' 1
```

```
confint(model4_after)
```

```
##                      2.5 %    97.5 %    Estimate
## (Intercept)    -1.646148495 1.5436645 -0.05124202
## log(Hist3yr)    0.008480559 0.3473672  0.17792385
## log(Baseline)  0.458753414 1.1042272  0.78149032
```

```
# Plausible exposure parameter = 0.78 (CIs 0.45, 1.04)
# But no effect of culling
AICc(model1_after, model3_after, model4_after)
```

```
##              df     AICc
## model1_after 13 225.9987
## model3_after 14 250.7726
## model4_after  4 169.9757
```

```
#Model4_follow has the lowest AICc compared to models which report a culling effect
```

#### Table 3, Model 5: Generalized Poisson with herd-years-at-risk offset

```
#Model 5
model5 <- glmmTMB(Incidence ~ Treatment + Triplet + log(Hist3yr)
                  + offset(log(hdyrsrisk)),
                family = genpois, data=rbctconf_outside)
summary(model5)
```

```
##  Family: genpois  ( log )
## Formula:
## Incidence ~ Treatment + Triplet + log(Hist3yr) + offset(log(hdyrsrisk))
## Data: rbctconf_outside
##
##      AIC      BIC   logLik deviance df.resid
##      150      163      -62      124        7
##
##
## Dispersion parameter for genpois family (): 1.11
##
## Conditional model:
##                     Estimate Std. Error z value Pr(>|z|)
## (Intercept)         -2.87216    0.28990  -9.907   <2e-16 ***
## TreatmentProactive  0.09493    0.09725   0.976   0.3290
## TripletA           -0.31860    0.22401  -1.422   0.1549
## TripletB           -0.13601    0.18320  -0.742   0.4578
## TripletC           -0.42375    0.19305  -2.195   0.0282 *
## TripletD            0.37845    0.23834   1.588   0.1123
## TripletE           -0.21437    0.20389  -1.051   0.2931
## TripletF           -0.46373    0.21996  -2.108   0.0350 *
## TripletG           -0.55056    0.20557  -2.678   0.0074 **
## TripletH            0.13117    0.19517   0.672   0.5015
## TripletI           -0.07469    0.24811  -0.301   0.7634
## log(Hist3yr)        0.11657    0.09243   1.261   0.2072
## ---
## Signif. codes:  0 '***' 0.001 '**' 0.01 '*' 0.05 '.' 0.1 ' ' 1
```

```
confint(model5)
```

```
##                           2.5 %      97.5 %    Estimate
## (Intercept)        -3.44035613 -2.30396461 -2.87216037
## TreatmentProactive -0.09567147  0.28553545  0.09493199
## TripletA           -0.75764722  0.12044252 -0.31860235
## TripletB           -0.49508367  0.22306450 -0.13600959
## TripletC           -0.80212791 -0.04537625 -0.42375208
## TripletD           -0.08869339  0.84559038  0.37844849
## TripletE           -0.61397666  0.18524451 -0.21436608
## TripletF           -0.89483885 -0.03262336 -0.46373111
## TripletG           -0.95346811 -0.14764592 -0.55055702
## TripletH           -0.25134598  0.51368860  0.13117131
## TripletI           -0.56098972  0.41160205 -0.07469384
## log(Hist3yr)       -0.06459181  0.29773775  0.11657297
```

```
exp(confint(model5))
```

```
##                          2.5 %     97.5 %   Estimate
## (Intercept)        0.03205327 0.09986214 0.05657657
```

```
## TreatmentProactive 0.90876252 1.33047423 1.09958407
## TripletA           0.46876804 1.12799590 0.72716465
## TripletB           0.60951991 1.24990119 0.87283427
## TripletC           0.44837385 0.95563786 0.65458615
## TripletD           0.91512612 2.32935261 1.46001761
## TripletE           0.54119444 1.20351268 0.80705289
## TripletF           0.40867345 0.96790304 0.62893265
## TripletG           0.38540208 0.86273654 0.57662853
## TripletH           0.77775324 1.67144514 1.14016309
## TripletI           0.57064401 1.50923372 0.92802757
## log(Hist3yr)       0.93745005 1.34680854 1.12363950
```

```
#No effect of culling. Treatment parameter =0.09 (CIs -0.10, 0.29)
```

##### Table 3, model5: Estimated effect of culling (95% CI)

```
round((exp(confint(model5, level = 0.95)[2,3])-1),3)*100 # point estimate
```

```
## [1] 10
```

```
round((exp(confint(model5, level = 0.95)[2,1])-1),3)*100 # lower 95% CI
```

```
## [1] -9.1
```

```
round((exp(confint(model5, level = 0.95)[2,2])-1),3)*100 # upper 95% CI
```

```
## [1] 33
```

##### Table 3, model5: BIC and AICc

```
round(BIC(model5),1)
```

```
## [1] 163
```

```
round(AICc(model5),1)
```

```
## [1] 210.7
```

##### Table 3, model5: LOOCV RMSE

Leave-One-Out Cross-Validation and Root Mean Squared Error, a metric used to evaluate the performance of a model.

```
rmse <- numeric()
actual_vs_predicted <- data.frame(Actual = numeric(), Predicted = numeric())
for (i in 1:nrow(rbctconf_outside)) {
  # Exclude the ith row
  test_data <- rbctconf_outside[i, ]
  train_data <-rbctconf_outside[-i, ]
  model <- glmmTMB(Incidence ~ Treatment + Triplet + log(Hist3yr)
                + offset(log(hdyrsrisk)),
              family = genpois,
                 data = train_data)
  predictions <- predict(model, newdata = test_data, type="response")
  rmse[i] <- sqrt(mean((test_data$Incidence - predictions)^2))
  actual_vs_predicted <- rbind(actual_vs_predicted,
                               data.frame(Actual = test_data$Incidence,
```

```r
                                            Predicted = predictions))
}
```

```
## Warning in (function (start, objective, gradient = NULL, hessian = NULL, :
## NA/NaN function evaluation
```

```r
rmse1<-actual_vs_predicted %>%
  mutate(rmse = sqrt((Actual - Predicted)^2))

round(mean(rmse1$rmse),2)
```

```
## [1] 12.08
```

**Table 3, model5: Number of estimated parameters**

```r
length(fixef(model5)$cond) # plus an additional parameter for overdispersion
```

```
## [1] 12
```

In the text there is also mentionned this model for other time periods.

```r
model5_follow <- glmmTMB(Incidence ~ Treatment + Triplet + log(Hist3yr)
                         + offset(log(hdyrsrisk)),
                       family = genpois, data=rbctconf_outside_follow)
summary(model5_follow)
```

```
##  Family: genpois  ( log )
## Formula:
## Incidence ~ Treatment + Triplet + log(Hist3yr) + offset(log(hdyrsrisk))
## Data: rbctconf_outside_follow
##
##      AIC      BIC   logLik deviance df.resid
##    143.0    155.9    -58.5    117.0        7
##
##
## Dispersion parameter for genpois family (): 1.02
##
## Conditional model:
##                     Estimate Std. Error z value Pr(>|z|)
## (Intercept)        -2.772138   0.325792  -8.509   <2e-16 ***
## TreatmentProactive  0.002633   0.105935   0.025   0.9802
## TripletA           -0.018268   0.268766  -0.068   0.9458
## TripletB            0.121350   0.211606   0.573   0.5663
## TripletC           -0.165824   0.219414  -0.756   0.4498
## TripletD            0.467306   0.267572   1.746   0.0807 .
## TripletE           -0.097019   0.232104  -0.418   0.6759
## TripletF           -0.267128   0.256849  -1.040   0.2983
## TripletG           -0.399769   0.242973  -1.645   0.0999 .
## TripletH            0.311718   0.232179   1.343   0.1794
## TripletI            0.202897   0.285919   0.710   0.4779
## log(Hist3yr)        0.037015   0.104309   0.355   0.7227
## ---
## Signif. codes:  0 '***' 0.001 '**' 0.01 '*' 0.05 '.' 0.1 ' ' 1
```

```r
confint(model5_follow)
```

```
##                          2.5 %      97.5 %     Estimate
## (Intercept)        -3.41067898 -2.13359759 -2.772138289
## TreatmentProactive -0.20499647  0.21026302  0.002633275
## TripletA           -0.54504024  0.50850377 -0.018268234
## TripletB           -0.29339016  0.53608972  0.121349779
## TripletC           -0.59586791  0.26421963 -0.165824141
## TripletD           -0.05712488  0.99173636  0.467305744
## TripletE           -0.55193520  0.35789722 -0.097018992
## TripletF           -0.77054330  0.23628811 -0.267127595
## TripletG           -0.87598734  0.07645005 -0.399768644
## TripletH           -0.14334487  0.76678186  0.311718494
## TripletI           -0.35749412  0.76328753  0.202896706
## log(Hist3yr)       -0.16742647  0.24145649  0.037015008
```

```r
#No effect of culling. Treatment parameter =0.002 (CIs -0.20, 0.21)

model5_after <- glmmTMB(Incidence ~ Treatment + Triplet + log(Hist3yr)
                        + offset(log(hdyrsrisk)),
                      family = genpois, data=rbctconf_outside_after)
```

```r
summary(model5_after)
```

```
##  Family: genpois  ( log )
## Formula:
## Incidence ~ Treatment + Triplet + log(Hist3yr) + offset(log(hdyrsrisk))
## Data: rbctconf_outside_after
##
##      AIC      BIC   logLik deviance df.resid
##    165.9    178.9    -70.0    139.9        7
##
##
## Dispersion parameter for genpois family (): 1.33
##
## Conditional model:
##                     Estimate Std. Error z value Pr(>|z|)
## (Intercept)         -2.58470    0.20917 -12.357  < 2e-16 ***
## TreatmentProactive -0.03058    0.07758  -0.394 0.693434
## TripletA           -0.39770    0.18039  -2.205 0.027478 *
## TripletB           -0.09058    0.14041  -0.645 0.518875
## TripletC           -0.18646    0.13856  -1.346 0.178413
## TripletD            0.02754    0.16630   0.166 0.868486
## TripletE           -0.19600    0.15248  -1.285 0.198652
## TripletF           -0.61268    0.17218  -3.558 0.000373 ***
## TripletG           -0.52655    0.14927  -3.528 0.000419 ***
## TripletH            0.06028    0.14384   0.419 0.675188
## TripletI           -0.35243    0.17827  -1.977 0.048046 *
## log(Hist3yr)        0.13297    0.07224   1.841 0.065683 .
## ---
## Signif. codes:  0 '***' 0.001 '**' 0.01 '*' 0.05 '.' 0.1 ' ' 1
```

```r
confint(model5_after)
```

```
##                            2.5 %       97.5 %    Estimate
## (Intercept)         -2.994659629 -2.174739723 -2.58469968
## TreatmentProactive -0.182631626  0.121469120 -0.03058125
## TripletA           -0.751264298 -0.044141772 -0.39770303
## TripletB           -0.365771356  0.184621313 -0.09057502
## TripletC           -0.458035894  0.085119777 -0.18645806
## TripletD           -0.298404777  0.353477264  0.02753624
## TripletE           -0.494865014  0.102859482 -0.19600277
## TripletF           -0.950147721 -0.275207390 -0.61267756
## TripletG           -0.819109117 -0.233986557 -0.52654784
## TripletH           -0.221651329  0.342202596  0.06027563
## TripletI           -0.701832856 -0.003029411 -0.35243113
## log(Hist3yr)       -0.008624583  0.274564805  0.13297011
```

```r
#No effect of culling. Treatment parameter =-0.03 (CIs -0.18, 0.12)
```

##### Table 3, Model 6: Generalized Poisson with herd-years-at risk offset and without a culling effect

```
model6 <- glmmTMB(Incidence ~ log(Hist3yr) + offset(log(hdyrsrisk)),
                family = genpois, data=rbctconf_outside)

summary(model6)
```

```
##  Family: genpois  ( log )
## Formula:          Incidence ~ log(Hist3yr) + offset(log(hdyrsrisk))
## Data: rbctconf_outside
##
##      AIC      BIC   logLik deviance df.resid
##    148.9    151.9    -71.5    142.9       17
##
##
## Dispersion parameter for genpois family (): 2.72
##
## Conditional model:
##              Estimate Std. Error z value Pr(>|z|)
## (Intercept)   -3.1631     0.2947 -10.734   <2e-16 ***
## log(Hist3yr)   0.1661     0.1134   1.464    0.143
## ---
## Signif. codes:  0 '***' 0.001 '**' 0.01 '*' 0.05 '.' 0.1 ' ' 1
```

###### Table 3, model6: BIC and AICc

```
round(BIC(model6),1)
```

```
## [1] 151.9
```

```
round(AICc(model6),1)
```

```
## [1] 150.4
```

```
AICc(model5, model6)
```

```
##        df     AICc
## model5 13 210.7029
## model6  3 150.4455
```

```
#model 6, no culling effect has much lower AICc than model 5 with culling effect as an explanatory vari
```

###### Table 3, model6: LOOCV RMSE

Leave-One-Out Cross-Validation and Root Mean Squared Error, a metric used to evaluate the performance of a model.

```
rmse <- numeric()
actual_vs_predicted <- data.frame(Actual = numeric(), Predicted = numeric())
for (i in 1:nrow(rbctconf_outside)) {
  # Exclude the ith row
  test_data <- rbctconf_outside[i, ]
  train_data <-rbctconf_outside[-i, ]
  model <- glmmTMB(Incidence ~ log(Hist3yr) + offset(log(hdyrsrisk)),
                family = genpois,
                   data = train_data)
```

```
  predictions <- predict(model, newdata = test_data, type="response")
  rmse[i] <- sqrt(mean((test_data$Incidence - predictions)^2))
  actual_vs_predicted <- rbind(actual_vs_predicted,
                              data.frame(Actual = test_data$Incidence,
                                         Predicted = predictions))
}

rmse1<-actual_vs_predicted %>%
  mutate(rmse = sqrt((Actual - Predicted)^2))

round(mean(rmse1$rmse),2)
```

```
## [1] 8
```

###### Table 3, model6: Number of estimated parameters

```
length(fixef(model6)$cond) # plus an additional parameter for overdispersion
```

```
## [1] 2
```

In the text there is also mentionned this model for other time periods.

```
model6_follow <- glmmTMB(Incidence ~ log(Hist3yr) + offset(log(hdyrsrisk)),
                       family = genpois, data=rbctconf_outside_follow)

#No effect of culling
AICc(model5_follow, model6_follow)
```

```
##               df     AICc
## model5_follow 13 203.6290
## model6_follow  3 138.2101
```

```
#model 6_follow, no culling effect has much lower AICc than
#model 5_follow with culling effect as explanatory variable.

model6_after<-glmmTMB(Incidence~log(Hist3yr)+offset(log(hdyrsrisk)),
                      family = genpois, data=rbctconf_outside_after)
#No effect of culling
AICc(model5_after, model6_after)
```

```
##              df     AICc
## model5_after 13 226.5731
## model6_after  3 168.5204
```

```
#model 6_follow, no culling effect has much lower AICc than model 5_follow
#with culling effect as explanatory variable.
```

##### Table 3, Model 8: Generalized Poisson with herd-years-at-risk covariate

```
#model 8. As model 6 without offset
model8 <- glmmTMB(Incidence ~ log(Hist3yr) + log(hdyrsrisk),
                 family = genpois, data=rbctconf_outside)

summary(model8)
```

```
##  Family: genpois  ( log )
## Formula:          Incidence ~ log(Hist3yr) + log(hdyrsrisk)
## Data: rbctconf_outside
##
##      AIC      BIC   logLik deviance df.resid
##    141.9    145.8    -66.9    133.9       16
##
##
## Dispersion parameter for genpois family (): 1.89
##
## Conditional model:
##                Estimate Std. Error z value Pr(>|z|)
## (Intercept)    -0.81732    0.73182  -1.117   0.2641
## log(Hist3yr)    0.18563    0.09263   2.004   0.0451 *
## log(hdyrsrisk)  0.61274    0.11468   5.343 9.14e-08 ***
## ---
## Signif. codes:  0 '***' 0.001 '**' 0.01 '*' 0.05 '.' 0.1 ' ' 1
```

```
confint(model8)
```

```
##                       2.5 %    97.5 %   Estimate
## (Intercept)    -2.251671126 0.6170231 -0.8173240
## log(Hist3yr)    0.004085174 0.3671810  0.1856331
## log(hdyrsrisk)  0.387967399 0.8375081  0.6127378
```

```
#Plausible parameter for exposure parameter 0.61 (CIs 0.39, 0.84)
```

###### Table 3, model8: BIC and AICc

```
round(BIC(model8),1)
```

```
## [1] 145.8
```

```
round(AICc(model8),1)
```

```
## [1] 144.5
```

```
AICc(model5, model6, model8)
```

```
##        df     AICc
## model5 13 210.7029
## model6  3 150.4455
## model8  4 144.5290
```

```
#No effect of culling
#model 8, now has the lowest AICc

BIC(model1,model8)
```

```
##        df      BIC
```

```
## model1 13 148.3017
## model8  4 145.8453
```

```
#lower BIC than model 1, the original model in Nature.
```

##### Table 3, model8: LOOCV RMSE

Leave-One-Out Cross-Validation and Root Mean Squared Error, a metric used to evaluate the performance of a model.

```
rmse <- numeric()
actual_vs_predicted <- data.frame(Actual = numeric(), Predicted = numeric())
for (i in 1:nrow(rbctconf_outside)) {
  # Exclude the ith row
  test_data <- rbctconf_outside[i, ]
  train_data <-rbctconf_outside[-i, ]
  model <- glmmTMB(Incidence ~ log(Hist3yr) + log(hdyrsrisk),
                family = genpois,
                   data = train_data)
  predictions <- predict(model, newdata = test_data, type="response")
  rmse[i] <- sqrt(mean((test_data$Incidence - predictions)^2))
  actual_vs_predicted <- rbind(actual_vs_predicted,
                               data.frame(Actual = test_data$Incidence,
                                          Predicted = predictions))
}

rmse1<-actual_vs_predicted %>%
  mutate(rmse = sqrt((Actual - Predicted)^2))

round(mean(rmse1$rmse),2)
```

```
## [1] 6.86
```

##### Table 3, model8: Number of estimated parameters

```
length(fixef(model8)$cond) # plus an additional parameter for overdispersion
```

```
## [1] 3
```

##### Table 3, model8: Parameter for exposure (95% CI)

```
#herd years at risk
# this is on the log scale
round(confint(model8, level = 0.95)[3,3],2) # point estimate
```

```
## [1] 0.61
```

```
round(confint(model8, level = 0.95)[3,1],2) # lower 95% CI
```

```
## [1] 0.39
```

```
round(confint(model8, level = 0.95)[3,2],2) # upper 95% CI
```

```
## [1] 0.84
```

In the text there is also mentioned this model for other time periods.

```
#model 8. As model 6 without offset
model8_follow <- glmmTMB(Incidence ~ log(Hist3yr) + log(hdyrsrisk),
                        family = genpois, data=rbctconf_outside_follow)
confint(model8_follow)
```

```
##                       2.5 %     97.5 %   Estimate
## (Intercept)    -2.96116123 -0.1725968 -1.5668790
## log(Hist3yr)   -0.03395898  0.3737313  0.1698862
## log(hdyrsrisk)  0.51222152  0.9615697  0.7368956
```

```
#Plausible parameter for exposure parameter 0.74 (CIs 0.51, 0.96)
#No effect of culling

AICc(model5_follow, model6_follow, model8_follow)
```

```
##               df     AICc
## model5_follow 13 203.6290
## model6_follow  3 138.2101
## model8_follow  4 136.6551
```

```
#model8_follow, now has the lowest AICc
BIC(model1_follow,model8_follow)
```

```
##               df      BIC
## model1_follow 13 144.3136
## model8_follow  4 137.9714
```

```
#lower BIC than model1_follow.

#model 8. As model 6 without offset
model8_after <- glmmTMB(Incidence ~ log(Hist3yr) + log(hdyrsrisk),
                       family = genpois, data=rbctconf_outside_after)
confint(model8_after)
```

```
##                        2.5 %    97.5 %   Estimate
## (Intercept)    -3.696333509 0.6682665 -1.5140335
## log(Hist3yr)    0.008480891 0.3473665  0.1779237
## log(hdyrsrisk)  0.458763041 1.1042166  0.7814898
```

```
#Plausible parameter for exposure parameter 0.78 (CIs 0.45, 1.10)
#No effect of culling
AICc(model5_after, model6_after, model8_after)
```

```
##              df     AICc
## model5_after 13 226.5731
## model6_after  3 168.5204
## model8_after  4 169.9757
```

```
#model6_after, still has the lowest AICc
BIC(model1_after,model8_after)
```

```
##              df      BIC
## model1_after 13 178.2765
## model8_after  4 171.2919
```

```
#lower BIC than model1_after.
```

#### Table 3, Model 11: Generalized poisson with no predictors

```
model11 <- glmmTMB(Incidence ~ 1, family = genpois, data=rbctconf_outside)
```

##### Table 3, model11: BIC and AICc

```
round(BIC(model11),1)
```

```
## [1] 161.1
```

```
round(AICc(model11),1)
```

```
## [1] 159.9
```

```
AICc(model1, model11)
```

```
##         df     AICc
## model1  13 196.0238
## model11  2 159.8567
```

```
#model 11 with no predictors has lower AICc than model 1
```

##### Table 3, model11: LOOCV RMSE

Leave-One-Out Cross-Validation and Root Mean Squared Error, a metric used to evaluate the performance of a model.

```
rmse <- numeric()
actual_vs_predicted <- data.frame(Actual = numeric(), Predicted = numeric())
for (i in 1:nrow(rbctconf_outside)) {
  # Exclude the ith row
  test_data <- rbctconf_outside[i, ]
  train_data <-rbctconf_outside[-i, ]
  model <- glmmTMB(Incidence ~ 1,
              family = genpois,
                 data = train_data)
  predictions <- predict(model, newdata = test_data, type="response")
  rmse[i] <- sqrt(mean((test_data$Incidence - predictions)^2))
  actual_vs_predicted <- rbind(actual_vs_predicted,
                               data.frame(Actual = test_data$Incidence,
                                          Predicted = predictions))
}

rmse1<-actual_vs_predicted %>%
  mutate(rmse = sqrt((Actual - Predicted)^2))

round(mean(rmse1$rmse),2)
```

```
## [1] 10.11
```

##### Table 3, model11: Number of estimated parameters

```
length(fixef(model11)$cond) # plus an additional parameter for overdispersion
```

```
## [1] 1
```

In the text there is also mentionned this model for other time periods.

```
model11_follow <- glmmTMB(Incidence ~ 1, family = genpois,
                          data=rbctconf_outside_follow)

AICc(model1_follow, model11_follow)
```

```
##                df     AICc
## model1_follow  13 192.0358
## model11_follow  2 154.4439
```

```
#model 11_follow with no predictors has lower AICc than model 1_follow

model11_after <- glmmTMB(Incidence ~ 1, family = genpois,
                         data=rbctconf_outside_after)
AICc(model1_after, model11_after)
```

```
##               df     AICc
## model1_after  13 225.9987
## model11_after  2 180.7595
```

```
#model 11_after with no predictors has lower AICc than model 1_after
```

### Bayesian Analysis Table 4

#### Table 4, Model a.1: Varying intercepts for triplets and covariates of culling efefct,

historical 3-year incidence and baseline herds at risk.

```
#Code in Mills et al, purported to be the same as the code for rs in Torgerson et al:
set.seed(125347599)
model_a1 <- stan_glm.nb(Incidence ~ Treatment + A + B + C + D + E + F +G + H
                        + I +log(Hist3yr) + log(Baseline),
                  data = rbctconf_outside, prior_intercept=normal(0, 10),
                  diagnostic_file = file.path(tempdir(), "df.csv"),refresh=0)

prior_summary(model_a1, digits = 2)
```

```
## Priors for model 'model_a1'
## ------
## Intercept (after predictors centered)
##  ~ normal(location = 0, scale = 10)
##
## Coefficients
##   Specified prior:
##     ~ normal(location = [0,0,0,...], scale = [2.5,2.5,2.5,...])
##   Adjusted prior:
##     ~ normal(location = [0,0,0,...], scale = [4.87,8.12,8.12,...])
##
## Auxiliary (reciprocal_dispersion)
##  ~ exponential(rate = 1)
## ------
## See help('prior_summary.stanreg') for more details
```

```
#Auxiliary (reciprocal_dispersion)
#~ exponential(rate = 1)
summary(model_a1, digits=3, probs=c(0.025,0.5,0.975))
```

```
##
## Model Info:
##  function:     stan_glm.nb
##  family:       neg_binomial_2 [log]
##  formula:      Incidence ~ Treatment + A + B + C + D + E + F + G + H + I + log(Hist3yr) +
##     log(Baseline)
##  algorithm:    sampling
##  sample:       4000 (posterior sample size)
##  priors:       see help('prior_summary')
##  observations: 20
##  predictors:   13
##
## Estimates:
##                         mean   sd     2.5%   50%    97.5%
## (Intercept)            1.781  3.832 -6.005  1.777  9.548
## TreatmentProactive     0.269  0.324 -0.392  0.272  0.905
## A1                    -0.270  0.941 -2.117 -0.292  1.632
## B1                     0.679  0.663 -0.659  0.691  1.990
## C1                     0.303  0.633 -0.980  0.302  1.539
## D1                    -0.321  0.948 -2.134 -0.344  1.581
## E1                     0.100  0.723 -1.356  0.109  1.481
```

```
## F1                      0.090  0.653 -1.245  0.087  1.382
## G1                      0.207  0.676 -1.179  0.225  1.524
## H1                      0.289  0.755 -1.242  0.297  1.804
## I1                     -0.601  0.846 -2.295 -0.608  1.077
## log(Hist3yr)            0.288  0.311 -0.331  0.294  0.885
## log(Baseline)           0.154  0.893 -1.598  0.144  2.016
## reciprocal_dispersion  3.850  1.716  1.339  3.582  7.758
##
## Fit Diagnostics:
##            mean   sd     2.5%   50%    97.5%
## mean_PPD 36.642 16.770 22.900 33.675 67.151
##
## The mean_ppd is the sample average posterior predictive distribution of the outcome variable (for de
##
## MCMC diagnostics
##                       mcse  Rhat  n_eff
## (Intercept)           0.115 1.007 1113
## TreatmentProactive    0.007 1.001 2108
## A1                    0.030 1.008  993
## B1                    0.019 1.005 1178
## C1                    0.017 1.002 1330
## D1                    0.029 1.008 1071
## E1                    0.022 1.006 1126
## F1                    0.018 1.003 1307
## G1                    0.017 1.001 1500
## H1                    0.024 1.008 1029
## I1                    0.027 1.007  982
## log(Hist3yr)          0.008 1.003 1374
## log(Baseline)         0.027 1.006 1095
## reciprocal_dispersion 0.049 1.002 1229
## mean_PPD              0.457 1.002 1348
## log-posterior         0.157 1.002  654
##
## For each parameter, mcse is Monte Carlo standard error, n_eff is a crude measure of effective sample
```

```r
#reciprocal_dispersion = 3.9
```

##### Table 4, model_a1: Estimated effect of culling (95% CI)

```r
round((exp(coef(model_a1))[2]-1),3)*100 # point estimate
```

```
## TreatmentProactive
##               31.2
```

```r
round((exp(posterior_interval(model_a1, prob = 0.95))[2,1]-1),3)*100
```

```
## [1] -32.4
```

```r
round((exp(posterior_interval(model_a1, prob = 0.95))[2,2]-1),3)*100
```

```
## [1] 147.2
```

##### Table 4, model_a1: LOO

```r
loo(model_a1, k_threshold = 0.7)
```

```
##
## Computed from 4000 by 20 log-likelihood matrix.
##
##          Estimate  SE
## elpd_loo    -85.6 2.2
## p_loo         7.6 0.6
## looic       171.1 4.4
## ------
## MCSE of elpd_loo is 0.2.
## MCSE and ESS estimates assume MCMC draws (r_eff in [0.3, 0.6]).
##
## All Pareto k estimates are good (k < 0.7).
## See help('pareto-k-diagnostic') for details.
```

#### Table 4, model rs: Actual code for model rs in Torgerson et al.

```
set.seed(93846)
rs <- stan_glm(Incidence~Treatment+A+B+C+D+E+F+G+H+I+log(Hist3yr)+log(Baseline),
                        data = rbctconf_outside, prior_intercept=normal(0, 10), poisson,
                        diagnostic_file = file.path(tempdir(), "df.csv"),refresh=0)

#This is effectively model 1, table 3 in Bayesian form. model_a1 forced
#overdispersion by using negative binomial regression and forcing a low reciprocal dispersion parameter

summary(rs, digits=3, probs=c(0.025,0.5,0.975))
```

```
##
## Model Info:
##  function:     stan_glm
##  family:       poisson [log]
##  formula:      Incidence ~ Treatment + A + B + C + D + E + F + G + H + I + log(Hist3yr) +
##     log(Baseline)
##  algorithm:    sampling
##  sample:       4000 (posterior sample size)
##  priors:       see help('prior_summary')
##  observations: 20
##  predictors:   13
##
## Estimates:
##                       mean   sd     2.5%   50%    97.5%
## (Intercept)          1.862  0.970 -0.010  1.863  3.814
## TreatmentProactive  0.255  0.099  0.062  0.254  0.454
## A1                 -0.279  0.270 -0.795 -0.285  0.269
## B1                  0.674  0.174  0.344  0.672  1.025
## C1                  0.340  0.183 -0.020  0.336  0.704
## D1                 -0.335  0.283 -0.892 -0.332  0.224
## E1                  0.109  0.212 -0.298  0.105  0.534
## F1                  0.122  0.210 -0.293  0.125  0.536
## G1                  0.247  0.205 -0.153  0.244  0.661
## H1                  0.314  0.204 -0.089  0.313  0.726
## I1                 -0.563  0.265 -1.081 -0.557 -0.036
## log(Hist3yr)        0.298  0.099  0.110  0.298  0.498
## log(Baseline)       0.106  0.226 -0.336  0.103  0.553
##
## Fit Diagnostics:
##            mean   sd     2.5%   50%    97.5%
## mean_PPD 28.532  1.665 25.300 28.500 31.900
##
## The mean_ppd is the sample average posterior predictive distribution of the outcome variable (for de
##
## MCMC diagnostics
##                    mcse  Rhat  n_eff
## (Intercept)        0.024 1.001 1691
## TreatmentProactive 0.002 1.000 2150
## A1                 0.007 1.001 1368
## B1                 0.004 1.000 1498
## C1                 0.004 0.999 1743
## D1                 0.008 1.000 1413
```

```
## E1                 0.006 1.001 1435
## F1                 0.005 1.000 1965
## G1                 0.005 1.000 2069
## H1                 0.006 1.000 1359
## I1                 0.007 1.000 1496
## log(Hist3yr)       0.002 1.000 2158
## log(Baseline)      0.006 1.002 1571
## mean_PPD           0.026 0.999 4038
## log-posterior      0.059 1.002 1742
##
## For each parameter, mcse is Monte Carlo standard error, n_eff is a crude measure of effective sample
```

##### Table 4, model rs: Estimated effect of culling (95% CI)

```
round((exp(coef(rs))[2]-1),3)*100    # point estimate
```

```
## TreatmentProactive
##               28.9
```

```
round((exp(posterior_interval(rs, prob = 0.95))[2,1]-1),3)*100 # lower 95% CI
```

```
## [1] 6.4
```

```
round((exp(posterior_interval(rs, prob = 0.95))[2,2]-1),3)*100 # upper 95% CI
```

```
## [1] 57.5
```

##### Table 4, model rs: LOO

```
loo(rs, k_threshold = 0.7)
```

```
##
## Computed from 4000 by 20 log-likelihood matrix.
##
##          Estimate  SE
## elpd_loo    -71.0 2.1
## p_loo        12.8 1.3
## looic       142.0 4.2
## ------
## MCSE of elpd_loo is 0.3.
## MCSE and ESS estimates assume MCMC draws (r_eff in [0.4, 1.4]).
##
## All Pareto k estimates are good (k < 0.7).
## See help('pareto-k-diagnostic') for details.
```

##### Table 4, model_a2

```
set.seed(6754)
model_a2 <- stan_glm.nb(Incidence~Treatment+A+B+C+D+E+F+G+H+I+log(Hist3yr)+log(Baseline),
                         data = rbctconf_outside, prior_intercept=normal(0, 1, autoscale=TRUE),
                         prior_aux= cauchy(0, 5),
                         prior = normal(0, 1),
                         diagnostic_file = file.path(tempdir(), "df.csv"),refresh=0)

prior_summary(model_a2, digits = 2)
```

```
## Priors for model 'model_a2'
## ------
## Intercept (after predictors centered)
##  ~ normal(location = 0, scale = 1)
##
## Coefficients
##  ~ normal(location = [0,0,0,...], scale = [1,1,1,...])
##
## Auxiliary (reciprocal_dispersion)
##  ~ half-cauchy(location = 0, scale = 5)
## ------
## See help('prior_summary.stanreg') for more details
```

```
#Auxiliary (reciprocal_dispersion)
#~ half-cauchy(location = 0, scale = 5)

summary(model_a2, digits=3, probs=c(0.025,0.5,0.975))
```

```
##
## Model Info:
##  function:     stan_glm.nb
##  family:       neg_binomial_2 [log]
##  formula:      Incidence ~ Treatment + A + B + C + D + E + F + G + H + I + log(Hist3yr) +
##     log(Baseline)
##  algorithm:    sampling
##  sample:       4000 (posterior sample size)
##  priors:       see help('prior_summary')
##  observations: 20
##  predictors:   13
##
## Estimates:
##                       mean   sd     2.5%   50%    97.5%
## (Intercept)            1.588  1.336 -1.119  1.578  4.258
## TreatmentProactive     0.243  0.137 -0.029  0.241  0.517
## A1                    -0.248  0.325 -0.873 -0.255  0.404
## B1                     0.638  0.242  0.144  0.643  1.101
## C1                     0.294  0.251 -0.222  0.296  0.789
## D1                    -0.300  0.345 -0.993 -0.303  0.380
## E1                     0.103  0.264 -0.419  0.106  0.624
## F1                     0.074  0.267 -0.467  0.079  0.577
## G1                     0.189  0.266 -0.334  0.192  0.698
## H1                     0.291  0.271 -0.252  0.295  0.814
## I1                    -0.541  0.329 -1.204 -0.547  0.094
## log(Hist3yr)           0.281  0.128  0.025  0.282  0.528
```

```
## log(Baseline)            0.182    0.309   -0.433    0.181    0.804
## reciprocal_dispersion  158.933 2265.644    6.703   37.006  847.301
##
## Fit Diagnostics:
##            mean   sd     2.5%   50%    97.5%
## mean_PPD 28.805  2.676 24.199 28.650 34.800
##
## The mean_ppd is the sample average posterior predictive distribution of the outcome variable (for de
##
## MCMC diagnostics
##                       mcse   Rhat   n_eff
## (Intercept)            0.035  1.001 1492
## TreatmentProactive     0.003  1.001 2303
## A1                     0.009  1.004 1209
## B1                     0.006  1.002 1634
## C1                     0.006  1.001 1811
## D1                     0.010  1.002 1294
## E1                     0.007  1.002 1475
## F1                     0.006  1.002 2030
## G1                     0.006  0.999 2340
## H1                     0.008  1.002 1275
## I1                     0.009  1.004 1395
## log(Hist3yr)           0.003  1.002 2020
## log(Baseline)          0.008  1.001 1367
## reciprocal_dispersion 37.636  1.000 3624
## mean_PPD               0.044  1.000 3739
## log-posterior          0.114  1.001 1029
##
## For each parameter, mcse is Monte Carlo standard error, n_eff is a crude measure of effective sample
```

```
#reciprocal_dispersion = 194.1
# Note as reciprocal dispersion increases in trends towards a poisson model.
#When reciprocal_dispersion = infinity, it is poisson
#With such a high reciprocal dispersion, this is close to poisson, ie Model 1 in Bayesian form.
```

##### Table 4, model_a2: Estimated effect of culling (95% CI)

```
round((exp(coef(model_a2))[2]-1),3)*100    # point estimate
```

```
## TreatmentProactive
##               27.3
```

```
round((exp(posterior_interval(model_a2, prob = 0.95))[2,1]-1),3)*100 # lower 95% CI
```

```
## [1] -2.9
```

```
round((exp(posterior_interval(model_a2, prob = 0.95))[2,2]-1),3)*100 # upper 95% CI
```

```
## [1] 67.7
```

##### Table 4, model_a2: LOO

```
loo(model_a2)
```

```
## Warning: Found 8 observation(s) with a pareto_k > 0.7. We recommend calling 'loo' again with argumen
```

```
##
## Computed from 4000 by 20 log-likelihood matrix.
##
##          Estimate  SE
## elpd_loo    -71.9 1.8
## p_loo         8.7 0.7
## looic       143.7 3.5
## ------
## MCSE of elpd_loo is NA.
## MCSE and ESS estimates assume MCMC draws (r_eff in [0.4, 1.3]).
##
## Pareto k diagnostic values:
##                          Count Pct.    Min. ESS
## (-Inf, 0.7]   (good)     12    60.0%   278
##    (0.7, 1]   (bad)       8    40.0%   <NA>
##    (1, Inf)   (very bad)  0     0.0%   <NA>
## See help('pareto-k-diagnostic') for details.
```

##### Table 4, Model b1

```r
set.seed(176254)
model_b1 <- stan_glm.nb(Incidence~Treatment+log(Hist3yr)+log(Baseline),
                        data = rbctconf_outside, prior_intercept=normal(0, 10),
                        diagnostic_file = file.path(tempdir(), "df.csv"),refresh=0)
summary(model_b1, digits=3, probs=c(0.025,0.5,0.975))
```

```
##
## Model Info:
##  function:     stan_glm.nb
##  family:       neg_binomial_2 [log]
##  formula:      Incidence ~ Treatment + log(Hist3yr) + log(Baseline)
##  algorithm:    sampling
##  sample:       4000 (posterior sample size)
##  priors:       see help('prior_summary')
##  observations: 20
##  predictors:   4
##
## Estimates:
##                          mean   sd     2.5%   50%    97.5%
## (Intercept)            -0.358  1.505 -3.291 -0.371  2.635
## TreatmentProactive      0.210  0.241 -0.271  0.212  0.689
## log(Hist3yr)            0.264  0.181 -0.115  0.270  0.597
## log(Baseline)           0.657  0.330  0.002  0.651  1.298
## reciprocal_dispersion   4.844  1.672  2.152  4.636  8.617
##
## Fit Diagnostics:
##            mean   sd     2.5%   50%    97.5%
## mean_PPD 29.673  5.300 20.649 29.150 41.351
##
## The mean_ppd is the sample average posterior predictive distribution of the outcome variable (for de
##
## MCMC diagnostics
##                       mcse  Rhat  n_eff
## (Intercept)           0.024 1.000 3776
## TreatmentProactive    0.004 1.001 3865
## log(Hist3yr)          0.003 1.000 3239
## log(Baseline)         0.006 1.000 3509
## reciprocal_dispersion 0.028 1.001 3680
## mean_PPD              0.087 1.000 3740
## log-posterior         0.047 1.001 1353
##
## For each parameter, mcse is Monte Carlo standard error, n_eff is a crude measure of effective sample
```

###### Table 4, model_b1: Estimated effect of culling (95% CI)

```r
round((exp(coef(model_b1))[2]-1),3)*100                          # point estimate -13.7%
```

```
## TreatmentProactive
##               23.7
```

```r
round((exp(posterior_interval(model_b1, prob = 0.95))[2,1]-1),3)*100 # lower 95% CI
```

```
## [1] -23.7
```

```r
round((exp(posterior_interval(model_b1, prob = 0.95))[2,2]-1),3)*100 # upper 95% CI
```

```
## [1] 99.2
```

###### Table 4, model_b1: LOO

```r
loo(model_b1)
```

```
##
## Computed from 4000 by 20 log-likelihood matrix.
##
##          Estimate  SE
## elpd_loo    -77.6 2.5
## p_loo         2.2 0.5
## looic       155.3 5.0
## ------
## MCSE of elpd_loo is 0.0.
## MCSE and ESS estimates assume MCMC draws (r_eff in [0.4, 1.3]).
##
## All Pareto k estimates are good (k < 0.7).
## See help('pareto-k-diagnostic') for details.
```

#### Table 4, Model b2

```r
set.seed(8745)
model_b2 <- stan_glm.nb(Incidence ~ Treatment + log(Hist3yr) + log(Baseline),
                        data = rbctconf_outside, prior_intercept=normal(0, 1, autoscale=TRUE),
                        prior_aux= cauchy(0, 5),
                        prior = normal(0, 1),
                        diagnostic_file = file.path(tempdir(), "df.csv"),refresh=0)

summary(model_b2, digits=3, probs=c(0.025,0.5,0.975))
```

```
##
## Model Info:
##  function:     stan_glm.nb
##  family:       neg_binomial_2 [log]
##  formula:      Incidence ~ Treatment + log(Hist3yr) + log(Baseline)
##  algorithm:    sampling
##  sample:       4000 (posterior sample size)
##  priors:       see help('prior_summary')
##  observations: 20
##  predictors:   4
##
## Estimates:
##                         mean   sd     2.5%   50%    97.5%
## (Intercept)           -0.336  1.058 -2.435 -0.349  1.727
## TreatmentProactive     0.218  0.171 -0.126  0.221  0.547
## log(Hist3yr)           0.281  0.130  0.007  0.282  0.538
## log(Baseline)          0.634  0.230  0.171  0.633  1.094
## reciprocal_dispersion 12.703  7.026  4.434 11.076 30.060
##
## Fit Diagnostics:
##            mean   sd     2.5%   50%    97.5%
## mean_PPD 28.279  3.430 21.950 28.100 35.550
##
## The mean_ppd is the sample average posterior predictive distribution of the outcome variable (for de
##
## MCMC diagnostics
##                       mcse  Rhat  n_eff
## (Intercept)           0.016 0.999 4536
## TreatmentProactive    0.003 1.000 3291
## log(Hist3yr)          0.002 1.000 3665
## log(Baseline)         0.004 1.000 4139
## reciprocal_dispersion 0.149 1.002 2215
## mean_PPD              0.056 1.000 3725
## log-posterior         0.040 1.001 1769
##
## For each parameter, mcse is Monte Carlo standard error, n_eff is a crude measure of effective sample
```

```r
# Parameter values give increase in incidence of 23.6% (CrI -12%, 174%)
#This does not align with values given in Mills et al 2024b
```

###### Table 4, model_b2: Estimated effect of culling (95% CI)

```
round((exp(coef(model_b2))[2]-1),3)*100                              # point estimate
```

```
## TreatmentProactive
##               24.7
```

```
round((exp(posterior_interval(model_b2, prob = 0.95))[2,1]-1),3)*100 # lower 95% CI
```

```
## [1] -11.8
```

```
round((exp(posterior_interval(model_b2, prob = 0.95))[2,2]-1),3)*100 # upper 95% CI
```

```
## [1] 72.9
```

###### Table 4, model_b2: LOO

```
loo(model_b2)
```

```
## Warning: Found 1 observation(s) with a pareto_k > 0.7. We recommend calling 'loo' again with argumen
```

```
##
## Computed from 4000 by 20 log-likelihood matrix.
##
##          Estimate  SE
## elpd_loo    -76.0 4.4
## p_loo         4.3 1.5
## looic       152.0 8.8
## ------
## MCSE of elpd_loo is NA.
## MCSE and ESS estimates assume MCMC draws (r_eff in [0.5, 1.2]).
##
## Pareto k diagnostic values:
##                          Count Pct.    Min. ESS
## (-Inf, 0.7]   (good)     19    95.0%   655
##    (0.7, 1]   (bad)       1     5.0%   <NA>
##    (1, Inf)   (very bad)  0     0.0%   <NA>
## See help('pareto-k-diagnostic') for details.
```

##### Table 4, Model c.1: Using herd-years-at-risk as an offset and an explanatory variable and no varying intercepts for triplets

```
#Code given by Mills et al: (asrs1_outside_aB)
set.seed(7264)
model_c1 <- stan_glm.nb(Incidence ~ Treatment + log(hdyrsrisk) + log(Hist3yr),
                            offset = log(hdyrsrisk),
                            prior_intercept=normal(0,10), prior=normal(0,10),
                            data = rbctconf_outside, refresh=0)
#offset appears as both an explanatory variable and offset
#software ignores the offset
#It is a non offset model
```

###### Table 4, model_c1: Estimated effect of culling (95% CI)

```
round((exp(coef(model_c1))[2]-1),3)*100                              # point estimate
```

```
## TreatmentProactive
##               26.6
```

```
round((exp(posterior_interval(model_c1, prob = 0.95))[2,1]-1),3)*100 # lower 95% CI
```

```
## [1] -19.9
```

```
round((exp(posterior_interval(model_c1, prob = 0.95))[2,2]-1),3)*100 # upper 95% CI
```

```
## [1] 99.7
```

###### Table 4, model_c1: LOO

```
loo(model_c1, k_threshold = 0.7)
```

```
##
## Computed from 4000 by 20 log-likelihood matrix.
##
##          Estimate  SE
## elpd_loo    -74.9 2.0
## p_loo         1.9 0.3
## looic       149.8 4.0
## ------
## MCSE of elpd_loo is 0.0.
## MCSE and ESS estimates assume MCMC draws (r_eff in [0.5, 1.2]).
##
## All Pareto k estimates are good (k < 0.7).
## See help('pareto-k-diagnostic') for details.
```

###### modelc_1a: in comparison to model_c1, without the offset coded

```
set.seed(9836)
model_c1a<-stan_glm.nb(Incidence~Treatment+log(hdyrsrisk)+log(Hist3yr),
                       prior_intercept=normal(0,10), prior=normal(0,10),
                       data = rbctconf_outside, refresh=0)
summary(model_c1)
```

```
##
```

```
## Model Info:
##  function:     stan_glm.nb
##  family:       neg_binomial_2 [log]
##  formula:      Incidence ~ Treatment + log(hdyrsrisk) + log(Hist3yr)
##  algorithm:    sampling
##  sample:       4000 (posterior sample size)
##  priors:       see help('prior_summary')
##  observations: 20
##  predictors:   4
##
## Estimates:
##                       mean   sd   10%   50%   90%
## (Intercept)          -0.8    1.3 -2.5  -0.8   0.7
## TreatmentProactive    0.2    0.2 -0.1   0.2   0.5
## log(hdyrsrisk)       -0.4    0.2 -0.7  -0.4  -0.2
## log(Hist3yr)          0.2    0.2  0.0   0.2   0.4
## reciprocal_dispersion  5.7    2.0  3.4   5.4   8.3
##
## Fit Diagnostics:
##            mean   sd   10%   50%   90%
## mean_PPD 29.7    5.1 23.6  29.1  36.1
##
## The mean_ppd is the sample average posterior predictive distribution of the outcome variable (for de
##
## MCMC diagnostics
##                       mcse Rhat n_eff
## (Intercept)           0.0  1.0  4459
## TreatmentProactive    0.0  1.0  3655
## log(hdyrsrisk)        0.0  1.0  3976
## log(Hist3yr)          0.0  1.0  3860
## reciprocal_dispersion 0.0  1.0  3522
## mean_PPD              0.1  1.0  3510
## log-posterior         0.0  1.0  1553
##
## For each parameter, mcse is Monte Carlo standard error, n_eff is a crude measure of effective sample
```

```r
summary(model_c1a)
```

```
##
## Model Info:
##  function:     stan_glm.nb
##  family:       neg_binomial_2 [log]
##  formula:      Incidence ~ Treatment + log(hdyrsrisk) + log(Hist3yr)
##  algorithm:    sampling
##  sample:       4000 (posterior sample size)
##  priors:       see help('prior_summary')
##  observations: 20
##  predictors:   4
##
## Estimates:
##                       mean   sd   10%   50%   90%
## (Intercept)          -0.8    1.2 -2.4  -0.8   0.8
## TreatmentProactive    0.2    0.2 -0.1   0.2   0.5
## log(hdyrsrisk)        0.6    0.2  0.3   0.6   0.8
## log(Hist3yr)          0.2    0.2  0.0   0.2   0.4
```

```
## reciprocal_dispersion  5.7    2.0  3.3   5.5   8.3
##
## Fit Diagnostics:
##            mean   sd   10%   50%   90%
## mean_PPD 29.6    5.0 23.8  29.1  35.9
##
## The mean_ppd is the sample average posterior predictive distribution of the outcome variable (for de
##
## MCMC diagnostics
##                       mcse Rhat n_eff
## (Intercept)           0.0  1.0  4436
## TreatmentProactive    0.0  1.0  4049
## log(hdyrsrisk)        0.0  1.0  3863
## log(Hist3yr)          0.0  1.0  3343
## reciprocal_dispersion 0.0  1.0  3581
## mean_PPD              0.1  1.0  3636
## log-posterior         0.0  1.0  1647
##
## For each parameter, mcse is Monte Carlo standard error, n_eff is a crude measure of effective sample
```

```
#identical results but with log(hrdyrsrisk) parameter translated by 1.
```

##### Correct code for model_c1 should be

```
set.seed(6472)
model_c1_offset<-stan_glm.nb(Incidence~Treatment+log(Hist3yr),
                      offset = log(hdyrsrisk),
                      prior_intercept=normal(0,10), prior=normal(0,10),
                      data = rbctconf_outside, refresh=0)
summary(model_c1_offset, digits=3, probs=c(0.025,0.5,0.975))
```

```
##
## Model Info:
##  function:     stan_glm.nb
##  family:       neg_binomial_2 [log]
##  formula:      Incidence ~ Treatment + log(Hist3yr)
##  algorithm:    sampling
##  sample:       4000 (posterior sample size)
##  priors:       see help('prior_summary')
##  observations: 20
##  predictors:   3
##
## Estimates:
##                         mean   sd     2.5%   50%    97.5%
## (Intercept)           -3.183  0.498 -4.152 -3.189 -2.193
## TreatmentProactive     0.218  0.250 -0.270  0.217  0.720
## log(Hist3yr)           0.172  0.177 -0.186  0.170  0.518
## reciprocal_dispersion  4.886  1.693  2.206  4.648  8.857
##
## Fit Diagnostics:
##            mean   sd     2.5%   50%    97.5%
## mean_PPD 32.496  5.882 22.650 31.950 45.351
##
## The mean_ppd is the sample average posterior predictive distribution of the outcome variable (for de
```

```
##
## MCMC diagnostics
##                         mcse  Rhat  n_eff
## (Intercept)             0.009 1.000 3398
## TreatmentProactive      0.004 0.999 3785
## log(Hist3yr)            0.003 1.000 3459
## reciprocal_dispersion 0.031 1.000 2958
## mean_PPD                0.094 1.000 3946
## log-posterior           0.040 1.001 1454
##
## For each parameter, mcse is Monte Carlo standard error, n_eff is a crude measure of effective sample
```

##### Table 4, model_c1_offset: Estimated effect of culling (95% CI)

```r
round((exp(coef(model_c1_offset))[2]-1),3)*100                    # point estimate
```

```
## TreatmentProactive
##               24.2
```

```r
round((exp(posterior_interval(model_c1_offset, prob = 0.95))[2,1]-1),3)*100 # lower 95% CI
```

```
## [1] -23.7
```

```r
round((exp(posterior_interval(model_c1_offset, prob = 0.95))[2,2]-1),3)*100 # upper 95% CI
```

```
## [1] 105.4
```

##### Table 4, model_c1: LOO

```r
loo(model_c1_offset)
```

```
##
## Computed from 4000 by 20 log-likelihood matrix.
##
##          Estimate  SE
## elpd_loo    -77.4 2.3
## p_loo         1.9 0.4
## looic       154.8 4.6
## ------
## MCSE of elpd_loo is 0.0.
## MCSE and ESS estimates assume MCMC draws (r_eff in [0.5, 1.1]).
##
## All Pareto k estimates are good (k < 0.7).
## See help('pareto-k-diagnostic') for details.
```

#### Table 4, Model c2 without offset

Likewise the code for model c2 has no offset for the same reasons. Code for model_c2 given by mills et al: (model rs_1_outside_aB_improved)

```
model_c2 <- stan_glm.nb(Incidence ~ Treatment + log(hdyrsrisk) + log(Hist3yr),
                                     prior_intercept = normal(0, 5),
                                     prior = normal(0, 1),
                                     prior_aux= cauchy(0, 5),
                                     offset = log(hdyrsrisk),
                                     data = rbctconf_outside, refresh=0)

#This is not an offset model.
```

#### Table 4, Model d1, no culling effect from table 4 (=model rs2B)

```r
set.seed(85673)
model_d1 <- stan_glm.nb(Incidence ~ log(hdyrsrisk) + log(Hist3yr),
                        prior_intercept=normal(0,10), prior=normal(0,10),
                        data=rbctconf_outside,
                    diagnostic_file = file.path(tempdir(),"df.csv"),refresh=0)

summary(model_d1, digits=3, probs=c(0.025,0.5,0.975))
```

```
##
## Model Info:
##  function:     stan_glm.nb
##  family:       neg_binomial_2 [log]
##  formula:      Incidence ~ log(hdyrsrisk) + log(Hist3yr)
##  algorithm:    sampling
##  sample:       4000 (posterior sample size)
##  priors:       see help('prior_summary')
##  observations: 20
##  predictors:   3
##
## Estimates:
##                          mean   sd     2.5%   50%    97.5%
## (Intercept)            -0.634  1.226 -3.004 -0.640  1.802
## log(hdyrsrisk)          0.587  0.206  0.181  0.588  0.986
## log(Hist3yr)            0.178  0.159 -0.151  0.180  0.488
## reciprocal_dispersion  5.708  1.944  2.697  5.463 10.143
##
## Fit Diagnostics:
##            mean   sd     2.5%   50%    97.5%
## mean_PPD 29.232  4.778 21.000 28.900 39.900
##
## The mean_ppd is the sample average posterior predictive distribution of the outcome variable (for de
##
## MCMC diagnostics
##                        mcse  Rhat  n_eff
## (Intercept)            0.019 1.000 4096
## log(hdyrsrisk)         0.003 0.999 3725
## log(Hist3yr)           0.003 0.999 3781
## reciprocal_dispersion 0.033 1.001 3543
## mean_PPD               0.079 1.000 3644
## log-posterior          0.036 1.001 1734
##
## For each parameter, mcse is Monte Carlo standard error, n_eff is a crude measure of effective sample
```

#### Table 4, Model_d1 improved

```r
model_d1i <- stan_glm.nb(Incidence ~ log(hdyrsrisk) + log(Hist3yr),
                                  prior=normal(0, 1),
                                  prior_aux = cauchy(0, 5),
                                  data=rbctconf_outside,
                       diagnostic_file = file.path(tempdir(), "df.csv"),
                       refresh=0)
```

```r
summary(model_d1i, digits=3, probs=c(0.025,0.5,0.975))
```

```
##
## Model Info:
##  function:     stan_glm.nb
##  family:       neg_binomial_2 [log]
##  formula:      Incidence ~ log(hdyrsrisk) + log(Hist3yr)
##  algorithm:    sampling
##  sample:       4000 (posterior sample size)
##  priors:       see help('prior_summary')
##  observations: 20
##  predictors:   3
##
## Estimates:
##                          mean   sd     2.5%   50%    97.5%
## (Intercept)           -0.684  0.796 -2.236 -0.702  0.903
## log(hdyrsrisk)         0.588  0.131  0.332  0.589  0.842
## log(Hist3yr)           0.193  0.109 -0.033  0.193  0.399
## reciprocal_dispersion 29.279 74.926  6.650 19.267 88.602
##
## Fit Diagnostics:
##            mean   sd     2.5%   50%    97.5%
## mean_PPD 28.700  2.952 23.400 28.525 34.800
##
## The mean_ppd is the sample average posterior predictive distribution of the outcome variable (for de
##
## MCMC diagnostics
##                       mcse  Rhat  n_eff
## (Intercept)           0.014 1.000 3316
## log(hdyrsrisk)        0.002 1.000 3089
## log(Hist3yr)          0.002 0.999 3316
## reciprocal_dispersion 3.656 1.007  420
## mean_PPD              0.049 1.000 3683
## log-posterior         0.039 1.000 1586
##
## For each parameter, mcse is Monte Carlo standard error, n_eff is a crude measure of effective sample
```

##### Table 4, model e, as coded in Mills et al. 2024b

```
set.seed(8489)
model_e <- stan_glm(Incidence~Treatment+A+B+C+D+E+F+G+H+I+log(Hist3yr)+log(Baseline),
                        data = rbctconf_outside,
                        offset=log(Baseline),
                        prior_intercept=normal(0, 2),
                        poisson,
                        prior = normal(0, 1),
                        diagnostic_file = file.path(tempdir(), "df.csv"),refresh=0)

#Again this codes the offset as both an explanatory variable and offset variable.
#This is effectively the original glm in Donnelly et al, but respecified in the
#Bayesian paradign
#All figures in the main text that relies on this model_e,
#assuming the offset are invalid.
#Also the effect size does not align with the reported effect size in Table 2b
#in Mills et al where it is reported.

#The same issues are also present in the coding and analysis for follow up and after.
```

###### Table 4, model e: Estimated effect of culling (95% CI)

```
round((exp(coef(model_e))[2]-1),3)*100                      # point estimate
```

```
## TreatmentProactive
##               26.8
```

```
round((exp(posterior_interval(model_e, prob = 0.95))[2,1]-1),3)*100 # lower 95% CI
```

```
## [1] 4.8
```

```
round((exp(posterior_interval(model_e, prob = 0.95))[2,2]-1),3)*100 # upper 95% CI
```

```
## [1] 53.4
```

###### Table 4, model e: LOO

```
loo(model_e, k_threshold = 0.7)
```

```
##
## Computed from 4000 by 20 log-likelihood matrix.
##
##          Estimate  SE
## elpd_loo    -70.4 2.3
## p_loo        12.2 1.5
## looic       140.7 4.6
## ------
## MCSE of elpd_loo is 0.3.
## MCSE and ESS estimates assume MCMC draws (r_eff in [0.4, 1.3]).
##
## All Pareto k estimates are good (k < 0.7).
## See help('pareto-k-diagnostic') for details.
```

##### Table 4, model e correctly specified

```
set.seed(18489)
model_e.c <- stan_glm(Incidence ~ Treatment + A + B + C + D + E + F + G + H
                      + I +log(Hist3yr),
                        data = rbctconf_outside,
                        offset=log(Baseline),
                        prior_intercept=normal(0, 2),
                        poisson,
                        prior = normal(0, 1),
                        diagnostic_file = file.path(tempdir(), "df.csv"),refresh=0)

summary(model_e.c, digits=5, probs=c(0.025,0.5,0.975))
```

```
##
## Model Info:
##  function:     stan_glm
##  family:       poisson [log]
##  formula:      Incidence ~ Treatment + A + B + C + D + E + F + G + H + I + log(Hist3yr)
##  algorithm:    sampling
##  sample:       4000 (posterior sample size)
##  priors:       see help('prior_summary')
##  observations: 20
##  predictors:   12
##
## Estimates:
##                       mean     sd       2.5%     50%      97.5%
## (Intercept)         -1.79246  0.26672 -2.32315 -1.79241 -1.27741
## TreatmentProactive   0.09281  0.09160 -0.07919  0.09052  0.27927
## A1                   0.28830  0.20395 -0.11433  0.29181  0.69266
## B1                   0.66884  0.16407  0.35614  0.66857  0.99668
## C1                   0.23877  0.16926 -0.08737  0.24077  0.56457
## D1                   0.26121  0.21720 -0.16732  0.25928  0.68518
## E1                   0.34205  0.18413 -0.00994  0.33644  0.70470
## F1                   0.05773  0.20037 -0.34239  0.05899  0.45208
## G1                  -0.07692  0.18445 -0.43931 -0.07508  0.28404
## H1                   0.57805  0.17351  0.23816  0.57395  0.92259
## I1                  -0.10895  0.22649 -0.56139 -0.10032  0.33119
## log(Hist3yr)         0.12220  0.08717 -0.04725  0.12105  0.29609
##
## Fit Diagnostics:
##            mean     sd       2.5%     50%      97.5%
## mean_PPD 28.53879  1.66903 25.40000 28.50000 31.85000
##
## The mean_ppd is the sample average posterior predictive distribution of the outcome variable (for de
##
## MCMC diagnostics
##                     mcse    Rhat    n_eff
## (Intercept)         0.00618 1.00059 1865
## TreatmentProactive 0.00183 1.00052 2501
## A1                  0.00478 1.00012 1817
## B1                  0.00415 1.00026 1566
## C1                  0.00430 1.00102 1552
## D1                  0.00473 1.00002 2107
```

```
## E1                 0.00442 1.00062 1736
## F1                 0.00495 1.00093 1637
## G1                 0.00445 1.00054 1722
## H1                 0.00431 0.99998 1617
## I1                 0.00492 1.00057 2119
## log(Hist3yr)       0.00170 1.00079 2624
## mean_PPD           0.02698 1.00014 3827
## log-posterior      0.05765 1.00043 1837
##
## For each parameter, mcse is Monte Carlo standard error, n_eff is a crude measure of effective sample
```

##### Table 4, model e.c: Estimated effect of culling (95% CI)

```
round((exp(coef(model_e.c))[2]-1),3)*100                       # point estimate
```

```
## TreatmentProactive
##                9.5
```

```
round((exp(posterior_interval(model_e.c, prob = 0.95))[2,1]-1),3)*100 # lower 95% CI
```

```
## [1] -7.6
```

```
round((exp(posterior_interval(model_e.c, prob = 0.95))[2,2]-1),3)*100 # upper 95% CI
```

```
## [1] 32.2
```

##### Table 4, model e.c: LOO

```
loo(model_e.c, k_threshold = 0.7)
```

```
##
## Computed from 4000 by 20 log-likelihood matrix.
##
##          Estimate   SE
## elpd_loo    -85.6  8.4
## p_loo        22.9  5.8
## looic       171.3 16.9
## ------
## MCSE of elpd_loo is 0.4.
## MCSE and ESS estimates assume MCMC draws (r_eff in [0.4, 1.5]).
##
## All Pareto k estimates are good (k < 0.7).
## See help('pareto-k-diagnostic') for details.
```

##### Model 11. as coded in Mills et al. 2024b

```
model11 <-glmmTMB(Incidence~offset(log(hdyrsrisk)),genpois,  data =rbctconf_outside)
```

```
## Warning in (function (start, objective, gradient = NULL, hessian = NULL, :
## NA/NaN function evaluation
```

```
set.seed(94060000)
pp_check(model11) #Figure 3 supplementary material 3.
```

```
## The model has an integer or a discrete response variable.
##   It is recommended to switch to a dot-plot style, e.g.
```

```
##   `plot(check_model(model), type = "discrete_dots"`.
```

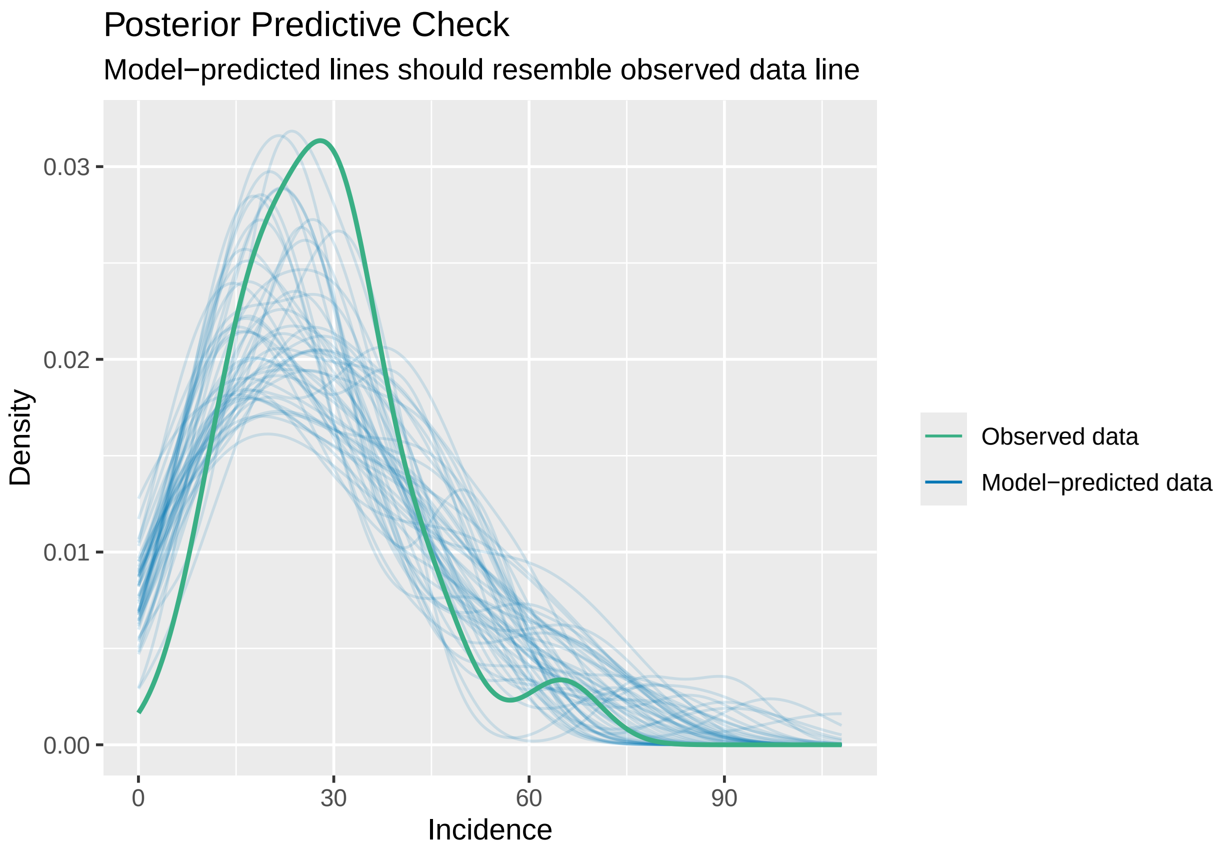
