## Supplemental Material 1 for "Randomised Badger Culling Trial lacks evidence for proactive badger culling effect on tuberculosis in cattle: comment on Mills et al. 2024, Parts I & II"

**Supplementary material file.**

Study justification and context.

**Timeline to reference documentation**

Mills et al. 2024a,b(1,2) published 21st August 2024 are based largely(3) on the online preprint of the primary re-evaluation of the Randomised Badger Culling Trial (RBCT), 1998-2005, first posted in 2022(4) by Torgerson et al., and published in *Scientific Reports* on 15^th^ July 2024(5). A second preprint in May 2023(6), included text adjustments and additional study context(7). In relation to the Mills et al. publications, a request to the publishing journal (Royal Society Open Science) and to the defending authors was made in August 2024 to delay publication due to issues within proof versions of the Mills et al. papers(8,9). This was not allowed. However changes to the following text from the proof were made to the final published version of Mills et al., 2024a:, “*Another approach to modelling overdispersed data is the usage of an offset variable which enables modelling the count variable (here confirmed herd breakdowns) as a rate, and the usage of an offset variable means that the corresponding regression coefficient is constrained to be 1.”* While this had escaped the attention of the peer-reviewers, it was updated after Torgerson drew attention to it, to: *“Another possible option for a Poisson regression model is the usage of an offset variable which enables modelling the count variable (here confirmed herd breakdowns) as a rate, and the usage of an offset variable means that the corresponding regression coefficient is constrained to be 1.”* It is of note that the Department of Environment Food and Rural Affairs (Defra) may use preprint/proof/unpublished documents as reference for advice and justification of the decisions regarding badger interventions. This despite initial cautionary remarks (10) that have been expanded upon since Mills et al.(2024 a,b) in the present study.

**Evidence of pre-planning of the Randomised Badger Culling Trial (RBCT)**

Effort to clarify a full citation of a reported note by Donnelly and Cox in 1998, from the year that the experiment began, and referred to in the Mills et al. proof version (8) was, upon checking with the first author, *“meant to refer to the published 2000 document, alongside the published report by Professor Mollison”* (Mills *pers com.* 22.08.24), for the final publications (1,2). Clarifying the timing of the available evidence surrounding the origin of RBCT analytical planning, involved a search for all other *Independent Specialist Group on Cattle TB (ISG)* documents relating to RBCT statistical planning and analysis, including checks by the defending authors. The first passing reference regarding statistics and the RBCT is within Appendix 3 “Statistical Analyses” of the *first report of the ISG*, published in July 1998(11). However, it simply states: *“ Incidence rates will be computed and analysed both on a per head of cattle and per farm basis.”* The second report of the ISG was published in December 1999 (12) Paragraphs 4.1.5 to 4.1.15 relating to “Statistical power” and “Assumptions”, within “4.1 Trial design”. The November 2000 Statistical Auditor Report(13) has a link to 'Notes on statistical aspects of the badger culling trial'(14) which is Appendix III of a report to a parliamentary agricultural select committee(15) that also states: “At the time of our evidence session with the ISG, we had available to us the Group's *Notes on statistical aspects of the badger culling trial* prepared for and endorsed by Professor Denis Mollison.” There was a third and fourth audit by Mollison in 2004 and 2005, respectively and this and additional information is reviewed in the file Supplementary Information 1 of Torgerson et al. 2024 (5). This further demonstrates that the original intention for statistical analysis was to analyse using rate rather than count.

Beyond this, there is no indication of any preplanned analytical protocol in the modern experimental sense of the term. The detailed practical logistics of the mass intensive cage-trapping and shooting of badgers and monitoring of cattle herds for disease, followed set guidelines. But not the detailed statistical analysis that, for any substantive intervention/control trial, is made public in advance. This insufficiency undermines any claim that the RBCT was, as conducted (as opposed to designed), a preplanned experiment. This allows speculation about which models were tested, at what point the choice of model was made, and upon whose advice. The 2006 proactive culling finding matched ideas in 1985 of Anderson and Trewhella(16), but was chosen against the outcomes of other more standard analytical options. It found the minimum detectable effect of 20%, below which the 10-pair study lacked statistical power to detect(17).

**Importance of epidemiological discovery in relation to tuberculin testing since 2006**

One of the key drivers of concerns regarding the RBCT analysis is large change to the understanding of bTB diagnosis since 1997 when the proposal for the study was published(18). Further, how deductions from extensive historic 19th and 20^th^ century studies(19) have meaning through subsequent replication, supporting veterinary evidence and interpretation. This includes insights gathered from slaughtered cattle during current epidemics in GB and Ireland since 2000 (e.g. refs 20-23), and particularly the discovery that OTF-W only (‘confirmed’) breakdowns do not better reflect infections originating from cattle contact with wildlife(24) than total breakdowns, and indeed are less likely to reflect rate of new infection or breakdown. Central to modern considerations is the more refined understanding of the meaning of the SICCT sensitivity and specificity (see main text) in 2018(25). This also helped show how the ‘hidden bTB reservoir’ was embedded in herds released from restrictions and traded, and removing the previous perception of the need for an external vector to explain new infection or recrudescence. This explains, for example, bTB proliferating in the first (pilot) badger cull area in Gloucestershire, despite heavy badger culling for nine years, since 2013. Regular Government epidemiological monitoring reports(26) found bTB infection retained in chronically infected herds in Cumbria, thought to be recently infected by imported stock from Northern Ireland(27), where badgers were subjected to unlimited intensive culling for a period of three years. More recently, an independent monitoring report of the same data was followed by the withdrawal of the Animal and Plant Agency veterinary *Disease Report Form* and *‘Risk Pathways’* guidance for inspecting veterinarians in 2023(28) due to large bias in attribution of infection to the presence of badger in the landscape when it was default assumption, founded on RBCT science.

**Fragility of study effect caused by restricted epidemiological understanding in 1997.**

There is a substantial confounding issue identified in hindsight of the greater understanding of the SICCT test limitations. In 1997, the general cattle veterinarian view was that infected cows could not be infectious unless they had visible lesions at the abattoir and that badgers were responsible for most new infections. Both views are accepted as wrong, which is not disputed, but this was the background of the RBCT specification and study. This matter impacts hugely on study design. Prior to each cull and control area commencing during the RBCT, efforts were made to bring them into the same annual testing frequency (from every-2 or 4 years) to create parity. However, there was trading of stock ‘cleared’ by the SICCT that were thought to be free from infection when 20% or more were not. The proportion of infected cattle moving into a trial area will have varied because the surrounding parishes were on variable test duration. Thus during the trial, unknown volumes of disease were moving into cull and control areas, according to distance travelled and testing frequency surrounding each study area. This was highly variable within and between the 10 pairs. While the pre-cull adjustments for analysis could to some extent adjust for the pre-trial period, the data during the trial were exposed to this unknown factor for the years of study, creating potentially a hidden random imbalance between the cull and control areas, the impact of which cannot be subsequently disentangled from the effect of the intervention. This is of particular concern here given the small number of areas in the trial and so the fragility of the observed results to any further replication, where a different random imbalance would be observed, were the same design used.

There is a common misconception that proactive culling intervention during the RBCT caused a reduction in all 10 cull areas compared with control. The real-time data show incidence approximately doubled in all areas(17). However, the effect of widely variable use of the SICCT being introduced within and beyond the area boundaries before the experiment cannot be factored in. The post-experiment discovery of unknown unevenness of such an important determining factor is unfortunate as it deadens the veracity of all analyses and any value of the experiment. Studies of herd incidence after the RBCT ended, from 2013 onwards show a consistent response in reducing incidence following the introduction of annual SICCT use over a 2-5 years period and before the roll out of badger culling(29) and the perception of a substantial year-3 effect.

**Contemporary comparative bTB control response at the national scale**

Mills et al. 2024a(1) refers to bTB control generally in England and Wales, in recent decades, countries where the general pattern of disease incidence detection has run in long and consistent parallel according to online Defra monitoring(30) despite Government in Wales prohibiting mass culling of badgers(31). From (31):,

*“In Wales, widespread badger culling no longer takes place and the Bovine TB Eradication Programme has outlined the government’s ambition to prohibit badger culling and instead promote a badger vaccination policy. There has been a general long-term trend in Wales of declining herd incidents over the past decade and 94.6% of herds were estimated to be TB-free as of Q3 2023, yet recent trends include 1.7% and 6.4% increases to the reported number of animals slaughtered and new herd incidents between October 2021 to September 2022 and October 2022 to September 2023.”*

Contrary to the suggestion of Mills et al. 2024a(1), badger culling has always been prohibited in Wales, other than on a very small experimental scale. An extensive programme of badger vaccination in an *Intensive Action Area* (IAA) in North Pembrokeshire was ended once 5,300 badgers had been vaccinated between 2012 and 2016, across approximately 260 sq.km of land(32), with no demonstrable benefits and no further badger vaccination at scale in the subsequent 8 years. Wales saw 8 farms subject to a mini-study of Test-Vaccinate-Remove (TVR) from 2017 to 2023(33) at a cost of around £1.6 Million, where around 16 badgers per year were euthanised. During 2020 it was revealed that many vaccinated badgers had tested false-positive, and were hence destroyed in error, due to a conflict between the cross reactivity of BCG Sofia vaccine and the Dual Path Platform antibody test. So the TVR work was abandoned. Any characterisation of long term comparative data between Wales and England observes extremely similar trends in herd incidence, providing convincing evidence that similar levels of disease control are achieved with or without badger culling. This strongly corroborates findings that proactive badger culling, under fewer operating constraints than the RBCT (2013-2020) in England has resulted in unmeasurable effects(29).

The reference to a recent increase of bTB incidents in new parts of north Wales since 2021 is well recorded and breaks the long term trend in a useful demonstration of how sudden trading of infected cattle into a large naïve area is a key threat to elimination. For example, Whole Genome Sequencing (WGS) checks of spoligotypes has demonstrated cattle movements to the Wales *Low TB Area* from a higher risk TB area during 2017. Similarly, the increase in incidents in the Wales *Intermediate TB Area North* is driven by movements across county and country borders of west Shropshire, southwest Cheshire and Wales. *High TB Area West* has a considerable number of large dairy herds. There is a high number of recurrent breakdowns which suggests either a high reinfection rate or infection that persists in the herd after it is declared Officially TB-Free(34). Breakdown figures up to March 2024 across most of Wales suggest good progress, despite an increase of 20% in the High West, but a decrease of **-53%** in Wales-Low, **-33%** in Intermediate North and **-29%** in Intermediate mid. While progress is held back by the new infections from England, it is wrong to infer Wales is a country where bTB control is less effective than in England(35). The Deputy first Minister of Wales, in a statement in May 2024(36),stated the following when comparing bTB control efforts across the two countries:

*“But just to be clear, from 2012, which is the year before badger control policy in England, to 2023, on the latest published data, the herd incidence in England decreased from 9.8 to 7.3; it was a 26 per cent decrease. In Wales, over the same period, herd incidence decreased from 10 to 6.8. It's a 31.3 per cent decrease. I simply put that on record—those are Department for Environment, Food and Rural Affairs figures, by the way—to say that we are doing things differently in Wales, in line with our programme for government, but we're also succeeding in many ways.”*

**The applied importance of intervention/control studies.**

The Dunnett Review in 1986(37) found with respect to the badger control policy, that *“no definitive statement concerning the success of the [trapping/gassing] strategy can be made, since the data seem equally consistent with two entirely different hypotheses..”* This came after local veterinary inspectors in South West England had proposed anecdotally that badgers were a cause of around half (50%) of bTB cattle herd breakdowns. Causality of significant badger infection of cattle had previously been prematurely accepted by Zuckerman’s Review in 1980(38). However, his carefully worded reflection in 1988(39) suggesting the need for *“further evidence to support the proposition that a direct association does exist between the disease in the two animals”* seems to reflect Dunnett’s subsequent caution.

The confusion of association and causation was of great scientific concern in the 1960’s (40)

and it was the fitting of a model to the 50% causation theory in 1985 by Anderson, who had also advised Zuckerman in 1980(43), that firmly anchored the belief-system within the farming and veterinary communities that badgers played a highly significant role in sustaining infection in cattle herds. Despite small-scale culling in England and Ireland, a 10-year scientific hiatus followed, and with bTB testing still reducing in the mid to late 1980’s(41) bTB breakdowns began to steadily rise, not helped by efforts to maximise the size of the national herd in the 1970’s under European Community rules that led to food mountains/lakes of over-production(42). A further review was led by Krebs at Oxford University in 1997(18), with Anderson appointed to his *Independent Scientific Review Group* and MacDonald (who was not a member) advising on aspects of detecting whether badgers are the key vector, or not. This group proposed a randomised field trial experiment that became the RBCT. It was by necessity a relatively small experiment with just 20 units with 2 treatments, giving ten units (proactive and reactive culling) compared with ten control areas. Yet logistically, in terms of field effort, it was extremely onerous. Conducted by Government staff, together with farmers, it was expensive, with the ambition to kill and remove all badgers within the proactive cull areas, an aim that was abandoned once the practicalities of doing so became apparent.

The most recent Governmental review was in 2018 by Godfray, who had played a significant statistical advisory role in the RBCT. The 2018 Review group included Donnelly and the report(43) addressed the original 2014 Governmental Bovine TB Strategy(44). It had recommended a post-cull comparison between a) a small number of culled areas with badgers left to recolonise, with b) those left to recolonise and subject to badger vaccination, that the Government of the day chose not to pursue. While the extensive restocking of bTB-untested stock after the Foot and Mouth Disease (FMD) epidemic of 2001 (following the relaxing of bTB testing for a period) is associated with a dramatic jump in bTB herd breakdowns(45), any claim that local disease radiation was caused by badgers, as opposed to traded cattle was anecdotal and not evidence-based. In retrospective consideration in 2013, Donnelly found that, if considered, the FMD impact effectively invalidated the analytical strength of the RBCT experiment(46), a finding largely ignored. While claims that the present English culls 2013-present were established in a manner where direct efficacy cannot be demonstrated(1), the database on English herds over that period, testing locations and results, and badger cull geolocation and data is not publicly accessible, so this is not verifiable and seems unlikely given appropriate resource. Following judicial scrutiny and submissions in relation to Supplementary Badger Culling, there is an expectation of data being sufficient for ‘learning and adaptation’ from the interpretation of results(47) to satisfy legislative need(48). Anecdotal criticism(49) of the main independent study of the to-2020 data, including pre-cull data(29) has still not been published, despite its central importance.

The inability to show any clear benefit from badger culling over the 2013-2020 period (29,50,51) and the manner in which disease has been retained in some of the High Risk Area’s earliest badger culling areas, has not brought academic confidence that it is ‘worthwhile’(52). While a group led by the Government's Chief Scientific Advisor in 2007 overturned the conclusions of the final RBCT report (17,53), there is no appearance of that reappraisal including scrutiny of the RBCT modelling. Therefore, it missed the problems raised in the first major RBCT re-evaluation(4) and in this present study, as a response to Mills et al. 2024a and b(1,2). It was the ‘King Group’ recommendations that were taken forward after 2010, without awareness of the statistical uncertainties that have grown since clarification of the SICCT sensitivity and specificity (see above), including at severe interpretation, and as an earlier report first flagged(54) how sensitive the 2006 model(55) is to potential variations in these diagnostic characteristics. Good research design and data analysis are recognised as a vital part of the science process(56).

3

Reviewer Report Mills et al. An extensive re-evaluation of evidence and analyses of the Randomised Badger Culling Trial (RBCT) I: Within proactive culling areas. 2024/05/03

https://www.webofscience.com/wos/review-content

4.

Torgerson PR, Hartnack S, Rasmussen P, Lewis F, Langton TES. Absence of effects of widespread badger culling on tuberculosis in cattle PREPRINT 2022

5.

Torgerson PR, Hartnack S, Rasmussen P, Lewis F, Langton TES. Absence of effects of widespread badger culling on tuberculosis in cattle. Sci Rep. 2024 Jul 15;14(1):16326.

6.

Torgerson PR, Hartnack S, Rasmussen P, Lewis F, Langton TES. Absence of effects of widespread badger culling on tuberculosis in cattle PREPRINT 2023

7.

Torgerson PR, Hartnack S, Rasmussen P, Lewis F, Langton TES.

Absence of effects of widespread badger culling on tuberculosis in cattle. 2023. Supplementary Information 1: Justification for study. <https://static-content.springer.com/esm/art%3A10.1038%2Fs41598-024-67160-0/MediaObjects/41598_2024_67160_MOESM1_ESM.pdf>

8.

Mills CL, Woodroffe R, Donnelly CA. a. An extensive re-evaluation of evidence and analyses of the Randomised Badger Culling Trial (RBCT) I: Within proactive culling areas. R Soc Open Sci. 2024 proof copy August 2024

9.

Mills CL, Woodroffe R, Donnelly CA. b. An extensive re-evaluation of evidence and analyses of the Randomised Badger Culling Trial II: In neighbouring areas. R Soc Open Sci. 2024 proof copy August 2024

10.

Torgerson, P.R. and T.E.S. Langton 2024 Interim report on the August 2024 pre-publication response to the July 2024 re-evaluation of evidence and analyses of proactive culling (published in 2006), as a part of the Randomised Badger Culling Trial (RBCT), 1998-2005. 18 August 2024, 14 pp

11.

Independent Scientific Group on Cattle TB. 1998. Bovine TB: Towards a sustainable policy to control TB in cattle. First Report to the Rt Hon Dr Jack Cunningham MP from Appendix 3. https://webarchive.nationalarchives.gov.uk/ukgwa/20081023164230/http:/www.defra.gov.uk/animalh/tb/isg/isgrep1.htm

12.

Independent Scientific Group on Cattle TB. 1999. An Epidemiological Investigation into Bovine Tuberculosis: Towards a Sustainable Policy to Control TB in Cattle, Second Report of the Independent Scientific Group on Cattle TB, Presented to the Minister of Agriculture, Fisheries and Food, The Rt Hon Nick Brown MP, December 1999. Published 1999.

https://webarchive.nationalarchives.gov.uk/ukgwa/20081023164252/http://www.defra.gov.uk/animalh/tb/isg/report/contents.htm.

13.

Mollison D. First report of the Statistical Auditor on the badger culling trial. London: Ministry of Agriculture, Fisheries and Food; 2000.

14.

Independent Scientific Group on Cattle TB. 2000. Notes on statistical aspects of the badger culling trial. Appendix 3 of a report the a parliamentary select committee ( 13) prepared for and endorsed by Professor Mollison.

<https://webarchive.nationalarchives.gov.uk/ukgwa/20081024234232mp_/http://www.defra.gov.uk/animalh/tb/isg/pdf/rbct_annex1.pdf>

15.

Select Committee on Agriculture. 2001. Badgers and Bovine Tuberculosis First Report: follow up: <https://publications.parliament.uk/pa/cm200001/cmselect/cmagric/92/9203.htm>:

16.

Anderson RM, and Trewhella W. Population dynamics of the badger (Meles meles) and the epidemiology of bovine tuberculosis (Mycobacterium bovis). Philos Trans R Soc Lond B Biol Sci. 1985 Sep 12;310(1145):327-81. doi: 10.1098/rstb.1985.0123. PMID: 2865760.

17.

Bourne FJ, Donnelly CA, Cox DR, Gettinby G, McInerney JP, Morrison WI, et al. The scientific evidence—final report of the independent scientific group on cattle TB. London: Independent Scientific Group on Cattle TB. London, Independent Scientific Group on Cattle TB; 2007 p. 289.

18.

Krebs, J.R., Anderson, R.M., Clutton-Brock, T., Morrison, W.I., Young, D., & Donnelly, C.A. Bovine Tuberculosis in Cattle and Badgers. Ministry of Agriculture, Fisheries and Food (MAFF) Publications, London (1997).

19.

Francis, J. 1947 Bovine Tuberculosis: Including Contrast with Human Tuberculosis. Pp. 220 Staples Press Ltd. London

20.

Allen AR, Skuce RA, Byrne AW. Bovine Tuberculosis in Britain and Ireland - A Perfect Storm? the Confluence of Potential Ecological and Epidemiological Impediments to Controlling a Chronic Infectious Disease. Front Vet Sci. 2018 Jun 5;5:109. doi: 10.3389/fvets.2018.00109. Erratum in: Front Vet Sci. 2019 Jul 02;6:213. doi: 10.3389/fvets.2019.00213. PMID: 29951489; PMCID: PMC6008655.

21.

Langton, Tom and Paul Torgerson. 2024c TB testing and transmission Veterinary Record. Letters and notices. 6 July 2024

22.

Langton, Tom. 2024a New TB breakdowns fall in England. Veterinary Record. Letters and notices. 17/24 February 2024

23.

Langton, Tom. 2024b Bovine TB in cattle: true burden of disease untouched? Veterinary Record. Letters and notices. 30 March 2024

24.

Birch, C.P.D., Bakrania, M., Prosser, A. et al. Difference in differences analysis evaluates the effects of the badger control policy on bovine tuberculosis in England. Sci Rep 14, 4849 (2024). https://doi.org/10.1038/s41598-024-54062-4

25.

Nuñez-Garcia J, Downs SH, Parry JE, Abernethy DA, Broughan JM, Cameron AR, et al. Meta-analyses of the sensitivity and specificity of ante-mortem and post-mortem diagnostic tests for bovine tuberculosis in the UK and Ireland. Prev Vet Med. 2018 May 1;153:94–107.

26.

Gov.uk 2024. Research and analysis. April 2024: TB hotspots in the Low Risk Area of England

Updated 11 April 2024

https://www.gov.uk/government/publications/bovine-tb-hotspots-in-the-low-risk-area-of-england/april-2024-tb-hotspots-in-the-low-risk-area-of-england

27.

Rossi, G., Crispell, J., Brough, T., Lycett, S. J., White, P. C. L., Allen, A., Ellis, R. J., Gordon, S. V., Harwood, R., Palkopoulou, E., Presho, E. L., Skuce, R., Smith, G. C., & Kao, R. R. (2022). Phylodynamic analysis of an emergent Mycobacterium bovis outbreak in an area with no previously known wildlife infections. Journal of Applied Ecology, 59, 210–222. https://doi.org/10.1111/1365-2664.14046

28.

Griffiths, L.M., Griffiths, M.J., Jones, B.M., Jones, M.W., Langton, T. E. S., Rendle, R.M., & P.R. Torgerson. 2023. A bovine tuberculosis policy conundrum in 2023. On the scientific evidence relating to the Animal and Plant Health Agency/DEFRA policy concept for ‘Epidemiological’ badger culling. An independent report by researchers and veterinarians to Defra and the UK Parliament. Plus 2024 addendum. Badger Crowd Web pages accessed August 29 2024.

29.

Langton TES, Jones MW, McGill I. Analysis of the impact of badger culling on bovine tuberculosis in cattle in the high-risk area of England, 2009–2020. Vet Rec 2022; doi:10.1002/vetr.1384

30.

GOV.UK Bovine TB Interactive . http://3.9.48.73/bTB/

31.

Welsh Government Consultation Document 2021. A Refreshed TB eradication programme. Issued 16 November 2022 No WG43511. Crown Copyright

32.

O’Connor, H. 2016. Differences between bovine TB indicators in the IAA and the Comparison Area: First six years, 1stMay 2010 to 30th April 2016 Report extension for project OG0142.

33.

Animal and Plant Health Agency, 2023. APHA report on the delivery of badger trap and test operations on chronic TB breakdown farms in Wales in 2022. Report for project TBOG0235 (Year 6) <https://www.gov.wales/sites/default/files/publications/2023-09/bovine-tb-badger-trapping-and-testing-chronic-tb-breakdown-farms-2022.pdf>

34.

Welsh Government Consultation Document 2021. A Refreshed TB eradication programme. Issued 16 November 2022 No WG43511. Crown Copyright

35.

Department of Environment Food and Rural Affairs. 2024. Bovine TB. Accredited official statistics. Figures to December 2023 (published 20 March 2024) Updated 12 June 2024 Applies to England, Scotland and Wales. <https://www.gov.uk/government/statistics/historical-statistics-notices-on-the-incidence-of-tuberculosis-tb-in-cattle-in-great-britain-2023-quarterly/figures-to-december-2023-published-20-march-2024>

36.

Irranca-Davis, H 2024 Statement by the Cabinet Secretary for Climate Change and Rural Affairs: The future of farming in Wales – in the Senedd at 3:40 pm on 14 May 2024. https://www.theyworkforyou.com/senedd/?id=2024-05-14.5.590522.h&s=badger#g5.590549

37.

Dunnett, G., Jones, D.M. and JP McInerey (1986), Badgers and Bovine Tuberculosis – review of policy Report to the Rt Hon Michael Jopling, MP and Nicholas Edwards MP. London: Her Majesty’s Stationery Office. [see esp. appendix 11 and 12]

38.

Zuckerman, Lord (1980) Badgers, Cattle and Tuberculosis (London: Her Majesty’s

Stationery Office).

39.

Zuckerman S Monkeys Men and Missiles An autobiography 1946-88. Collins

40.

Hill, A.B “The Environment and Disease: Association or Causation?.” Proceedings of the Royal Society of Medicine 58 (1965): 295.

41.

Atkins, P.J. 2016. A History of Uncertainty Bovine Tuberculosis in Britain from 1850 to the Present. Winchester University Press

42.

Seidel, K. 2023. Milk lakes and butter mountains: The common agricultural policy. in: Leucht, B., Seidel, K. and Warlouzet, L. (ed.) Reinventing Europe: The History of the European Union, 1945 to the Present London Bloomsbury Academic. pp. 167-182

43.

Godfray, H. C. J., Donnelly, C., Hewinson, G., Winter, M., & Wood, J. (2018). Bovine TB strategy review. Department for Environment, Food and Rural Affairs. <https://www.gov.uk/government/publications/a-strategy-forachieving-bovine-tuberculosis-free-status-for-england-2018-review>

44.

Department of the Environment Food and Rural Affairs 2014.The Strategy for achieving Officially Bovine Tuberculosis Free status for England. April 2014 Crown Copyright.

45.

Cassidy A. Vermin, Victims and Disease. 2019. British Debates over Bovine Tuberculosis and Badgers [Internet]. Cham (CH): Palgrave Macmillan.

46.

Donnelly, C. A. & Nouvellet, P. The contribution of badgers to confirmed tuberculosis in cattle in high-incidence areas in England. PLoS Curr. 10, 5 (2013).

47.

Cranston R. 2018. R (on the application of LANGTON) v. Secretary of State for Environment, Food and Rural Affairs v Natural England [2018] EWHC 2190 (Admin).

2018, Judge Cranston had found that the government’s approach was not unlawful due to “a policy of maintaining a reduced badger population through supplementary culling coupled with the commitment to change tack as evidence became available”.

48.

The Stationery Office Ltd. 1992 Protection of Badgers Act 1992, LSOL, London

49.

Middlemiss and Henderson. 2022. Badger culling to control bovine TB. Letter to the

Veterinary Record, published 19th March 2022. And Correction on 21/28 May.

50.

Mcgill I, Jones M. Cattle infectivity is driving the bTB epidemic. Vet Record. 2019; 185(22), 699–700. https://pubmed.ncbi.nlm.nih.gov/31806839/.

51.

Birch, C. P. D. et al. Difference in differences analysis evaluates the effects of the badger control policy on bovine tuberculosis in England. Sci Rep 14, 4849. https://doi.org/10.1038/s41598-024-54062-4 (2024).

52.

Macdonald, D. 2023 A Commentary on Current Policy. A preamble to Badger Trust’s report ‘Tackling Bovine TB Together’: Towards Sustainable, Scientific and Effective bTB Solutions. Badger Trust publication. Brighton.

53.

King, D. 2007 Bovine tuberculosis in cattle and badgers. A report by the Chief Scientific Adviser, Sir David King. Report to the Secretary of State, Defra on 30 July 2007

54.

Langton T E S. Badger culling and bovine TB in cattle: a re-evaluation of proactive culling benefit in the randomized badger culling trial. J Dairy Vet Sci. 2019; 12(1):555826. https://juniperpublishers.com/jdvs/pdf/JDVS.MS.ID.555826.pdf.

55.

Donnelly CA, Woodroffe R, Cox DR, Bourne FJ, Cheeseman CL, Clifton-Hadley RS, et al. Positive and negative effects of widespread badger culling on tuberculosis in cattle. Nature. 2006 Feb;439(7078):843–6.

56.

Smaldino Paul E. and McElreath Richard 2016The natural selection of bad science R. Soc. Open Sci.3160384 <http://doi.org/10.1098/rsos.160384>
