## Supplemental Material 3 for "Randomised Badger Culling Trial lacks evidence for proactive badger culling effect on tuberculosis in cattle: comment on Mills et al. 2024, Parts I & II"

### Further details on statistical models and issues

**Details of errors in Mills et al. 2024ab(1,2). Documented in the main text.**

**Models in Mills et al 2024a(1)**

1. Model a.1.

This model in table 2b in Mills et al. 2024a(1) is said to be identical to model rs in Torgerson et al. 2024(3) . This is confirmed by page 2 of the supplementary information in Mills et al. 2024a(1).

Model a.1. is described as varying intercepts for triplets and covariates of culling effect, historical 3-year incidence of cattle TB, and baseline herds-at-risk. It was also a negative binomial model.

The code given in github (MillsCathal. 2024 Rbct_analyses. for this model is:GitHub. See https://github.com/cathalmills/rbct_analyses/) (4)

```
rs <- stan_glm.nb(Incidence~Treatment+A+B+C+D+E+F+G+H+I+log(Hist3yr)+log(Baseline),
        data = rbctconf, prior_intercept=normal(0, 10),
        diagnostic_file = file.path(tempdir(), "df.csv"),refresh=0
```

The authors of the code say one of the problems with this model is the strongly informative prior distribution for dispersion.

Note here that stan_glm.nb automatically gives this prior if the prior is not defined by the user.

But model rs in Torgerson et al. 2024(3) was NOT a negative binomial model. It was a Poisson model, and the original model of Donnelly et al. in the Bayesian paradigm, to act as a starting point for model development and selection. The code of the model in Torgerson et al. 2024(3) is:

```
rs <- stan_glm(Incidence~Treatment+A+B+C+D+E+F+G+H+I+log(Hist3yr)+log(Baseline),
        data = rbctconf, prior_intercept=normal(0, 10), poisson,
        diagnostic_file = file.path(tempdir(), "df.csv"),refresh=0
```

Here, we used stan_glm and defined the Poisson family in the code. Mills et al.(1,4) used stan_glm.nb, which is defined by use of the negative binomial comparison.

Consequently, model a.1 in Mills et al. 2024a(1) and model rs from Torgerson et al. 2024 (3) give different results.

2. Model a.2

This is model a.1. improved (table 2a Mills et al, 2024a)(1), and is trying to improve a model we never used in Torgerson et al. 2024 and is thus irrelevant and misleading.

3. Model c.1

table 2a Mills et al. 2024a.(1)

Model description: “herd years at risk as an offset and no varying intercepts for triplets”.

The code for model c.1 given in github(4) (line 1871) is:

```
rs1aB<-stan_glm.nb(Incidence~Treatment+log(hdyrsrisk)+log(Hist3yr),
                offset = log(hdyrsrisk),
                prior_intercept=normal(0,10), prior=normal(0,10),
                data = rbctconf, refresh=0)
```

This should have been equivalent to model rs1aB in Torgerson et al. 2024 (3).

Note that, in the code, the authors of the code give log(hdyrsrisk) as both an offset variable and an explanatory variable.

This effectively ignores the offset as log(hdyrsrisk) and offset(log(hrdrsrsik)) can be combined. Suppose the former has some unknown parameter n, whilst for the offset it is fixed at 1

Therefore,

n*log(hdyrsrisk) = log(hdyrsrisk^n)

1*log(hdyrsrisk) + log(hdyrsrisk^n) = log(hdyrsrisk*hdyrsrisk^n) = (n+1)log(hdyrsrisk)

So, by having a variable as both an offset and an exploratory variable, the parameter of variable log(hdyrsrisk) is just shifted by 1.

Thus, with the code in Mills et al. 2024a (1,4), the parameter values of log(hdyrsrisk) = -0.518
Running the same code, but without offset=log(hdyrsrisk), the parameter value of log(hdyrsrisk) = 0.482.

Now -0.518+1 = 0.482

All other parameters remain the same. So, the code used in Mills et al. 2024a(1,4) did not have an offset, and therefore, has no equivalence to model rs1aB in Torgerson et al. 2024 (3)

3. Model c.2.

(c1. Improved) table 2b also has the same offset error as model c.1.

4. Model e.

From Table 2b.

Model e is described as “Poisson with herds-at-risk as an offset”

The code for model e in Mills et al. is given in GitHub (MillsCathal. 2024 Rbct_analyses. GitHub. See https://github.com/cathalmills/rbct_analyses/) (4).

Model e is in line 1406 in the GitHub code (4). This was confirmed in email correspondence with Mills.

The code given for model e is:

```
set.seed(150499)
rs_pois <- rstanarm::stan_glm(Incidence~Treatment+A+B+C+D+E+F+G+H+I+log(Hist3yr),
                 data = rbctconf, family = "poisson",
                 offset = log(Baseline),
                 prior = normal(0, 1),
                 prior_intercept = normal(0, 2),
                 diagnostic_file = file.path(tempdir(), "df.csv"),refresh=0)
```

When this model is run, the summary file to give the parameter estimates is:

```
summary(rs_pois, digits=3, probs=c(0.025,0.5,0.975))

Estimates:
                      mean   sd     2.5%   50%    97.5%
(Intercept)          -3.383  0.498 -4.354 -3.382 -2.406
TreatmentProactive   -0.146  0.074 -0.285 -0.147 -0.001
A1                    0.327  0.165  0.006  0.326  0.651
B1                    0.220  0.158 -0.076  0.219  0.535
C1                    0.243  0.147 -0.043  0.241  0.538
D1                   -0.005  0.166 -0.333 -0.005  0.320
E1                    0.204  0.155 -0.086  0.202  0.503
F1                   -0.391  0.165 -0.709 -0.395 -0.070
G1                   -0.001  0.150 -0.298  0.000  0.290
H1                   -0.051  0.170 -0.377 -0.053  0.286
I1                   -0.277  0.180 -0.642 -0.277  0.079
log(Hist3yr)          0.731  0.158  0.419  0.732  1.038
```

The effect of badger culling (Treatment proactive), in terms of % reduction, can be found by taking the exponents of the parameter values.

Mean reduction = 1-exp(-0.145) = 0.136 = 13.6%

Likewise, the 95% credible intervals are 0.10%, 24.7%.

Mills et al. 2024a(1) cite the reduction using this model to be 18.0% with 95% CrI 5% reduction to 28.9% reduction.

Mill et al. 2024a(1) also give ELPD LOO as -86.5. This agrees with the actual ELPD LOO. When running the code, the ELPD LOO is:

```
loo(rs_pois, k_threshold = 0.7)

 Estimate   SE
elpd_loo    -86.5  9.5
p_loo        19.9  6.6
looic       172.9 18.9
```

So, the results reported in Mills et al.  2024a(1)  , Table 2b do not correspond to those when the code is run. Mills et al. 2024a(1)  used these results to construct Figure 1. So, therefore, Figure 1 is incorrect.

<u>5. Other offset code issues</u>

The issue of the offset variable being also coded as an explanatory variable is repeated many times in the code for the post trial period.

This can be shown in code from Github (4) commencing lines: 1421; 1436; 1898; 1913; 1943; 1960.

**Models from Mills et al 2024b**.(2)

<u>6. Model a.1.</u>

Model description: varying intercepts for triplets and covariates of culling effect, historical 3-year incidence and baseline herds at risk.

This has the same issues as error number 1 (above) in Mills et al. 2024a(1) . So, it is invalid. This also applies to Model a.2 (a.1. improved)

<u>7. model c.1.</u>

Model description: using herd-years-at-risk as an offset and no varying intercepts for triplets.

The code from Mills et al. 2024a (GitHub, line 1931) (4) is:

```
rs1_outside_aB<-stan_glm.nb(Incidence~Treatment+log(hdyrsrisk)+log(Hist3yr),
                        offset = log(hdyrsrisk),
                        prior_intercept=normal(0,10), prior=normal(0,10),
                        data = rbctconf_outside, refresh=0)
```

Note that Mills et al.(4) had log(hdyrsrisk) defined as an explanatory variable and offset variable, so effectively there was no offset and the conclusions are invalid.

8. model c.2.

(c1 improved)

Code given in Mills et al. 2024b. GitHub, Line 1939. (4)

```
rs1_outside_aB_improved<-stan_glm.nb(Incidence~Treatment+log(hdyrsrisk)+log(Hist3yr),
                                prior_intercept = normal(0, 5),
                                prior = normal(0, 1),
                                prior_aux= cauchy(0, 5),
                                offset = log(hdyrsrisk),
                                data = rbctconf_outside, refresh=0)
```

Again, the same issue of log(hdyrsrisk) as both an offset and explanatory variable and the conclusions are invalid.

9. Model e,

Mills et al. 2024b(2)

Model described as: Poisson with baseline herds at risk as an offset.

Code from Mills et al. 2024b, github (4) line 1559:

```
rs_pois_outside <- stan_glm(Incidence~Treatment+A+B+C+D+E+F+G+H+I+log(Hist3yr)
+log(Baseline),
                        data = rbctconf_outside, family = "poisson",
                        offset = log(Baseline),
                        prior = normal(0, 1),
                        prior_intercept = normal(0, 2),
                        diagnostic_file = file.path(tempdir(), "df.csv"),refresh=0)
```

Again, the same issue this time log(Baseline) as both an offset and explanatory variable. This is effectively the nature model in Bayesian paradigm.

10. Offset variable

The issue of the offset variable being also coded as an explanatory variable is repeated many times in the code for the post trial period. This can be shown in the Github code (4) commencing on lines 1582; 1807; 1953; 1963; 1900; 2026; 2042. Therefore, all conclusions assuming an offset for these models are invalid.

#### Posterior Predictive Checks

This compares the original Poisson GLM from Donnelly et al.(5) to the optimal model reported in Torgerson et al.(3) for confirmed herd breakdowns from initial cull until 4 September 2005 within proactive culling areas of the RBCT. Here, we illustrate Model 1, the model used in Donnelly et al. and defended as robust by Mills et al. (Figure 1) and compare directly with Model 8, the most parsimonious of the (generalized) Poisson models examined in Torgerson et al. (Figure 2).

Note that from Model 8, the incidence (counts, in this case) is independent of culling. For each model we present six simulations that demonstrate that there are inconsistent results between different simulation. Also, the visual check suggests no obvious differences between the two models. Note that Mills et al. claim that, for Model 1, "a visual posterior predictive check indicates that the model's posterior predictive distribution resembles the observed data", whilst for model 8, it is claimed that "the posterior predictive check implies potential model misfit due to systematic discrepancies between model-predicted data and confirmed incidence".

Likewise for model 11, Mills et al, 2024b(2) claim " *the posterior predictive check the systematic discrepancies between the model-based predictions and the confirmed incidence hint at model misspecification for a model which assumes identical triplets*". Figure 3 demonstrates the "misspecification" reported in Mills et al 2024b, compared to a simulation, from a different random seed, where the "misspecification" is less convincing.

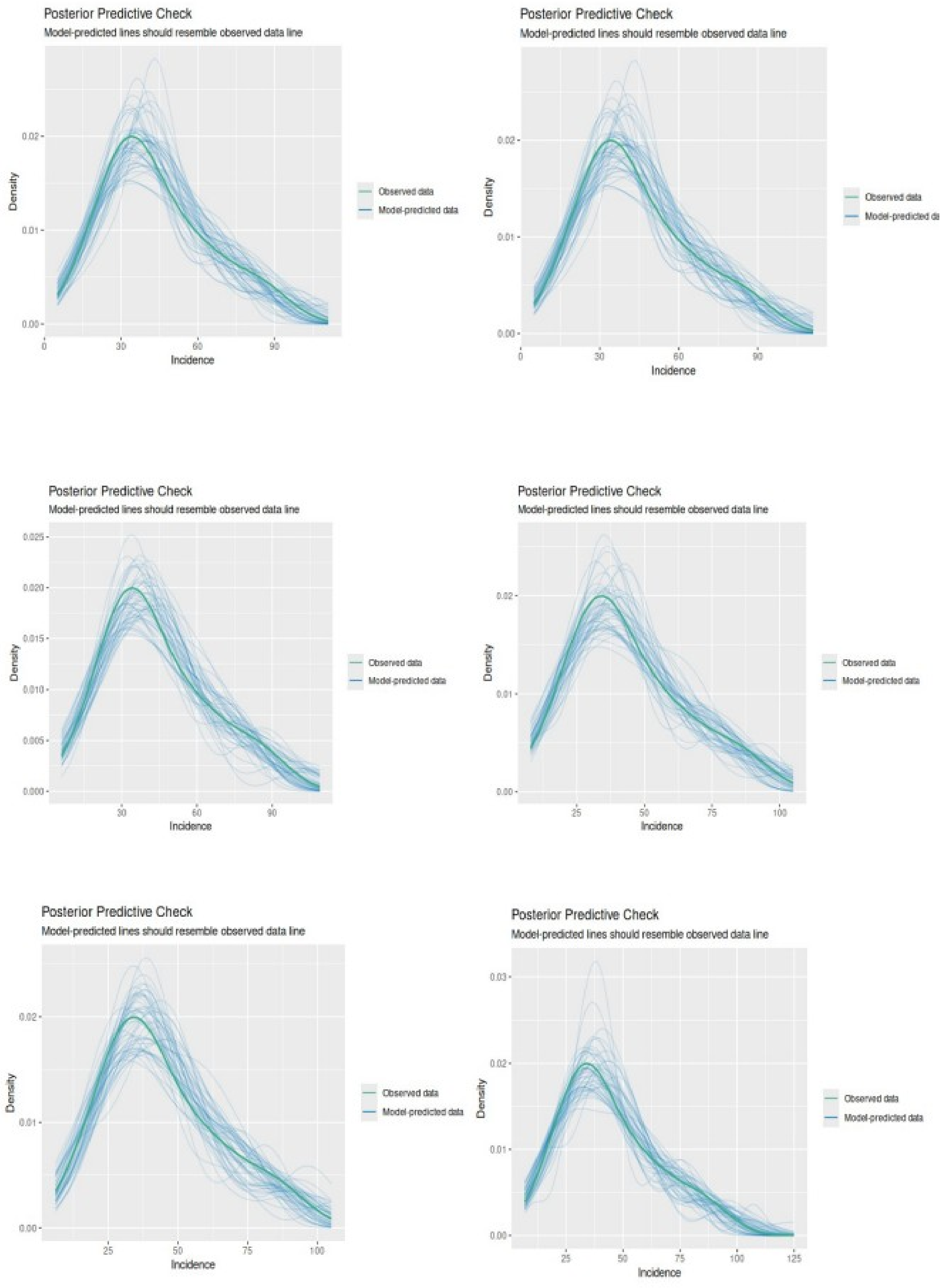


Figure 1. Visual posterior predictive check of Model 1. Note the inconsistencies of the model-predicted data from the six simulations.

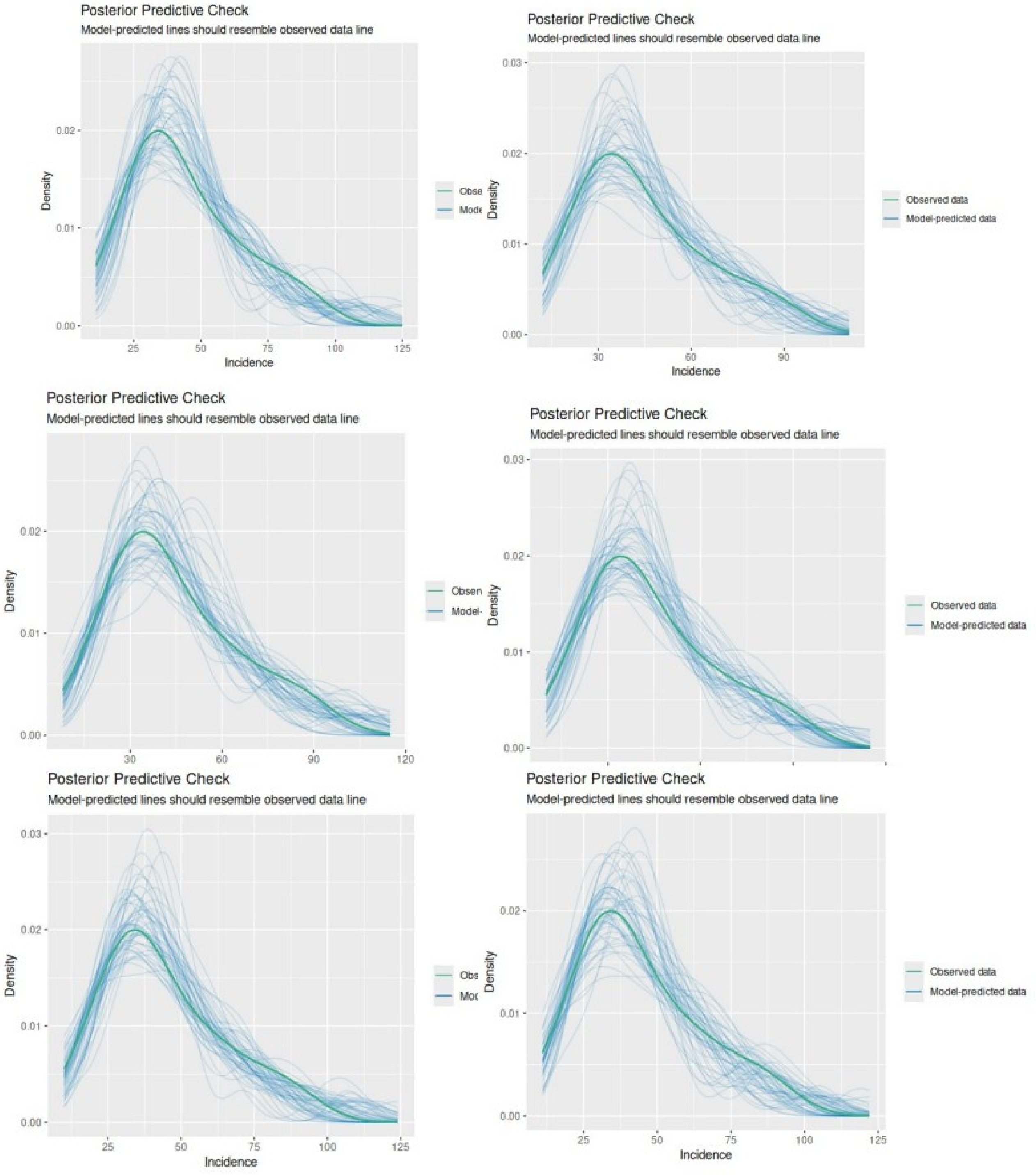


Figure 2. Visual posterior predictive check of Model 8.

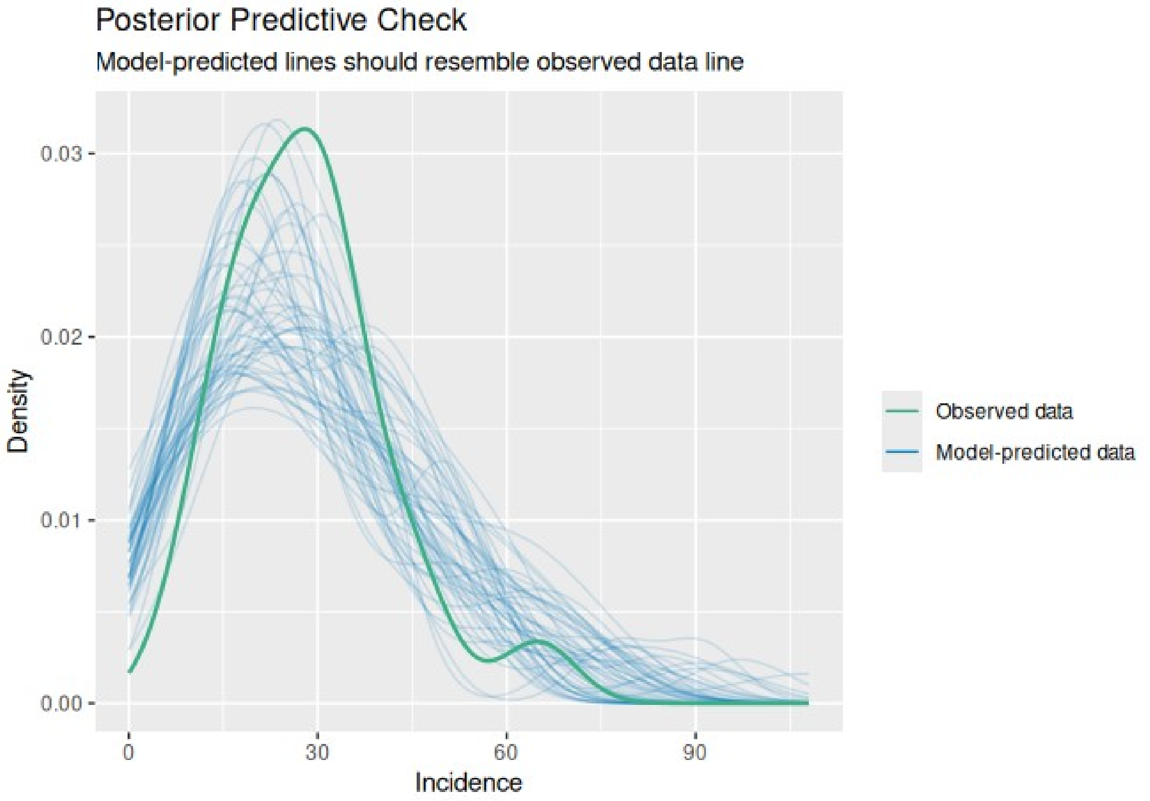


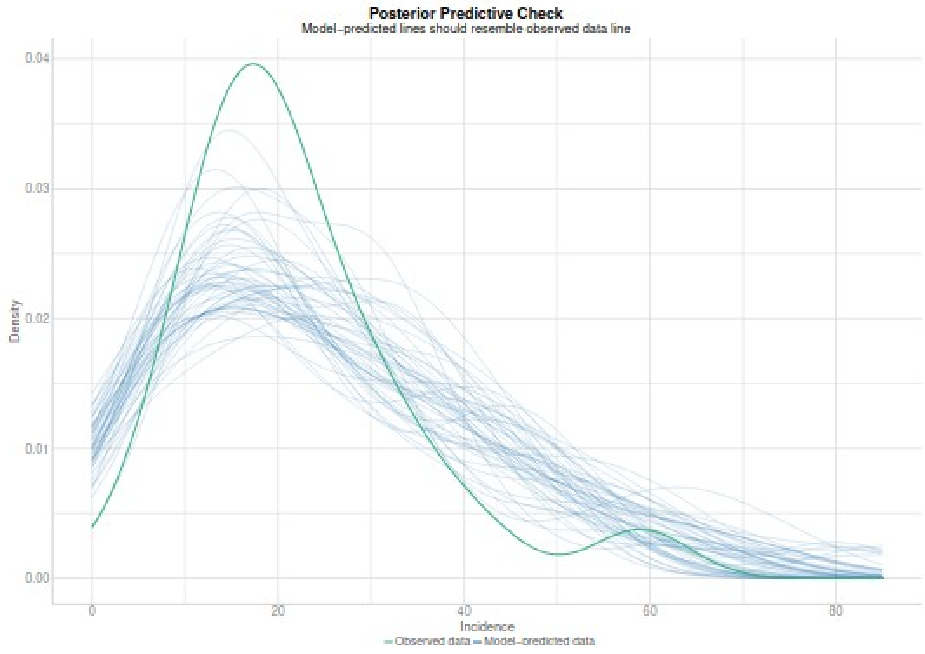


Figure 3. Visual posterior predictive check of model 11 from Mills et al 2024b(2). Top is the simulation as in the code in the supplementary information the seed is set. =set.seed(94060000) pp_check(model11). Bottom is the visual PPC from the supplementary material of Mills et al. 2024b (Figure SI 5)(2). The important point here is that the PPC can give markedly different results with a different random seed. Therefore it seems to be unreliable and allows for selection of a result to fit the narrative.

#### Check of Residuals

This compares the original Poisson GLM from Donnelly et al.(5) to the optimal model reported in Torgerson et al.(3) for confirmed herd breakdowns from initial cull until 4 September 2005 within proactive culling areas of the RBCT.

Here, quantile deviation is detected in the simulated-residual-versus-predicted plot in the Donnelly model. Furthermore, significant results are attained for uniformity and deviations from uniformity, but the test of overdispersion or underdispersion yields a significant result (*p*-value 0.001). For the model in Torgerson et al. 2024, no issues are detected (Figure 3).

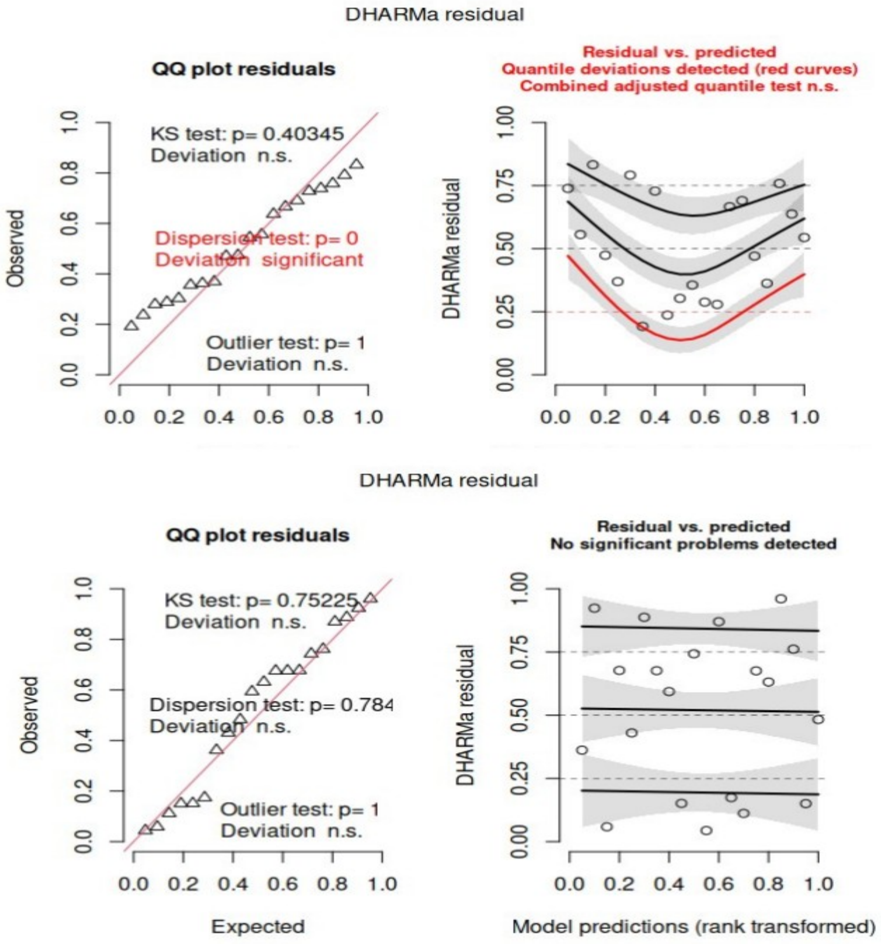


Figure 3. For confirmed herd breakdowns from initial cull until 4 September 2005 within proactive culling areas of the RBCT, we present the residual plot of the Model in Donnelly et al.(5), which claimed an effect of culling (top), compared to residuals of the most parsimonious model in Torgerson et al. 2024(3), where there was no effect of culling.

#### Statistical appraisal and diagnostic techniques

Table 1 gives the statistical appraisal and diagnostic techniques to compare the original Poisson GLM from Donnelly et al.(5) (Model 1) to the optimal model reported in Torgerson et al.(3) (Model 8) for confirmed herd breakdowns from initial cull until 4 September 2005 within proactive culling areas of the RBCT. For rigour, we also include Model 1a, which is identical to Model 1, but with log(herd years at risk) as the explanatory variable instead of log(number of herds at risk). This is to ensure direct comparison with Model 8 (i.e., the same data). But note that there is no change in the diagnostics between Models 1 and Model 1a, while Mills et al. defend model 1 as "robust".

Table 1. Statistical appraisal and diagnostic techniques to compare the original Poisson GLM from Donnelly et al.(5)(Model 1) to the optimal model reported in Torgerson et al.(3)(Model 8) for confirmed herd breakdowns from initial cull until 4 September 2005 within proactive culling areas of the RBCT.

| | BIC# | AICc# | LOO# | Outliers | Number of estimated parameters# | Parameter for exposure (95% CI)* |
|---|---|---|---|---|---|---|
| Model 1 | 155.2 | 203.0 | 9.97 | None | 13 | 0.05 (-0.44,0.53 |
| Model 1a | 155.2 | 203.0 | 9.97 | None | 13 | 0.05 (-0.44,0.53) |
| Model 8 | 155.5 | 154.2 | 8.81 | None | 3+1§ | 0.51 (0.32,0.70) |

#Lower is better

§Three explanatory variables and one to model overdispersion.

* A parameter for exposure that is not significantly different from zero indicates no variation of incidence (counts) with samples size or time-at-risk. This is implausible and is, therefore, a strong indicator of misspecification.
