## Supplementary material R code for "Randomised Badger Culling Trial lacks evidence for proactive badger culling effect on tuberculosis in cattle: comment on Mills et al. 2024, Parts I & II"

---
title: "Supplementary material R code"
output: pdf_document
date: "2024-09-11"
---

```{r setup, include=FALSE}
knitr::opts_chunk$set(echo = TRUE)
```

  
## Comment on  Mills et al. 2024ab: "Extensive re-evaluation the Randomised Badger Culling Trial (RBCT) fails to show evidence of effects of widespread badger culling on tuberculosis in cattle" 

## Authors: Torgerson PR, Hartnack S, Rasmussen P, Lewis F, O’Donnell P, and TES Langton


Code and explanatory text is given for all models in the text. Where necessary it is important to refer to the models and model code given in 
Mills et al. 2024ab (<https://royalsocietypublishing.org/doi/10.1098/rsos.240386> and <https://doi.org/10.1098/rsos.240385>) and Torgerson et al. 2024 (<https://www.nature.com/articles/s41598-024-67160-0>) and their supplementary materials for full understanding.

```{r, message = FALSE, warning = FALSE}
library(glmmTMB)
library(MuMIn)
library(rstanarm)
library(bridgesampling)
library(DHARMa)
library(readxl)
library(AER)
library(vcd)
library(performance)
library(cowplot)
library(rstan)
library(boot)
library(olsrr)
library(lme4)
library(caret)
library(bayesplot)
library(gridExtra)
library(vctrs)
library(purrr)
library(loo)
library(plyr)
```

\newpage
Data as per Donnelly et al, Nature (2006) <http://dx.doi.org/10.1038/nature04454>

# Codebook

| Variable   | Value     |
|------------|-----------|
| Triplet    | This is   |
| Treatment  | This is   |
| Incidence  | This is   |
| Baseline   | This is   |
| Hist3yr    | This is   |
| hdyrsrisk  | This is   |


```{r}
# Incidence data from initial cull until September 2005
rbctconf <- data.frame(read_xlsx("confirmed_vetnet.xlsx"))

rbctconf[c(1:6)] <- lapply(rbctconf[c(1:6)], factor)
```

```{r}
# Relevel triplets and treatment (proactive culling vs survey-only)
# to correspond to Mills et al. and Torgerson et al. and thus, enable model review

rbctconf$Triplet <- relevel(factor(rbctconf$Triplet), ref="J") 
rbctconf$Treatment <- relevel(factor(rbctconf$Treatment), ref="Survey-only")
```

\newpage
# Table 1
## Model 1. Original Poisson GLM as per Donnelly et al, Nature (2006). 
<http://dx.doi.org/10.1038/nature04454>. 
```{r}
# Fit to initial period
model1 <- glm(Incidence ~ Treatment + Triplet + log(Hist3yr) + log(Baseline), 
            family = poisson, data=rbctconf)

summary(model1)
```

### Table 1, model1: Estimated effect of culling (95% CI)
```{r}
round((exp(coef(model1))[2] - 1),3)*100   # point estimate
round((exp(coefci(model1))[2,1]-1),3)*100 # lower 95% CI
round((exp(coefci(model1))[2,2]-1),3)*100 # upper 95% CI
```

### Table 1, model1: BIC and AICc
```{r}
round(BIC(model1),1)
round(AICc(model1),1)
```

### Table 1, model1: LOOCV RMSE 
Leave-One-Out Cross-Validation and Root Mean Squared Error, 
a metric used to evaluate the performance of a model.
```{r}
rmse <- numeric()
actual_vs_predicted <- data.frame(Actual = numeric(), Predicted = numeric())
for (i in 1:nrow(rbctconf)) {
  # Exclude the ith row
  test_data <- rbctconf[i, ]
  train_data <-rbctconf[-i, ]
  model <-glm(Incidence ~ Treatment + Triplet + log(Hist3yr) + log(Baseline),
              family = poisson, data = train_data)
  predictions <- predict(model, newdata = test_data, type="response")
  rmse[i] <- sqrt(mean((test_data$Incidence - predictions)^2))
  actual_vs_predicted <- rbind(actual_vs_predicted, 
                               data.frame(Actual = test_data$Incidence, Predicted = predictions))
}

rmse1<-actual_vs_predicted %>%
  mutate(rmse = sqrt((Actual - Predicted)^2))

round(mean(rmse1$rmse),2)
```

### Table 1, model1: Number of explanatory parameters
```{r}
length(coef(model1))
```

### Table 1, model1: Parameter for exposure (95% CI)
```{r}
# this is on the log scale
round(coef(model1)[13],2)     # point estimate
round(coefci(model1)[13,1],2) # lower 95% CI
round(coefci(model1)[13,2],2) # upper 95% CI
```

\newpage
In the text there is also mentionned this model for other time periods.
```{r}
### Incidence data from first follow-up cull until September 2005
rbctconf_follow <- data.frame(read_xlsx("confirmed_vetnet_FOLLOW_UP.xlsx"))
rbctconf_follow[c(1:6)] <- lapply(rbctconf_follow[c(1:6)], factor)

# relevel
rbctconf_follow$Triplet <- relevel(factor(rbctconf_follow$Triplet), ref="J")
rbctconf_follow$Treatment <- relevel(factor(rbctconf_follow$Treatment), 
                                     ref="Survey-only")

#Follow-Up Period
model1_follow<-glm(Incidence ~ Treatment + Triplet + log(Hist3yr) 
                   + log(Baseline), 
                   family = poisson, data=rbctconf_follow)

summary(model1_follow)

BIC(model1_follow)

AICc(model1_follow)


### Incidence data from post-trial period 
### (1 year after last proactive cull until 2013) 
rbctconf_after <- data.frame(read_xlsx("confirmed_rbctconf_after_trial.xlsx"))
rbctconf_after[c(1:6)] <- lapply(rbctconf_after[c(1:6)], factor)

# relevel
rbctconf_after$Triplet <- relevel(factor(rbctconf_follow$Triplet), ref="J")
rbctconf_after$Treatment <- relevel(factor(rbctconf_follow$Treatment), ref="Survey-only")

#Post trial  period
model1_after<-glm(Incidence ~ Treatment + Triplet + log(Hist3yr) +log(Baseline), 
                  family = poisson, data=rbctconf_after)

summary(model1_after)

BIC(model1_after)

AICc(model1_after)
```

\newpage
## Model 3: Original GLM in generalized Poisson form
### Table 1, model3
```{r}
model3 <- glmmTMB(Incidence ~ Treatment + Triplet + log(Hist3yr) + log(Baseline), 
                family = genpois, data=rbctconf)

summary(model3)
```

### Table 1, model3: Estimated effect of culling (95% CI)
```{r}
round((exp(confint(model3, level = 0.95)[2,3])-1),3)*100 # point estimate
round((exp(confint(model3, level = 0.95)[2,1])-1),3)*100 # lower 95% CI
round((exp(confint(model3, level = 0.95)[2,2])-1),3)*100 # upper 95% CI
```

### Table 1, model3: BIC and AICc
```{r}
round(BIC(model3),1)
round(AICc(model3),1)
```

### Table 1, model3: LOOCV RMSE 
Leave-One-Out Cross-Validation and Root Mean Squared Error, a metric used to evaluate the performance of a model.
```{r}
rmse <- numeric()
actual_vs_predicted <- data.frame(Actual = numeric(), Predicted = numeric())
for (i in 1:nrow(rbctconf)) {
  # Exclude the ith row
  test_data <- rbctconf[i, ]
  train_data <-rbctconf[-i, ]
  model <-glmmTMB(Incidence ~ Treatment + Triplet + log(Hist3yr) + log(Baseline), 
                family = genpois, data = train_data)
  predictions <- predict(model, newdata = test_data, type="response")
  rmse[i] <- sqrt(mean((test_data$Incidence - predictions)^2))
  actual_vs_predicted <- rbind(actual_vs_predicted, 
                               data.frame(Actual = test_data$Incidence, 
                                          Predicted = predictions))
}

rmse1<-actual_vs_predicted %>%
  mutate(rmse = sqrt((Actual - Predicted)^2))

round(mean(rmse1$rmse),2)
```

### Table 1, model3: Number of explanatory parameters
```{r}
length(fixef(model3)$cond) # plus an additional parameter for overdispersion
```

### Table 1, model3: Parameter for exposure (95% CI)
```{r}
# this is on the log scale
round(confint(model3, level = 0.95)[13,3],2) # point estimate
round(confint(model3, level = 0.95)[13,1],2) # lower 95% CI
round(confint(model3, level = 0.95)[13,2],2) # upper 95% CI
```

\newpage
In the text there is also mentionned this model for other time periods.
```{r}
#Follow-Up Period
model3_follow <- glmmTMB(Incidence ~ Treatment + Triplet + log(Hist3yr)
                         + log(Baseline), 
                       family = genpois, data=rbctconf_follow)

summary(model3_follow)

BIC(model3_follow)

AICc(model3_follow)

#Post trial  period
model3_after <- glmmTMB(Incidence ~ Treatment + Triplet + log(Hist3yr)
                        + log(Baseline), 
                      family = genpois, data=rbctconf_after)

summary(model3_after)

BIC(model3_after)

AICc(model3_after)
```

\newpage
## Model 4: Generalized Poisson (without any culling effect)
### Tabe 1, model4
```{r}
model4 <- glmmTMB(Incidence ~ log(Hist3yr) + log(Baseline), family = genpois, 
                  data=rbctconf)

summary(model4)

confint(model4)
```

### Table 1, model4: BIC and AICc
```{r}
round(BIC(model4),1)
round(AICc(model4),1)
```

### Table 1, model4: LOOCV RMSE 
Leave-One-Out Cross-Validation and Root Mean Squared Error, 
a metric used to evaluate the performance of a model.
```{r}
rmse <- numeric()
actual_vs_predicted <- data.frame(Actual = numeric(), Predicted = numeric())
for (i in 1:nrow(rbctconf)) {
  # Exclude the ith row
  test_data <- rbctconf[i, ]
  train_data <-rbctconf[-i, ]
  model <- glmmTMB(Incidence ~ log(Hist3yr) + log(Baseline), family = genpois, 
                   data = train_data)
  predictions <- predict(model, newdata = test_data, type="response")
  rmse[i] <- sqrt(mean((test_data$Incidence - predictions)^2))
  actual_vs_predicted <- rbind(actual_vs_predicted, 
                               data.frame(Actual = test_data$Incidence, 
                                          Predicted = predictions))
}

rmse1<-actual_vs_predicted %>%
  mutate(rmse = sqrt((Actual - Predicted)^2))

round(mean(rmse1$rmse),2)
```

### Table 1, model4: Number of explanatory parameters
```{r}
length(fixef(model4)$cond) # plus an additional parameter for overdispersion
```

### Table 1, model4: Parameter for exposure (95% CI)
```{r}
# this is on the log scale
round(confint(model4, level = 0.95)[3,3],2) # point estimate
round(confint(model4, level = 0.95)[3,1],2) # lower 95% CI
round(confint(model4, level = 0.95)[3,2],2) # upper 95% CI
```

\newpage
In the text there is also mentionned this model for other time periods.
```{r}
#Follow-Up Period
model4_follow <- glmmTMB(Incidence ~ log(Hist3yr) + log(Baseline), 
                       family = genpois, data=rbctconf_follow)


summary(model4_follow)

BIC(model4_follow)

AICc(model4_follow)


#Post trial  period
model4_after <- glmmTMB(Incidence ~ log(Hist3yr) + log(Baseline), 
                      family = genpois, data=rbctconf_after)

summary(model4_after)

BIC(model4_after)

AICc(model4_after)
```


## Model 8: Generalized Poisson with herd-years-at-risk covariate without any culling effect
### Table 1, model8
```{r}
model8 <- glmmTMB(Incidence ~ log(Hist3yr) + log(hdyrsrisk), family = genpois, 
                  data=rbctconf)

summary(model8)
```

### Table 1, model8: BIC and AICc
```{r}
round(BIC(model8),1)
round(AICc(model8),1)
```

### Table 1, model8: LOOCV RMSE 
Leave-One-Out Cross-Validation and Root Mean Squared Error, 
a metric used to evaluate the performance of a model.
```{r}
rmse <- numeric()
actual_vs_predicted <- data.frame(Actual = numeric(), Predicted = numeric())
for (i in 1:nrow(rbctconf)) {
  # Exclude the ith row
  test_data <- rbctconf[i, ]
  train_data <-rbctconf[-i, ]
  model <- glmmTMB(Incidence ~ log(Hist3yr) + log(hdyrsrisk), family = genpois, 
                   data = train_data)
  predictions <- predict(model, newdata = test_data, type="response")
  rmse[i] <- sqrt(mean((test_data$Incidence - predictions)^2))
  actual_vs_predicted <- rbind(actual_vs_predicted, 
                               data.frame(Actual = test_data$Incidence, 
                                          Predicted = predictions))
}

rmse1<-actual_vs_predicted %>%
  mutate(rmse = sqrt((Actual - Predicted)^2))

round(mean(rmse1$rmse),2)
```

### Table 1, model8: Number of explanatory parameters
```{r}
length(fixef(model8)$cond) # plus an additional parameter for overdispersion
```

### Table 1, model8: Parameter for exposure (95% CI)
```{r}
# this is on the log scale
round(confint(model8, level = 0.95)[3,3],2) # point estimate
round(confint(model8, level = 0.95)[3,1],2) # lower 95% CI
round(confint(model8, level = 0.95)[3,2],2) # upper 95% CI
```

\newpage
In the text there is also mentionned this model for other time periods.
```{r}
#Follow-Up Period
model8_follow <- glmmTMB(Incidence ~ log(Hist3yr) + log(hdyrsrisk), 
                       family = genpois, data=rbctconf_follow)

summary(model8_follow)

BIC(model8_follow)

AICc(model8_follow)


#Post trial  period
model8_after <- glmmTMB(Incidence ~ log(Hist3yr) + log(hdyrsrisk), 
                      family = genpois, data=rbctconf_after)

summary(model8_after)

BIC(model8_after)

AICc(model8_after)
```

## Null model without any predictors
### Table 1, null model
```{r}
model.null <- glmmTMB(Incidence ~ 1, family = genpois, 
                  data=rbctconf)
summary(model.null)
```

### Table 1, model null: BIC and AICc
```{r}
round(BIC(model.null),1)

round(AICc(model.null),1)
```

### Table 1, model null: LOOCV RMSE 
Leave-One-Out Cross-Validation and Root Mean Squared Error, a metric used to evaluate the performance of a model.
```{r}
rmse <- numeric()
actual_vs_predicted <- data.frame(Actual = numeric(), Predicted = numeric())
for (i in 1:nrow(rbctconf)) {
  # Exclude the ith row
  test_data <- rbctconf[i, ]
  train_data <-rbctconf[-i, ]
  model <- glmmTMB(Incidence ~ 1, family = genpois, 
                   data = train_data)
  predictions <- predict(model, newdata = test_data, type="response")
  rmse[i] <- sqrt(mean((test_data$Incidence - predictions)^2))
  actual_vs_predicted <- rbind(actual_vs_predicted, 
                               data.frame(Actual = test_data$Incidence, 
                                          Predicted = predictions))
}

rmse1<-actual_vs_predicted %>%
  mutate(rmse = sqrt((Actual - Predicted)^2))

round(mean(rmse1$rmse),2)
```

### Table 1, model8: Number of explanatory parameters
```{r}
length(fixef(model.null)$cond) # plus an additional parameter in generalized Poisson model to model overdispersion
```

\newpage
# Table 2: Bayesian analysis
## Model a.1 as coded by Mills et al.
Note the stan_glm.nb function, which is modelling a negative binomial regression
```{r}
modela1 <- stan_glm.nb(Incidence~Treatment+A+B+C+D+E+F+G+H+I+log(Hist3yr)+log(Baseline),
                  data = rbctconf, prior_intercept=normal(0, 10),
                  diagnostic_file = file.path(tempdir(), "df.csv"),refresh=0)

summary(modela1, digits=5, probs=c(0.025,0.5,0.975))
#This code allows the software to provide minimally informative priors automatically

prior_summary(modela1)
#prior for auxiliary (reciprocal dispersion) ~ exponential(rate1)
#This strongly informative prior forces a negative binomial model,
#when the data is consistent with a poisson.
#This was not the model rs proposed by Torgerson et al.
```


## Table 2, Modelrs as coded by Torgerson et al.
This is the properly coded model proposed by Torgerson et al,
with weakly informative priors, note the stan_glm function and poisson family defined
```{r}
modelrs<-stan_glm(Incidence~Treatment+A+B+C+D+E+F+G+H+I+log(Hist3yr)+log(Baseline),poisson,
                     data = rbctconf, prior_intercept=normal(0, 10),
                     diagnostic_file = file.path(tempdir(), "df.csv"),refresh=0)

summary(modelrs, digits=5, probs=c(0.025,0.5,0.975))
```


## Table 2, Model b.1.
```{r}
modelb1 <- stan_glm.nb(Incidence ~ Treatment + log(hdyrsrisk) + log(Hist3yr),
                  prior_intercept=normal(0,10), prior=normal(0,10),
                  data=rbctconf,diagnostic_file =file.path(tempdir(),"df.csv"),
                  refresh=0)

summary(modelb1, digits=5, probs=c(0.025,0.5,0.975))
#Effect size does not appear to align with that reported as model rsB1 in Torgerson et al.
```

## Table 2 model c.1. in table 2 in the text
```{r}
#Code as reported by Mills et al. Note log(hdyrsrisk) is both an explanatory
#variable and an offset variable
modelc1 <- stan_glm.nb(Incidence ~ Treatment + log(hdyrsrisk) + log(Hist3yr),
                   offset = log(hdyrsrisk),
                   prior_intercept=normal(0,10), prior=normal(0,10),
                   data = rbctconf, refresh=0)

summary(modelc1, digits=5, probs=c(0.025,0.5,0.975))

loo(modelc1)

#Summary gives parameter value for explanatory variable log(hdyrsrisk)
#Parameter value for Treatment and LOO ELPD align with values reported
#in table 2a in Mills et al.


#The correct code so that log(hdyrsrisk) is an offset variable is:
modelc1 <- stan_glm.nb(Incidence ~ Treatment + log(Hist3yr),
                     offset = log(hdyrsrisk),
                     prior_intercept=normal(0,10), prior=normal(0,10),
                     data = rbctconf, refresh=0)

#Same coding issue for follow up and post trial periods in code written by Mills et al.

summary(modelc1, digits=5, probs=c(0.025,0.5,0.975))

#Coding the offset correctly gives different parameter estimates

loo(modelc1)
#and different LOO ELPD
```


## Table 2, Model c2
```{r}
#Code used by Mills et al.
set.seed(98356)
modelc2 <- stan_glm.nb(Incidence ~ Treatment + log(hdyrsrisk) + log(Hist3yr),
                            prior_intercept = normal(0, 1, autoscale = TRUE),
                            prior = normal(0, 1, autoscale = TRUE),
                            prior_aux= cauchy(0, 5, autoscale = TRUE),
                            offset = log(hdyrsrisk),
                            data = rbctconf, refresh=0)

#As with model c1, Mills et al. coded the log(hdyrsrisk) as both an explanatory
#and offset variable

#Correct code is:
modelc2 <- stan_glm.nb(Incidence ~ Treatment + log(Hist3yr),
                     prior_intercept = normal(0, 1, autoscale = TRUE),
                     prior = normal(0, 1, autoscale = TRUE),
                     prior_aux= cauchy(0, 5, autoscale = TRUE),
                     offset = log(hdyrsrisk),
                     data = rbctconf, refresh=0)

#Same coding issue for follow up and post trial periods in code written by Mills et al.

summary(modelc2, digits=5, probs=c(0.025,0.5,0.975))
```


# Table 2, Model d1
```{r}
#Code given by Mills et al. (table 2b)
modeld1 <- stan_glm.nb(Incidence ~ log(hdyrsrisk) + log(Hist3yr),
                  prior_intercept=normal(0,10), prior=normal(0,10),
                  data=rbctconf,
                  diagnostic_file = file.path(tempdir(), "df.csv"),refresh=0)

summary(modeld1, digits=5, probs=c(0.025,0.5,0.975))
#This aligns with the results reported in table 2b of Mills et al,
#and model rs2b in Torgerson et al
```

# Table 2, Model e
```{r}
#Model e, table 2b Mills et al.
set.seed(150499)
modele <- stan_glm(Incidence ~ Treatment + A + B + C + D + E + F + G + H + I
                   +log(Hist3yr),
                              data = rbctconf, family = "poisson",
                              offset = log(Baseline),
                              prior = normal(0, 1),
                              prior_intercept = normal(0, 2),
                              diagnostic_file = file.path(tempdir(), "df.csv"),
                   refresh=0)

summary(modele, digits=5, probs=c(0.025,0.5,0.975))
#The results do not align with the reported results in table 2b Mills et al.

round((exp(coef(modele))[2]-1),3)*100                              # point estimate -13.7%
round((exp(posterior_interval(modele, prob = 0.95))[2,1]-1),3)*100 # lower 95% CI
round((exp(posterior_interval(modele, prob = 0.95))[2,2]-1),3)*100 # upper 95% CI

#These results with this model, give the culling effect of 13.6% reduction( CrI 0.1%,24.8%)
#This is considerable less than the effect reported in Mills et al.
#And there is less probability distribution for a culling effect
#Therefore, figure 1 in the main text is incorrect
```


## Model selection by Bayes Factor
# Original log linear poisson model Model proposed by Donnelly et al as Bayesian
```{r}
set.seed(150499)
modelrs <- stan_glm(Incidence ~ Treatment + A + B + C + D + E + F + G + H + I
                    + log(Hist3yr) +log(Baseline), poisson,
                  data = rbctconf, prior_intercept=normal(0, 10),
                  diagnostic_file = file.path(tempdir(), "df.csv"),refresh=0)

bridge_1 <- bridge_sampler(modelrs)

modelrs_hdyrsrisk <- stan_glm(Incidence ~ Treatment + A + B + C + D + E + F + G
                              + H + I + log(Hist3yr)+ log(hdyrsrisk), poisson,
                  data = rbctconf, prior_intercept=normal(0, 10),
                  diagnostic_file = file.path(tempdir(), "df.csv"),refresh=0)

bridge_2 <- bridge_sampler(modelrs_hdyrsrisk)


modeld1 <- stan_glm.nb(Incidence ~ log(hdyrsrisk) + log(Hist3yr),
                     prior_intercept=normal(0,10), prior=normal(0,10),
                     data=rbctconf,
                     diagnostic_file = file.path(tempdir(), "df.csv"),refresh=0)

bridge_3 <- bridge_sampler(modeld1)

bf(bridge_3, bridge_1)
#Estimated Bayes factor in favor of bridge_3 over bridge_1: 75962823
#Model without culling effect is supported by a large amount over the original
#log linear poisson model by Donnelly et al.

#compare using the same exposure variable
bf(bridge_3, bridge_2)
#Estimated Bayes factor in favor of bridge_3 over bridge_2: 48949011
#Model with no effect of culling is supported by a large amount over one that has.


model_d2 <- stan_glm.nb(Incidence ~ log(hdyrsrisk) + log(Hist3yr),
                      prior=normal(0, 1),
                      prior_aux = cauchy(0, 5),
                      data=rbctconf,
                      diagnostic_file = file.path(tempdir(), "df.csv"),
                      refresh=0)

bridge_4 <- bridge_sampler(model_d2)

bf( bridge_4,bridge_1)
#Estimated Bayes factor in favor of bridge_3 over bridge_1: 405.82877

bf( bridge_4,bridge_2)
#better comparison as has the same exposure variable: hdyrsrisk
#Estimated Bayes factor in favor of bridge_3 over bridge_2: 367.21850
```


\newpage
# Table 3
## Mills et al, 2024b: paper 2.
```{r}
rbctconf_outside <- read_xlsx("outside_areas_rbct.xlsx", sheet="During_Trial")

#relevel variables so correspond to Mills et al and Nature
rbctconf_outside$Triplet<-relevel(factor(rbctconf_outside$Triplet), ref="J")
rbctconf_outside$Treatment<-relevel(factor(rbctconf_outside$Treatment), ref="Survey-only")
```

## Table 3, model1: Original Poisson GLM as per Donnelly et al, Nature (2006)
<http://dx.doi.org/10.1038/nature04454>

```{r}
model1 <- glm(Incidence ~ Treatment + Triplet + log(Hist3yr)
              + log(Baseline), 
            family = poisson, data=rbctconf_outside)

summary(model1)
```

### Table 3, model1: Estimated effect of culling (95% CI)
```{r}
round((exp(coef(model1))[2] - 1),3)*100   # point estimate
round((exp(coefci(model1))[2,1]-1),3)*100 # lower 95% CI
round((exp(coefci(model1))[2,2]-1),3)*100 # upper 95% CI
```

### Table 3, model1: BIC and AICc
```{r}
round(BIC(model1),1)
round(AICc(model1),1)
```

### Table 3, model1: LOOCV RMSE 
Leave-One-Out Cross-Validation and Root Mean Squared Error, 
a metric used to evaluate the performance of a model.
```{r}
rmse <- numeric()
actual_vs_predicted <- data.frame(Actual = numeric(), Predicted = numeric())
for (i in 1:nrow(rbctconf_outside)) {
  # Exclude the ith row
  test_data <- rbctconf_outside[i, ]
  train_data <-rbctconf_outside[-i, ]
  model <- glm(Incidence ~ Treatment + Triplet + log(Hist3yr)
              + log(Baseline), family = poisson, 
                   data = train_data)
  predictions <- predict(model, newdata = test_data, type="response")
  rmse[i] <- sqrt(mean((test_data$Incidence - predictions)^2))
  actual_vs_predicted <- rbind(actual_vs_predicted, 
                               data.frame(Actual = test_data$Incidence, 
                                          Predicted = predictions))
}

rmse1<-actual_vs_predicted %>%
  mutate(rmse = sqrt((Actual - Predicted)^2))

round(mean(rmse1$rmse),2)
```

### Table 3, model1: Number of estimated parameters
```{r}
length(coef(model1))
```

### Table 3, model1: Parameter for exposure (95% CI)
```{r}
# this is on the log scale
round(coef(model1)[13],2)     # point estimate
round(coefci(model1)[13,1],2) # lower 95% CI
round(coefci(model1)[13,2],2) # upper 95% CI
```

\newpage
In the text there is also mentionned this model for other time periods.
```{r}
# Follow-up period
rbctconf_outside_follow <- read_xlsx("outside_areas_rbct.xlsx", 
                                     sheet="Follow_Up")

# relevel
rbctconf_outside_follow$Triplet<-relevel(factor(rbctconf_outside_follow$Triplet), 
                                         ref="J")
rbctconf_outside_follow$Treatment<-relevel(factor(rbctconf_outside_follow$Treatment), 
                                           ref="Survey-only")

model1_follow <- glm(Incidence ~ Treatment + Triplet + log(Hist3yr)
                     + log(Baseline), 
                   family = poisson, data=rbctconf_outside_follow)

summary(model1_follow)

confint(model1_follow)


# Post-Trial period
rbctconf_outside_after <- read_xlsx("outside_areas_rbct.xlsx", sheet ="Post_Trial")

# relevel
rbctconf_outside_after$Triplet <- relevel(factor(rbctconf_outside_after$Triplet), 
                                        ref="J")
rbctconf_outside_after$Treatment <- relevel(factor(rbctconf_outside_after$Treatment), 
                                          ref="Survey-only")

model1_after <- glm(Incidence ~ Treatment + Triplet + log(Hist3yr) 
                  + log(Baseline), family = poisson, 
                  data=rbctconf_outside_after)

summary(model1_after)
confint(model1_after)
# Plausible exposure paramter = 0.77 (CIs 0.40,1.15) 
# But no effect of culling
```

\newpage
## Table 3, Model 3: Original GLM in generalized Poisson form
```{r}
#Model 3 (=model 1 in generalized poisson form)
model3 <- glmmTMB(Incidence ~ Treatment + Triplet + log(Hist3yr) 
                  + log(Baseline), 
                family = genpois, data=rbctconf_outside)
summary(model3)
confint(model3)

#Implausible exposure parameter = 0.11 (CIs-0.16, 0.38)
exp(confint(model3))
```

### Table 3, model3: Estimated effect of culling (95% CI)
```{r}
round((exp(confint(model3, level = 0.95)[2,3])-1),3)*100 # point estimate
round((exp(confint(model3, level = 0.95)[2,1])-1),3)*100 # lower 95% CI
round((exp(confint(model3, level = 0.95)[2,2])-1),3)*100 # upper 95% CI
```


### Table 3, model1: BIC and AICc
```{r}
round(BIC(model3),1)
round(AICc(model3),1)
```

### Table 3, model3: LOOCV RMSE 
Leave-One-Out Cross-Validation and Root Mean Squared Error, 
a metric used to evaluate the performance of a model.
```{r}
rmse <- numeric()
actual_vs_predicted <- data.frame(Actual = numeric(), Predicted = numeric())
for (i in 1:nrow(rbctconf_outside)) {
  # Exclude the ith row
  test_data <- rbctconf_outside[i, ]
  train_data <-rbctconf_outside[-i, ]
  model <- glmmTMB(Incidence ~ Treatment + Triplet + log(Hist3yr) 
                  + log(Baseline), 
                family = genpois, 
                   data = train_data)
  predictions <- predict(model, newdata = test_data, type="response")
  rmse[i] <- sqrt(mean((test_data$Incidence - predictions)^2))
  actual_vs_predicted <- rbind(actual_vs_predicted, 
                               data.frame(Actual = test_data$Incidence, 
                                          Predicted = predictions))
}

rmse1<-actual_vs_predicted %>%
  mutate(rmse = sqrt((Actual - Predicted)^2))

round(mean(rmse1$rmse),2)
```

### Table 3, model3: Number of estimated parameters
```{r}
length(fixef(model3)$cond) # plus an additional parameter for overdispersion
```

### Table 3, model3: Parameter for exposure (95% CI)
```{r}
# this is on the log scale
round(confint(model3, level = 0.95)[13,3],2) # point estimate
round(confint(model3, level = 0.95)[13,1],2) # lower 95% CI
round(confint(model3, level = 0.95)[13,2],2) # upper 95% CI
```

\newpage
In the text there is also mentionned this model for other time periods.
```{r}
model3_follow <- glmmTMB(Incidence ~ Treatment + Triplet + log(Hist3yr)
                         + log(Baseline), family = genpois, 
                         data=rbctconf_outside_follow)

summary(model3_follow)

confint(model3_follow)
#Implausible exposure parameter = 0.12 (CIs-0.19, 0.12)

model3_after <- glmmTMB(Incidence ~ Treatment + Triplet + log(Hist3yr)
                        + log(Baseline), 
                      family = genpois, data=rbctconf_outside_after)
summary(model3_after)
confint(model3_after)
# Plausible exposure parameter = 0.77 (CIs 0.35, 1.19) 
# But no effect of culling
```


\newpage
## Table 3, Model 4: Generalized Poisson without culling
```{r}
model4 <- glmmTMB(Incidence ~ log(Hist3yr) + log(Baseline), 
                family = genpois, data=rbctconf_outside)
summary(model4)

confint(model4)
# Plausible exposure parameter = 0.73 (CIs 0.33, 1.14) 
# But no effect of culling
```


### Table 3, model4: BIC and AICc
```{r}
round(BIC(model4),1)
round(AICc(model4),1)

AICc(model1, model3, model4)
#Model 4 has lowest AICc compared to models which report a culling effect
```

### Table 3, model4: LOOCV RMSE 
Leave-One-Out Cross-Validation and Root Mean Squared Error, 
a metric used to evaluate the performance of a model.
```{r}
rmse <- numeric()
actual_vs_predicted <- data.frame(Actual = numeric(), Predicted = numeric())
for (i in 1:nrow(rbctconf_outside)) {
  # Exclude the ith row
  test_data <- rbctconf_outside[i, ]
  train_data <-rbctconf_outside[-i, ]
  model <- glmmTMB(Incidence ~ log(Hist3yr) + log(Baseline), 
                family = genpois, 
                   data = train_data)
  predictions <- predict(model, newdata = test_data, type="response")
  rmse[i] <- sqrt(mean((test_data$Incidence - predictions)^2))
  actual_vs_predicted <- rbind(actual_vs_predicted, 
                               data.frame(Actual = test_data$Incidence, 
                                          Predicted = predictions))
}

rmse1<-actual_vs_predicted %>%
  mutate(rmse = sqrt((Actual - Predicted)^2))

round(mean(rmse1$rmse),2)
```

### Table 3, model4: Number of estimated parameters
```{r}
length(fixef(model4)$cond) # plus an additional parameter for overdispersion
```

### Table 3, model4: Parameter for exposure (95% CI)
```{r}
# this is on the log scale
round(confint(model4, level = 0.95)[3,3],2) # point estimate
round(confint(model4, level = 0.95)[3,1],2) # lower 95% CI
round(confint(model4, level = 0.95)[3,2],2) # upper 95% CI
```

\newpage
In the text there is also mentionned this model for other time periods.
```{r}
model4_follow <- glmmTMB(Incidence ~ log(Hist3yr) + log(Baseline), 
                       family = genpois, data=rbctconf_outside_follow)
summary(model4_follow)
confint(model4_follow)
# Plausible exposure parameter = 0.71 (CIs 0.19, 1.23) 
# But no effect of culling

AICc(model1_follow, model3_follow, model4_follow)
#Model4_follow has the lowest AICc compared to models which report a culling effect


model4_after <- glmmTMB(Incidence ~ log(Hist3yr) + log(Baseline), 
                      family = genpois, data=rbctconf_outside_after)
summary(model4_after)
confint(model4_after)
# Plausible exposure parameter = 0.78 (CIs 0.45, 1.04) 
# But no effect of culling
AICc(model1_after, model3_after, model4_after)
#Model4_follow has the lowest AICc compared to models which report a culling effect
```

\newpage
## Table 3, Model 5: Generalized Poisson with herd-years-at-risk offset
```{r}
#Model 5
model5 <- glmmTMB(Incidence ~ Treatment + Triplet + log(Hist3yr) 
                  + offset(log(hdyrsrisk)), 
                family = genpois, data=rbctconf_outside)
summary(model5)
confint(model5)
exp(confint(model5))
#No effect of culling. Treatment parameter =0.09 (CIs -0.10, 0.29)
```

### Table 3, model5: Estimated effect of culling (95% CI)
```{r}
round((exp(confint(model5, level = 0.95)[2,3])-1),3)*100 # point estimate
round((exp(confint(model5, level = 0.95)[2,1])-1),3)*100 # lower 95% CI
round((exp(confint(model5, level = 0.95)[2,2])-1),3)*100 # upper 95% CI
```

### Table 3, model5: BIC and AICc
```{r}
round(BIC(model5),1)
round(AICc(model5),1)
```

### Table 3, model5: LOOCV RMSE 
Leave-One-Out Cross-Validation and Root Mean Squared Error, 
a metric used to evaluate the performance of a model.
```{r}
rmse <- numeric()
actual_vs_predicted <- data.frame(Actual = numeric(), Predicted = numeric())
for (i in 1:nrow(rbctconf_outside)) {
  # Exclude the ith row
  test_data <- rbctconf_outside[i, ]
  train_data <-rbctconf_outside[-i, ]
  model <- glmmTMB(Incidence ~ Treatment + Triplet + log(Hist3yr) 
                  + offset(log(hdyrsrisk)), 
                family = genpois, 
                   data = train_data)
  predictions <- predict(model, newdata = test_data, type="response")
  rmse[i] <- sqrt(mean((test_data$Incidence - predictions)^2))
  actual_vs_predicted <- rbind(actual_vs_predicted, 
                               data.frame(Actual = test_data$Incidence, 
                                          Predicted = predictions))
}

rmse1<-actual_vs_predicted %>%
  mutate(rmse = sqrt((Actual - Predicted)^2))

round(mean(rmse1$rmse),2)
```

### Table 3, model5: Number of estimated parameters
```{r}
length(fixef(model5)$cond) # plus an additional parameter for overdispersion
```

\newpage
In the text there is also mentionned this model for other time periods.
```{r}
model5_follow <- glmmTMB(Incidence ~ Treatment + Triplet + log(Hist3yr)
                         + offset(log(hdyrsrisk)), 
                       family = genpois, data=rbctconf_outside_follow)
summary(model5_follow)
confint(model5_follow)
#No effect of culling. Treatment parameter =0.002 (CIs -0.20, 0.21)

model5_after <- glmmTMB(Incidence ~ Treatment + Triplet + log(Hist3yr)
                        + offset(log(hdyrsrisk)), 
                      family = genpois, data=rbctconf_outside_after)
summary(model5_after)
confint(model5_after)
#No effect of culling. Treatment parameter =-0.03 (CIs -0.18, 0.12)
```

\newpage
## Table 3, Model 6: Generalized Poisson with herd-years-at risk offset and without a culling effect
```{r}
model6 <- glmmTMB(Incidence ~ log(Hist3yr) + offset(log(hdyrsrisk)), 
                family = genpois, data=rbctconf_outside)

summary(model6)
```

### Table 3, model6: BIC and AICc
```{r}
round(BIC(model6),1)
round(AICc(model6),1)

AICc(model5, model6)
#model 6, no culling effect has much lower AICc than model 5 with culling effect as an explanatory variable.
```

### Table 3, model6: LOOCV RMSE 
Leave-One-Out Cross-Validation and Root Mean Squared Error, 
a metric used to evaluate the performance of a model.
```{r}
rmse <- numeric()
actual_vs_predicted <- data.frame(Actual = numeric(), Predicted = numeric())
for (i in 1:nrow(rbctconf_outside)) {
  # Exclude the ith row
  test_data <- rbctconf_outside[i, ]
  train_data <-rbctconf_outside[-i, ]
  model <- glmmTMB(Incidence ~ log(Hist3yr) + offset(log(hdyrsrisk)), 
                family = genpois, 
                   data = train_data)
  predictions <- predict(model, newdata = test_data, type="response")
  rmse[i] <- sqrt(mean((test_data$Incidence - predictions)^2))
  actual_vs_predicted <- rbind(actual_vs_predicted, 
                               data.frame(Actual = test_data$Incidence, 
                                          Predicted = predictions))
}

rmse1<-actual_vs_predicted %>%
  mutate(rmse = sqrt((Actual - Predicted)^2))

round(mean(rmse1$rmse),2)
```

### Table 3, model6: Number of estimated parameters
```{r}
length(fixef(model6)$cond) # plus an additional parameter for overdispersion
```

\newpage
In the text there is also mentionned this model for other time periods.
```{r}
model6_follow <- glmmTMB(Incidence ~ log(Hist3yr) + offset(log(hdyrsrisk)), 
                       family = genpois, data=rbctconf_outside_follow)

#No effect of culling
AICc(model5_follow, model6_follow)
#model 6_follow, no culling effect has much lower AICc than 
#model 5_follow with culling effect as explanatory variable.

model6_after<-glmmTMB(Incidence~log(Hist3yr)+offset(log(hdyrsrisk)), 
                      family = genpois, data=rbctconf_outside_after)
#No effect of culling
AICc(model5_after, model6_after)
#model 6_follow, no culling effect has much lower AICc than model 5_follow 
#with culling effect as explanatory variable.
```

\newpage
## Table 3, Model 8: Generalized Poisson with herd-years-at-risk covariate
```{r}
#model 8. As model 6 without offset
model8 <- glmmTMB(Incidence ~ log(Hist3yr) + log(hdyrsrisk), 
                family = genpois, data=rbctconf_outside)

summary(model8)
confint(model8)
#Plausible parameter for exposure parameter 0.61 (CIs 0.39, 0.84)
```

### Table 3, model8: BIC and AICc
```{r}
round(BIC(model8),1)
round(AICc(model8),1)

AICc(model5, model6, model8)
#No effect of culling
#model 8, now has the lowest AICc

BIC(model1,model8)
#lower BIC than model 1, the original model in Nature.
```

### Table 3, model8: LOOCV RMSE 
Leave-One-Out Cross-Validation and Root Mean Squared Error, 
a metric used to evaluate the performance of a model.
```{r}
rmse <- numeric()
actual_vs_predicted <- data.frame(Actual = numeric(), Predicted = numeric())
for (i in 1:nrow(rbctconf_outside)) {
  # Exclude the ith row
  test_data <- rbctconf_outside[i, ]
  train_data <-rbctconf_outside[-i, ]
  model <- glmmTMB(Incidence ~ log(Hist3yr) + log(hdyrsrisk), 
                family = genpois, 
                   data = train_data)
  predictions <- predict(model, newdata = test_data, type="response")
  rmse[i] <- sqrt(mean((test_data$Incidence - predictions)^2))
  actual_vs_predicted <- rbind(actual_vs_predicted, 
                               data.frame(Actual = test_data$Incidence, 
                                          Predicted = predictions))
}

rmse1<-actual_vs_predicted %>%
  mutate(rmse = sqrt((Actual - Predicted)^2))

round(mean(rmse1$rmse),2)
```

### Table 3, model8: Number of estimated parameters
```{r}
length(fixef(model8)$cond) # plus an additional parameter for overdispersion
```

### Table 3, model8: Parameter for exposure (95% CI)
```{r}
#herd years at risk
# this is on the log scale
round(confint(model8, level = 0.95)[3,3],2) # point estimate
round(confint(model8, level = 0.95)[3,1],2) # lower 95% CI
round(confint(model8, level = 0.95)[3,2],2) # upper 95% CI
```

\newpage
In the text there is also mentioned this model for other time periods.
```{r}
#model 8. As model 6 without offset
model8_follow <- glmmTMB(Incidence ~ log(Hist3yr) + log(hdyrsrisk), 
                       family = genpois, data=rbctconf_outside_follow)
confint(model8_follow)
#Plausible parameter for exposure parameter 0.74 (CIs 0.51, 0.96)
#No effect of culling

AICc(model5_follow, model6_follow, model8_follow)

#model8_follow, now has the lowest AICc
BIC(model1_follow,model8_follow)
#lower BIC than model1_follow.

#model 8. As model 6 without offset
model8_after <- glmmTMB(Incidence ~ log(Hist3yr) + log(hdyrsrisk), 
                      family = genpois, data=rbctconf_outside_after)
confint(model8_after)
#Plausible parameter for exposure parameter 0.78 (CIs 0.45, 1.10)
#No effect of culling
AICc(model5_after, model6_after, model8_after)
#model6_after, still has the lowest AICc
BIC(model1_after,model8_after)
#lower BIC than model1_after.
```

\newpage
## Table 3, Model 11: Generalized poisson with no predictors
```{r}
model11 <- glmmTMB(Incidence ~ 1, family = genpois, data=rbctconf_outside)
```


### Table 3, model11: BIC and AICc
```{r}
round(BIC(model11),1)

round(AICc(model11),1)

AICc(model1, model11)
#model 11 with no predictors has lower AICc than model 1
```

### Table 3, model11: LOOCV RMSE 
Leave-One-Out Cross-Validation and Root Mean Squared Error, 
a metric used to evaluate the performance of a model.
```{r}
rmse <- numeric()
actual_vs_predicted <- data.frame(Actual = numeric(), Predicted = numeric())
for (i in 1:nrow(rbctconf_outside)) {
  # Exclude the ith row
  test_data <- rbctconf_outside[i, ]
  train_data <-rbctconf_outside[-i, ]
  model <- glmmTMB(Incidence ~ 1, 
                family = genpois, 
                   data = train_data)
  predictions <- predict(model, newdata = test_data, type="response")
  rmse[i] <- sqrt(mean((test_data$Incidence - predictions)^2))
  actual_vs_predicted <- rbind(actual_vs_predicted, 
                               data.frame(Actual = test_data$Incidence, 
                                          Predicted = predictions))
}

rmse1<-actual_vs_predicted %>%
  mutate(rmse = sqrt((Actual - Predicted)^2))

round(mean(rmse1$rmse),2)
```

### Table 3, model11: Number of estimated parameters
```{r}
length(fixef(model11)$cond) # plus an additional parameter for overdispersion
```


\newpage
In the text there is also mentionned this model for other time periods.
```{r}
model11_follow <- glmmTMB(Incidence ~ 1, family = genpois, 
                          data=rbctconf_outside_follow)

AICc(model1_follow, model11_follow)
#model 11_follow with no predictors has lower AICc than model 1_follow

model11_after <- glmmTMB(Incidence ~ 1, family = genpois, 
                         data=rbctconf_outside_after)
AICc(model1_after, model11_after)
#model 11_after with no predictors has lower AICc than model 1_after 
```

\newpage
# Bayesian Analysis Table 4
## Table 4, Model a.1: Varying intercepts for triplets and covariates of culling efefct,
historical 3-year incidence and baseline herds at risk. 
```{r}
#Code in Mills et al, purported to be the same as the code for rs in Torgerson et al:
set.seed(125347599)
model_a1 <- stan_glm.nb(Incidence ~ Treatment + A + B + C + D + E + F +G + H 
                        + I +log(Hist3yr) + log(Baseline),
                  data = rbctconf_outside, prior_intercept=normal(0, 10),
                  diagnostic_file = file.path(tempdir(), "df.csv"),refresh=0)

prior_summary(model_a1, digits = 2)

#Auxiliary (reciprocal_dispersion)
#~ exponential(rate = 1)
summary(model_a1, digits=3, probs=c(0.025,0.5,0.975))
#reciprocal_dispersion = 3.9
```

### Table 4, model_a1: Estimated effect of culling (95% CI)
```{r}
round((exp(coef(model_a1))[2]-1),3)*100 # point estimate

round((exp(posterior_interval(model_a1, prob = 0.95))[2,1]-1),3)*100
round((exp(posterior_interval(model_a1, prob = 0.95))[2,2]-1),3)*100
```

### Table 4, model_a1: LOO
```{r, message = FALSE}
loo(model_a1, k_threshold = 0.7)
```

\newpage
## Table 4, model rs: Actual code for model rs in Torgerson et al.
```{r}
set.seed(93846)
rs <- stan_glm(Incidence~Treatment+A+B+C+D+E+F+G+H+I+log(Hist3yr)+log(Baseline),
                        data = rbctconf_outside, prior_intercept=normal(0, 10), poisson,
                        diagnostic_file = file.path(tempdir(), "df.csv"),refresh=0)

#This is effectively model 1, table 3 in Bayesian form. model_a1 forced
#overdispersion by using negative binomial regression and forcing a low reciprocal dispersion parameter

summary(rs, digits=3, probs=c(0.025,0.5,0.975))
```

### Table 4, model rs: Estimated effect of culling (95% CI)
```{r}
round((exp(coef(rs))[2]-1),3)*100    # point estimate                        
round((exp(posterior_interval(rs, prob = 0.95))[2,1]-1),3)*100 # lower 95% CI
round((exp(posterior_interval(rs, prob = 0.95))[2,2]-1),3)*100 # upper 95% CI
```

### Table 4, model rs: LOO
```{r, message = FALSE}
loo(rs, k_threshold = 0.7)
```

\newpage
## Table 4, model_a2
```{r}
set.seed(6754)
model_a2 <- stan_glm.nb(Incidence~Treatment+A+B+C+D+E+F+G+H+I+log(Hist3yr)+log(Baseline),
                           data = rbctconf_outside, prior_intercept=normal(0, 1, autoscale=TRUE),
                           prior_aux= cauchy(0, 5),
                           prior = normal(0, 1),
                           diagnostic_file = file.path(tempdir(), "df.csv"),refresh=0)

prior_summary(model_a2, digits = 2)

#Auxiliary (reciprocal_dispersion)
#~ half-cauchy(location = 0, scale = 5)

summary(model_a2, digits=3, probs=c(0.025,0.5,0.975))
#reciprocal_dispersion = 194.1
# Note as reciprocal dispersion increases in trends towards a poisson model. 
#When reciprocal_dispersion = infinity, it is poisson
#With such a high reciprocal dispersion, this is close to poisson, ie Model 1 in Bayesian form.
```

### Table 4, model_a2: Estimated effect of culling (95% CI)
```{r}
round((exp(coef(model_a2))[2]-1),3)*100    # point estimate                        
round((exp(posterior_interval(model_a2, prob = 0.95))[2,1]-1),3)*100 # lower 95% CI
round((exp(posterior_interval(model_a2, prob = 0.95))[2,2]-1),3)*100 # upper 95% CI
```

### Table 4, model_a2: LOO
```{r, message = FALSE}
loo(model_a2)
```

\newpage
## Table 4, Model b1
```{r}
set.seed(176254)
model_b1 <- stan_glm.nb(Incidence~Treatment+log(Hist3yr)+log(Baseline),
                        data = rbctconf_outside, prior_intercept=normal(0, 10),
                        diagnostic_file = file.path(tempdir(), "df.csv"),refresh=0)
summary(model_b1, digits=3, probs=c(0.025,0.5,0.975))
```

### Table 4, model_b1: Estimated effect of culling (95% CI)
```{r}
round((exp(coef(model_b1))[2]-1),3)*100                              # point estimate -13.7%
round((exp(posterior_interval(model_b1, prob = 0.95))[2,1]-1),3)*100 # lower 95% CI
round((exp(posterior_interval(model_b1, prob = 0.95))[2,2]-1),3)*100 # upper 95% CI
```

### Table 4, model_b1: LOO
```{r, message = FALSE}
loo(model_b1)
```

\newpage
## Table 4, Model b2
```{r}
set.seed(8745)
model_b2 <- stan_glm.nb(Incidence ~ Treatment + log(Hist3yr) + log(Baseline),
                        data = rbctconf_outside, prior_intercept=normal(0, 1, autoscale=TRUE),
                        prior_aux= cauchy(0, 5),
                        prior = normal(0, 1),
                        diagnostic_file = file.path(tempdir(), "df.csv"),refresh=0)

summary(model_b2, digits=3, probs=c(0.025,0.5,0.975))
# Parameter values give increase in incidence of 23.6% (CrI -12%, 174%)
#This does not align with values given in Mills et al 2024b
```

### Table 4, model_b2: Estimated effect of culling (95% CI)
```{r}
round((exp(coef(model_b2))[2]-1),3)*100                              # point estimate 
round((exp(posterior_interval(model_b2, prob = 0.95))[2,1]-1),3)*100 # lower 95% CI
round((exp(posterior_interval(model_b2, prob = 0.95))[2,2]-1),3)*100 # upper 95% CI

```

### Table 4, model_b2: LOO
```{r}
loo(model_b2)
```

\newpage
## Table 4, Model c.1: Using herd-years-at-risk as an offset and an explanatory variable and no varying intercepts for triplets
```{r}
#Code given by Mills et al: (asrs1_outside_aB)
set.seed(7264)
model_c1 <- stan_glm.nb(Incidence ~ Treatment + log(hdyrsrisk) + log(Hist3yr),
                            offset = log(hdyrsrisk), 
                            prior_intercept=normal(0,10), prior=normal(0,10),
                            data = rbctconf_outside, refresh=0)
#offset appears as both an explanatory variable and offset
#software ignores the offset
#It is a non offset model
```

### Table 4, model_c1: Estimated effect of culling (95% CI)
```{r}
round((exp(coef(model_c1))[2]-1),3)*100                              # point estimate 
round((exp(posterior_interval(model_c1, prob = 0.95))[2,1]-1),3)*100 # lower 95% CI
round((exp(posterior_interval(model_c1, prob = 0.95))[2,2]-1),3)*100 # upper 95% CI
```

### Table 4, model_c1: LOO
```{r, message = FALSE}
loo(model_c1, k_threshold = 0.7)
```


### modelc_1a: in comparison to model_c1, without the offset coded
```{r}
set.seed(9836)
model_c1a<-stan_glm.nb(Incidence~Treatment+log(hdyrsrisk)+log(Hist3yr), 
                      prior_intercept=normal(0,10), prior=normal(0,10),
                      data = rbctconf_outside, refresh=0)
summary(model_c1)
summary(model_c1a)
#identical results but with log(hrdyrsrisk) parameter translated by 1.
```

### Correct code for model_c1 should be
```{r}
set.seed(6472)
model_c1_offset<-stan_glm.nb(Incidence~Treatment+log(Hist3yr),
                      offset = log(hdyrsrisk), 
                      prior_intercept=normal(0,10), prior=normal(0,10),
                      data = rbctconf_outside, refresh=0)
summary(model_c1_offset, digits=3, probs=c(0.025,0.5,0.975))
```

### Table 4, model_c1_offset: Estimated effect of culling (95% CI)
```{r}
round((exp(coef(model_c1_offset))[2]-1),3)*100                     # point estimate 
round((exp(posterior_interval(model_c1_offset, prob = 0.95))[2,1]-1),3)*100 # lower 95% CI
round((exp(posterior_interval(model_c1_offset, prob = 0.95))[2,2]-1),3)*100 # upper 95% CI
```

### Table 4, model_c1: LOO
```{r, message = FALSE}
loo(model_c1_offset)
```

\newpage
## Table 4, Model c2 without offset
Likewise the code for model c2 has no offset for the same reasons.
Code for model_c2 given by mills et al: (model rs_1_outside_aB_improved)
```{r}
model_c2 <- stan_glm.nb(Incidence ~ Treatment + log(hdyrsrisk) + log(Hist3yr),
                                     prior_intercept = normal(0, 5),
                                     prior = normal(0, 1),
                                     prior_aux= cauchy(0, 5),
                                     offset = log(hdyrsrisk),
                                     data = rbctconf_outside, refresh=0)

#This is not an offset model.
```

\newpage
## Table 4, Model d1, no culling effect from table 4 (=model rs2B)
```{r}
set.seed(85673)
model_d1 <- stan_glm.nb(Incidence ~ log(hdyrsrisk) + log(Hist3yr),
                          prior_intercept=normal(0,10), prior=normal(0,10),
                          data=rbctconf_outside,
                      diagnostic_file = file.path(tempdir(),"df.csv"),refresh=0)

summary(model_d1, digits=3, probs=c(0.025,0.5,0.975))
```


## Table 4, Model_d1 improved
```{r}
model_d1i <- stan_glm.nb(Incidence ~ log(hdyrsrisk) + log(Hist3yr),
                                   prior=normal(0, 1),
                                   prior_aux = cauchy(0, 5),
                                   data=rbctconf_outside,
                       diagnostic_file = file.path(tempdir(), "df.csv"),
                       refresh=0)

summary(model_d1i, digits=3, probs=c(0.025,0.5,0.975))

```

\newpage
## Table 4, model e, as coded in Mills et al. 2024b
```{r}
set.seed(8489)
model_e <- stan_glm(Incidence~Treatment+A+B+C+D+E+F+G+H+I+log(Hist3yr)+log(Baseline),
                        data = rbctconf_outside, 
                        offset=log(Baseline),
                        prior_intercept=normal(0, 2),
                        poisson,
                        prior = normal(0, 1),
                        diagnostic_file = file.path(tempdir(), "df.csv"),refresh=0)

#Again this codes the offset as both an explanatory variable and offset variable.
#This is effectively the original glm in Donnelly et al, but respecified in the 
#Bayesian paradign
#All figures in the main text that relies on this model_e, 
#assuming the offset are invalid.
#Also the effect size does not align with the reported effect size in Table 2b 
#in Mills et al where it is reported.

#The same issues are also present in the coding and analysis for follow up and after.

```

### Table 4, model e: Estimated effect of culling (95% CI)
```{r}
round((exp(coef(model_e))[2]-1),3)*100                     # point estimate 
round((exp(posterior_interval(model_e, prob = 0.95))[2,1]-1),3)*100 # lower 95% CI
round((exp(posterior_interval(model_e, prob = 0.95))[2,2]-1),3)*100 # upper 95% CI
```

### Table 4, model e: LOO
```{r, message = FALSE}
loo(model_e, k_threshold = 0.7)
```

## Table 4, model e correctly specified
```{r}
set.seed(18489)
model_e.c <- stan_glm(Incidence ~ Treatment + A + B + C + D + E + F + G + H 
                      + I +log(Hist3yr),
                        data = rbctconf_outside, 
                        offset=log(Baseline),
                        prior_intercept=normal(0, 2),
                        poisson,
                        prior = normal(0, 1),
                        diagnostic_file = file.path(tempdir(), "df.csv"),refresh=0)

summary(model_e.c, digits=5, probs=c(0.025,0.5,0.975))
```

### Table 4, model e.c: Estimated effect of culling (95% CI)
```{r}
round((exp(coef(model_e.c))[2]-1),3)*100                     # point estimate 
round((exp(posterior_interval(model_e.c, prob = 0.95))[2,1]-1),3)*100 # lower 95% CI
round((exp(posterior_interval(model_e.c, prob = 0.95))[2,2]-1),3)*100 # upper 95% CI
```

### Table 4, model e.c: LOO
```{r, message = FALSE}
loo(model_e.c, k_threshold = 0.7)
```

### Model 11. as coded in Mills et al. 2024b
```{r}
model11 <-glmmTMB(Incidence~offset(log(hdyrsrisk)),genpois,  data =rbctconf_outside)

set.seed(94060000)
pp_check(model11) #Figure 3 supplementary material 3.
```
